# PVT1 splicing activity predicts genome-wide gene expression with miRNA regulatory signatures

**DOI:** 10.1101/2025.07.25.666741

**Authors:** Angelos Kozonakis, Christos Katsioulas, Evgenia Ntini

**Affiliations:** Institute of Molecular Biology and Biotechnology, IMBB-FORTH; University of Crete

## Abstract

Long non-coding RNAs (lncRNAs) have emerged as key regulators of gene expression, yet the functional impact of their splicing activity remains poorly understood. Here, we focus on the chromatin-associated lncRNA PVT1, whose splicing efficiency at specific 3’ splice sites predicts expression of distal genes in breast cancer. Using elastic net modeling across hundreds of breast cancer samples, we identify PVT1 splicing activity as a predictor of gene expression signatures enriched in miR-200/205 targets. Causal inference analysis using tumor-specific SNVs confirms that perturbation of splicing at specific sites alters the expression of distal gene sets enriched in miR-200/205 targets. We suggest a mechanistic model in which inefficient splicing and intron retention expose intronic miR-200 seed sites, perfect 7-mer’s, found exclusively within PVT1 introns, enabling PVT1 to function as a chromatin-associated lncRNA with intron-retained competing endogenous (ceRNA) activity. Artificial splicing enhancement of PVT1 in *vitro* with CASFx altered the expression of miR-200 target genes supporting this model. A genome-wide screen for intronic miR-200 seed sites, and splicing-based modeling across additional lncRNAs and protein-coding genes, identified PVT1 as a top candidate for splicing-modulated intron-retained ceRNA activity. Cross-tissue model generalization to prostate cancer supports the broader relevance of PVT1 splicing-based regulation. Our results highlight the regulatory potential of lncRNA splicing activity in *trans*, suggesting that chromatin-tethered, intron-retained lncRNAs such as PVT1 may serve as key post-transcriptional regulators in cancer.

## INTRODUCTION

An increasing body of data underpins RNA as a basic component of dynamic subnuclear compartments that contribute to global regulation of gene expression, providing a rather RNA-centric view of transcription regulation and genome organization [1–3]. RNA processing is a complex and composite mechanism with interlinked steps involving spatial and temporal dimensions, and the crosstalk between transcription, splicing dynamics and chromatin dissociation rates of nascent RNA is undergoing characterization [4–7]. Understanding the dynamics of RNA localization and interactions with proximal and/or distal genomic loci, and subnuclear compartments (i.e. transcriptional and splicing multifunctional hubs, like speckles [8]), is a major focus of research.

The role of long non-coding (lnc) RNAs has been characterized in several contexts (reviewed in [9–11]), with recent studies highlighting the lack of single RNA-driven target gene regulation [12]. Plausibly, enhancer-transcribed lncRNAs (i.e. lncRNAs transcribed from enhancer regions and the anchor points of chromosomal loops) exert regulatory potential in *cis* upon their release from chromatin, and within pre-established chromosomal proximity [13, 14]. Proximity to genomic and transcriptional sites emerges as a key factor shaping RNA-chromatin interactions, with recent evidence underpinning proximity-dominated RNA-chromatin interactions [12]. Furthermore, the role of splicing as a co- or post-transcriptional process is being intensively studied as a factor dictating or regulating RNA localization dynamics [15–20]. By means of expansion microscopy (and live-cell imaging) a transcription-proximal zone where slowly chromatin-released RNA transcripts may undergo post-transcriptional splicing, was identified [5]. What governs the timing of splicing relative to transcription termination and chromatin dissociation of the nascent RNA transcript across different cell states and genes, and if there are specific signaling pathways or nuclear factors that regulate whether splicing occurs co-transcriptionally or post-transcriptionally (in a transcription site-‘proximal zone’), are still outstanding questions. The findings from these studies hold implications for counteracting diseases where RNA processing (splicing) and RNA-chromatin interactions are disrupted, such as cancer or neurodegeneration [21].

By combining pulse-chase metabolic labeling with chromatin fractionation, we profiled the chromatin dissociation dynamics of nascent RNA transcripts and built machine-learning models integrating variables associated with either fast or slow chromatin-release kinetics of lncRNAs and mRNAs [7]. In that work, we found PVT1 among top lncRNA candidates with very slow chromatin dissociation dynamics, essentially appearing as a chromatin-associated or chromatin-retained lncRNA in MCF-7 cells at steady state (see also Suppl. Fig. S1A). By means of FISH, PVT1 was detected accumulating near its transcription locus in different cell lines (HeLa [22] and HEK293 [23]). By profiling RNA-chromatin interactions globally with CHAR-seq, Limouse et al. also uncovered PVT1 as an RNA localized predominantly near its transcription locus [12]. Thus, PVT1 is an example of chromatin-associated RNAs (caRNAs) with specific localization characteristics, namely a low delocalization score. In that study, PVT1 was identified as a lncRNA with a potential role in interacting with chromatin to regulate gene expression, and like other lncRNAs, PVT1 may have regulatory functions that rely on proximity-driven interactions with chromatin.

Most lncRNAs are thought to remain retained at chromatin, appearing as chromatin-tethered or chromatin-associated at steady state, or localized near their transcription site affecting local gene expression [24]. Accumulation of lncRNAs in the chromatin fraction has been reported in several other studies [25–27], potentially modulating gene expression locally by accumulation at or near their site of transcription [9, 28]. Although lncRNAs in general are considered potential chromatin regulatory RNAs, the recent data from Limouse et al. indicate that lncRNAs (exons) constitute only ∼3% of the total caRNA population and less than 1% when excluding the top 10 most abundant lncRNAs [12]. lncRNAs had a wide range of delocalization scores, with a score distribution that mirrored those of mRNAs. In concordance, by profiling chromatin dissociation dynamics of nascent RNA transcripts, we found that lncRNAs exhibited a wide range of chromatin dissociation rates, ranging from the slow chromatin-released (or chromatin-retained) lncRNAs, like PVT1, to the fast, efficiently chromatin-dissociated lncRNAs, that included enhancer-transcribed lncRNAs (elncRNAs), like A-ROD [7]. PVT1 as a chromatin-enriched lncRNA was also found to stay resident near its transcription locus (at the site of its production) in HEK293 cells [23] and in HeLa cells [22] (i.e. smFISH detected accumulation of the lncRNA transcript at the gene locus).

Apart from the implications in *vitro*, few studies have shed light on the role of PVT1 in *vivo*, albeit less in humans. In mouse models, a p53-dependent PVT1 isoform induced by DNA damage accumulates near its transcription locus and represses MYC transcription in *cis* [29]. The PVT1 gene frequently occurs in tandem with MYC amplification in patient tumors [30], and is overexpressed in several types of cancer. Potentially through its interaction with MYC, it may be promoting progression, invasion and chemo/radiotherapy resistance in different types of tumor. How PVT1 transcriptional activity and its RNA processing impacts MYC regulation and tumor progression has been studied before: silencing PVT1 promoter enhances breast cancer cell competition in *vivo*, while PVT1 and MYC promoters compete for enhancer contacts in *cis* [31]. As a chromatin-associated lncRNA, with very slow chromatin dissociation dynamics, PVT1 is expected to undergo extensive alternative splicing, potentially due to low splicing efficiency at individual splice sites and stochastic splice site selection. However, to what extent any alternative co- or post-transcriptional splicing activity at or near the PVT1 locus impacts global or local gene expression, and/or chromatin structure and function, and what is the role of alternative PVT1 splicing isoforms in different tumors and/or tumor subtypes, and tumor development and progression, have yet to be elucidated.

Furthermore, besides the proximity-favored regulatory crosstalk between PVT1 and MYC [29, 31], the impact of PVT1 splicing dynamics and alternative splicing activity at the PVT1 locus on a broader gene expression regulation remains to be determined.

Functional validation of specific lncRNAs, like PVT1, in chromatin regulation and regulation of gene expression remains challenging. In this study, we framed PVT1 as an example of a lncRNA that may function at the interface of transcriptional networks, and whose RNA processing may contribute to complex regulatory networks controlling gene expression. This complexity is underscored by changes in splicing activity at the PVT1 locus and by the dynamic interactions of the PVT1 lncRNA with chromatin, which may change during cell differentiation, tumor development and progression, and/or in response to stress or DNA damage.

We aim to address specific questions regarding PVT1 and similar chromatin-associated lncRNAs, including the specific mechanisms by which PVT1 interacts with chromatin and affects gene expression, and what is the role of stimulated changes in its splicing activity in regulatory function. Furthermore, at least two outstanding questions remain to be resolved 1. What is the mechanistic basis of lncRNA accumulation at or near transcription sites? (i.e. is it mediated by specific RNA binding proteins? chromatin structure or structural elements, either at chromatin or RNA level, and/or transcription or splicing dynamics?) 2. What governs a transition from co- to post-transcriptional RNA splicing mode, i.e. what determines whether splicing occurs co- or post-transcriptionally, and how does this influence lncRNA functional potential?

We focused on PVT1 as a lncRNA whose co- and/or post-transcriptional splicing dynamics may affect regulation of gene expression, in *cis* and/or at distal sites. To uncover a role of PVT1 splicing activity in shaping genome-wide regulation of gene expression, we leveraged RNA-sequencing data from The Cancer Genome Atlas (TCGA). We extracted splicing efficiency at all PVT1 acceptor 3’ splice sites from hundreds of normal and cancer samples in several types of tumors where PVT1 is expressed, with a primary focus on breast cancer (BRCA). Using the extracted splicing efficiency values, we built machine learning models, linear regression and regularized linear regression (elastic nets), to predict gene expression genome-wide, through genome-wide in *silico* screens (Figure 1). Clustering gene expression based on splice site regression coefficients revealed that distinct PVT1 splice sites affect different sets of genes. By analysis of alternative splicing, we uncovered the potential of distinct PVT1 splicing isoforms in driving tumor progression and distinguishing between different breast cancer tumor subtypes. Mechanistically, PVT1 contains multiple seed sites for miR-200/205 located exclusively within its intronic regions. Notably, miR-200/205 target genes are significantly enriched among the sets of genes predicted by the PVT1 splicing-based machine learning models. To validate this model, that a PVT1 regulatory impact on distal target gene expression, —through its splicing dynamics, which is deregulated in cancer—, can be mediated via an intron-retained competing endogenous (ceRNA) activity, we leveraged TCGA-annotated tumor-specific mutations (single nucleotide variants, SNVs) clustered near PVT1 splice sites. Causal inference analysis indicates that perturbation of PVT1 splicing at certain splice sites influences target gene expression, uncovering a link between splicing dynamics of a chromatin-associated lncRNA and regulation of gene expression in *trans*.

**Figure 1.**
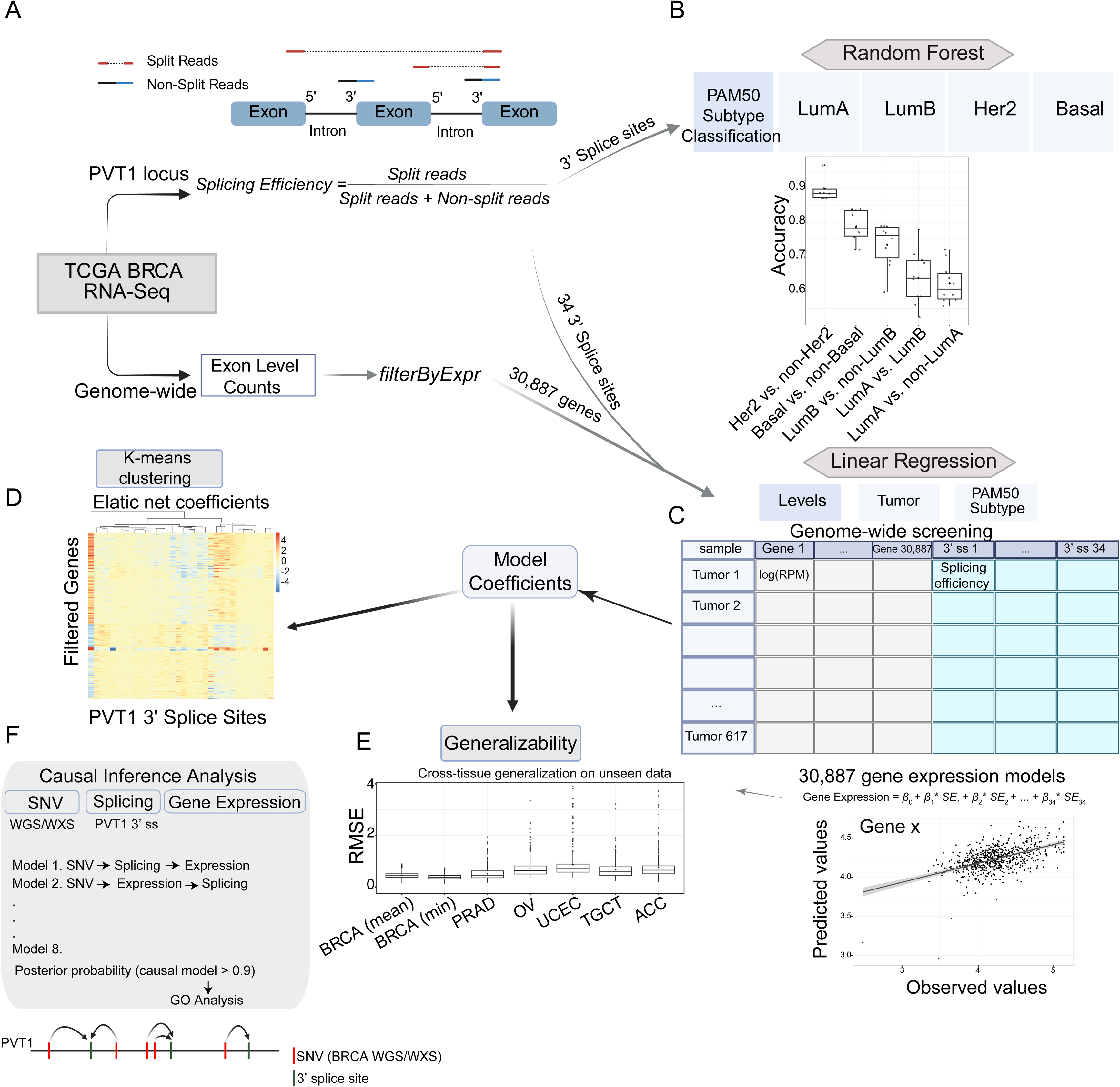
Schematic representation of the workflow used in the study. (A) The initial cohort was built on hundreds of de-identified breast cancer (BRCA) samples for which RNA-seq data was downloaded from The Cancer Genome Atlas (TCGA) and used to extract i) splicing efficiency values at the PVT1 locus and ii) genome-wide gene expression using exon level read counts. Splicing efficiency was calculated as the fraction of split reads to the sum of split plus non-split reads at the 3’ splice sites of PVT1 (top schematic). (B) A total of 34 unique PVT1 3’ splice sites were used in classification analysis of TCGA-BRCA tumor samples. Samples were subtyped with the PAM50 signature (LumA, LumB, Her2, Basal) and Random Forest with 10 x cross-validation was applied using the extracted splicing efficiencies as features in binary classifications (specific subtype versus rest samples). (C) To assess whether splicing activity at the PVT1 locus can be predictive of gene expression, the same set of 34 3’ splice site splicing efficiencies were employed as predictor variables in linear regression models with 10 x cross-validation (linear regression, lm and regularized linear regression with elastic net, glmnet). Genome-wide screening using elastic net models (30,887 genes) revealed sets of genes predicted at high-confidence (based on specific significance cutoffs) across all-tumor samples and within PAM50 subtyped cohorts. (D) In order to uncover regulatory relationships between splicing activity at PVT1 3’ splice sites and their predicted target genes, k-means clustering of high-confidence predicted genes was performed using the regression coefficients from the splicing-based elastic net models. Clustering at both all-tumor and PAM50 subtype levels identified distinct sets of splice sites predicting different sets of genes. (E) Cross-tissue generalization of BRCA-trained PVT1 splicing-based gene expression models was tested on unseen data from TCGA Prostate (PRAD), Ovary (OV), Uterus (UCEC), Testis (TGCT) and Adrenal gland (ACC). (F) To uncover causal relationships between splicing activity of PVT1 and regulation of gene expression in *trans*, we leveraged BRCA whole genome (WGS) and exon (WXS) SNV data. Tumor-specific somatic SNVs near PVT1 splice sites were assigned to their closest splice site. Upon identification of samples with SNV-associated perturbation in splice-site splicing efficiency, causal inference analysis was performed to identify gene expression changes best explained by the causal model (SNV → Splicing → Expression). Posterior probabilities were calculated for eight alternative Bayesian network models.

## RESULTS & DISCUSSION

### Splicing activity of PVT1 lncRNA is predictive of gene expression with an enrichment for miR-200/205 target genes

While the association between lncRNA expression levels and tumor prognosis or progression has been studied, the connection between differential lncRNA splicing activity and its mechanistic role in disease development remains unclear in various contexts [32, 33]. PVT1 is a complex locus with several alternative splicing isoforms annotated (> 30 in GENCODE basic V44 or V48; Suppl. Fig. S1A-B, Figure 2). Still, it remains an open question whether splicing of lncRNAs in general, and of PVT1 in frame, has any biological relevance. For lncRNAs transcribed from enhancers, splicing activity has been associated with increased enhancer activity and regulation of target gene expression in *cis,* or within pre-established proximity of chromosomal loops [7, 9, 14, 34, 35]. Through nascent RNA transcriptomics and machine-learning predictive modeling, we showed that it is more likely the overall splicing activity at a lncRNA locus, rather than the splicing efficiency at certain donor or acceptor splice sites per *se*, that relates with chromatin dissociation dynamics of nascent RNA transcript, and functional potential of chromatin-released lncRNAs [7]. These models remain to be experimentally validated through splicing modulation of lncRNAs, which is now possible by using CRISPR-based artificial splicing systems (CASFx [36, 37]).

**Figure 2.**
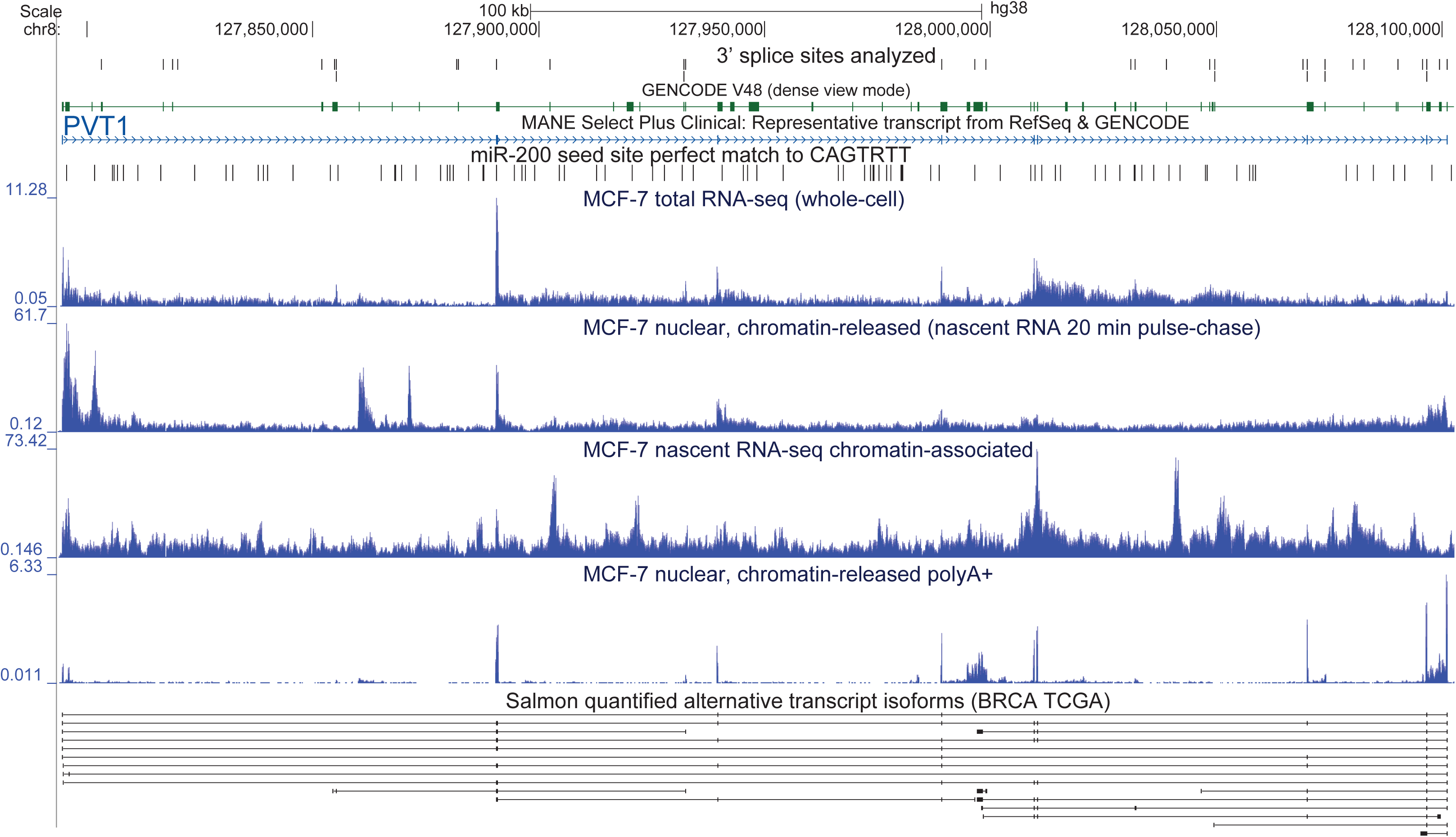
UCSC screenshot of the PVT1 locus (hg38). Tracks include coordinate positions of the 3’ splice sites analyzed, their splicing efficiency extracted from TCGA BRCA RNA-seq. miR-200 seed sites, perfect matches to the 7-mer CAGTRUU, are found exclusively within PVT1 intronic intervals. PVT1 expression is shown with plus-strand-specific bigwig (reads per million of uniquely mapped read at nucleotide position) in MCF-7 whole-cell total RNA-seq (from [90]), in nascent RNA from the nuclear chromatin-released RNA fraction upon 20 min 4-SU pulse-chase [7], in nascent chromatin-associated RNA (0 min pulse-chase from [7]), and in steady-state nuclear chromatin-associated polyA+ enriched RNA (this study, GEO-submitted). Lower track shows alternative PVT1 transcript isoforms identified and quantified with Salmon [57] using TCGA BRCA RNA-seq; only the top 19 transcripts out of 190 identified are shown, the most variable in expression across all BRCA tumor samples analyzed (see also Suppl. Figure S7B-C).

Like most lncRNAs, PVT1 is expressed in a cell type-specific manner, and in normal tissue it is highly expressed in breast, ovary and adrenal gland (Suppl. Fig. S1C, GTEx [38]). In MCF-7 (ER+) breast cancer cells, as well as in other cancer cell lines, PVT1 appears as a chromatin-tethered or chromatin-associated lncRNA at steady state; it accumulates near its site of transcription, shown either by measuring its chromatin dissociation dynamics with pulse-chase nascent RNA-sequencing [7] (Suppl. Fig. S1A), or by FISH [22]. lncRNA expression is dysregulated in various cancers, however the roles of lncRNAs in tumor development and progression remain largely unknown, and only few case studies have shed light on specific examples (e.g. the role of lncRNA SNHG6 in hepatocellular carcinoma progression [39]). In some cases, lncRNAs may stabilize proteins via RNA-protein interactions (e.g. PVT1 stabilizes Lin28 promoting colorectal cancer [40]), or may function as ‘competing-endogenous’ RNA (ceRNA) sequestering miRNAs (e.g. SNHG6 sponging miR-26a/b [39]). PVT1 expression is upregulated in several cancers including gastric, liver, lung, esophageal, colorectal ([41] and references therein). Upregulated lncRNAs in several types of cancer can be useful as early diagnostic markers and/or putative therapeutic targets. For instance, PVT1 enhances MYC protein levels, and since MYC protein is not druggable by small molecules, targeting its dependence on PVT1 lncRNA could provide a promising therapeutic intervention [30]. Due to its upregulation in several types of cancer, PVT1 is considered an oncogene in various tumors, however the exact molecular mechanisms and downstream effector pathways of PVT1 remain to be characterized. Identification of the upstream and downstream targets of PVT1 and the molecular mechanisms underlying its oncogenic character, and characterizing the functions of its individual alternative transcripts, could help uncover its role in tumorigenesis.

PVT1 was previously reported to act as a putative ‘competing endogenous’ lncRNA (ceRNA) sequestering (sponging) members of the miR-128 family [41] and miR-200 family [42, 43]. Indeed, each of three PVT1 exons bear one 6-mer seed site either for miR-200a/miR-141 (AGU<u>G</u>UU) or miR-200b/200c/429 (AGU<u>A</u>UU). However, several 7-mer miR-200 consensus seed sites (CAGURUU), and 10 perfect 8-mer seed sites (CAGURUUA) are found exclusively within PVT1 intronic sequences (Suppl. Figure S1A-B; miR-200 seed sites and putative target genes were extracted from [44–50] and cross-validated at miRPath DataBase (v 2.0) [51]. This finding points at the possibility that differential PVT1 splicing activity may be involved in regulation of gene expression post-transcriptionally, via a ceRNA – miR-200 regulatory axis. PVT1 splicing deregulated in cancer may lead to increased intron retention and abnormal (elevated) sponging effect leading to deregulation of miR-200 target gene expression. In line with this hypothesis, Gregory et al. showed that all five members of the miR-200 family are downregulated in cancer cells that have undergone epithelial to mesenchymal transition (EMT), followed by upregulation of ZEB1/ZEB2 miR-200 target genes, underlying a crucial step in tumor progression (marking the onset of metastasis) [46]. Apart from miR-200 family members deregulated in cancer, miR-205 target genes are also involved in regulating EMT [46]; of note, miR-205 seed sites (AUGAAGG for -5p and CUGAAAU for -3p) are also found within intronic PVT1 sequences (Suppl. Fig S1E).

To uncover the role and biological relevance of PVT1 splicing dynamics in gene regulation, in a tumorigenic context, we leveraged breast cancer TCGA RNA-seq data pre-applying specific filters (Methods), ending up with 696 breast cancer samples and 80 matched control samples. To account for alternative splicing activity at the PVT1 locus, we focused on splicing at the 3’ (acceptor) splice sites, treating it as the determinant step that defines the final splicing products [15, 19]. We extracted the ratio of split to non-split reads at the 3’ splice sites of PVT1, focusing on 34 unique 3’ splice sites that passed specific filtering criteria, namely a certain read coverage cutoff and measured splicing efficiency value in most samples analyzed (details outlined in Methods).

We used the PVT1 splicing efficiency values to train machine learning models, both linear regression with 10 x cross-validation, and regularized linear models via penalized maximum likelihood using elastic nets (i.e. regression for continuous outcomes implemented with glmnet [52]), to predict gene expression genome-wide, through genome-wide in *silico* screens (Figure 1 Pipeline). PVT1 expression is in general upregulated in cancer, confirmed also in TCGA data (Suppl. Fig. S1D). By comparing splicing efficiency values at 3’ splice sites between control and tumor condition we found that certain sites show lower splicing efficiency in the tumor (Suppl. Fig. S2A-C). Based on that, we extracted variance in splicing efficiency across all BRCA samples and set an arbitrary cutoff to select the top 12 most variable 3’ splice sites (Suppl. Fig. S2D). We noted fewer significant differences in 5’ donor splicing efficiency between control and tumor, and different variance in splicing efficiency at PVT1 5’ splice sites across BRCA samples (Suppl. Fig. S2B-E). By using the 12 most variable PVT1 3’ splice sites in simple linear regression models to predict gene expression genome-wide, we found 16 genes that pass an R^2 cutoff of 0.2 (Suppl. Fig. S2F). By including more splice sites in the model, performance improved with more genes passing the cutoff (Suppl. Fig. S2F), thus eventually we included all 34 3’ splice sites in linear regression model with 10 x cross validation, which identified 316 genes passing a cutoff of performance (R-squared from the best fold > 0.2 and mean correlation from 10 x cross-validation > 0.2) (Suppl. Fig. S2G-H). Among those 316 genes, miR-200 target genes are significantly enriched (Fisher’s exact test 4.572e-16, odds ratio ∼3.66). Gene ontology analysis and terms enrichment (using clusterProfiler [53]) suggests genes involved in mitochondrial oxidative phosphorylation and redox processes (energy metabolism), RNA splicing (RNA processing) and translation (Suppl. Fig. S2I).

Expression of PVT1 itself has no predictive capacity in linear regression model (only 53 predicted genes pass the performance cutoff with no miR-200 target gene enrichment; Suppl. Fig. S2J), aligning with recent findings that expression and splicing may underlie distinct biological signals [54]. Moreover, neither PVT1 nor MYC expression are efficiently predicted by PVT1 splicing (Suppl. Fig. S2J), suggesting putative roles of PVT1 splicing activity in regulating gene expression post-transcriptionally, in *trans*. To account for correlations between 3’ splice site splicing efficiencies (Suppl. Fig. S3A) we applied elastic nets (with 10 x cross-validation, Methods) to identify subsets of splicing events (among the 34 3’ splice sites) that are the most informative in predicting target gene expression. High correlation in splicing efficiency values between splice sites could indicate concurrent splicing, and/or splice sites processed co-transcriptionally in the same transcript (Suppl. Fig. S1A, S2A, S3A).

Elastic nets offer the advantage of combining two methods of regularization, the lasso (L1) and Ridge (L2). Lasso regularization encourages sparsity by shrinking some (non-important) coefficients to zero, helping feature selection, while the Ridge method shrinks coefficients toward zero, while handling multicollinearity among features. Thus, elastic net combining both methods, balances sparsity and stability of feature selection [55]. We found 365 genes passing the average performance cutoffs (from 10 x cross-validation), with miR-200 target genes significantly enriched among them (Fisher’s exact test p-value 1.51e-14, odds ratio 3.12) (Figure 3A-C). Gene ontology (GO) term enrichment analysis suggests terms skewed toward ribosomal and translation-associated components (Figure 3C). Between the two models, linear regression and elastic net, we found 100 common genes predicted by both models at the pre-defined performance cutoffs (Methods), and 20 of those are miR-200 target genes. Together these results indicate that PVT1 splicing at 3’ splice sites is predictive of distal gene expression, potentially —and at least in part— through a mechanism that involves miR-200/205 ceRNA activity.

**Figure 3.**
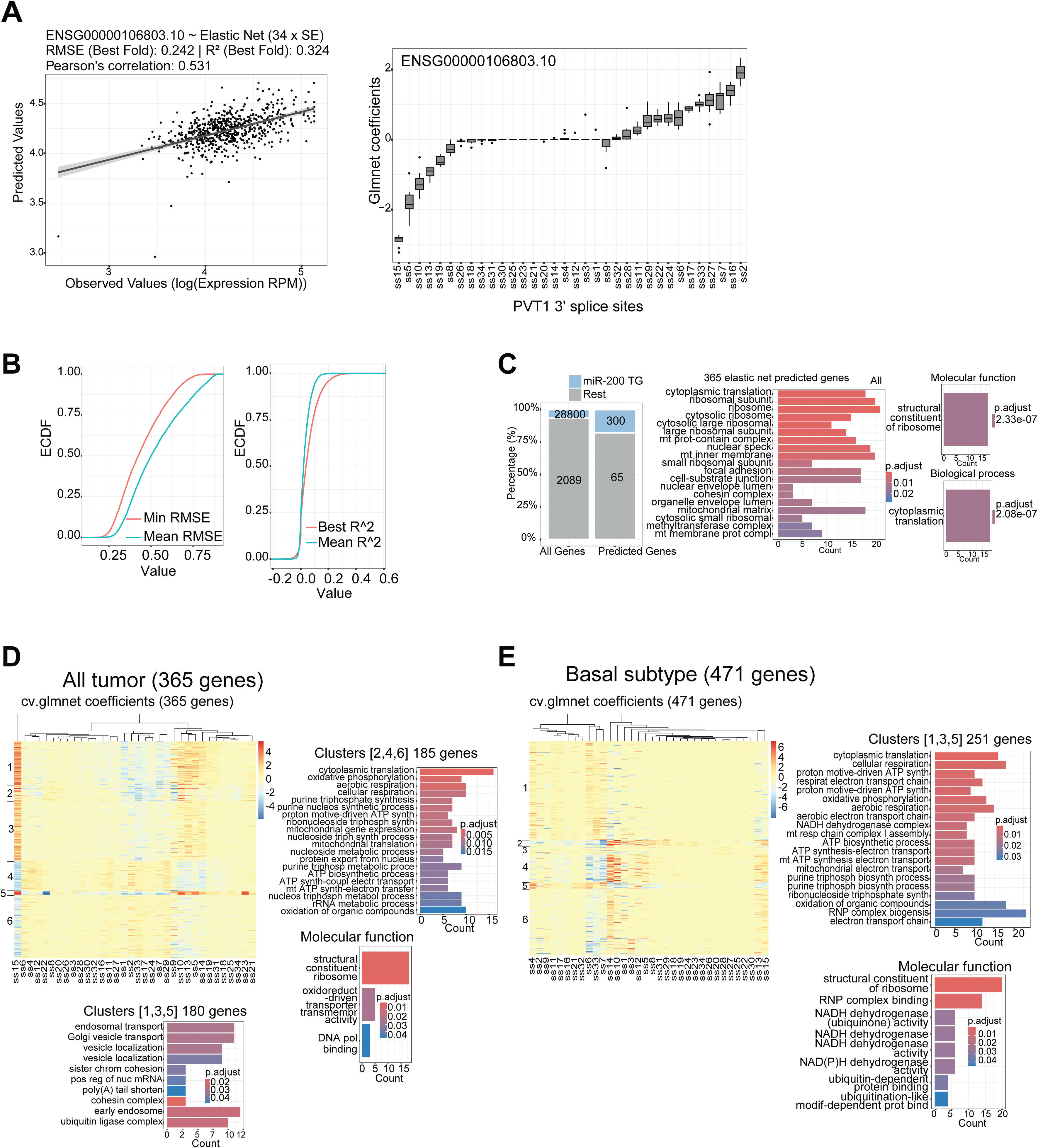
PVT1 splicing activity predicts gene expression in elastic net models. (**A**) Left panel, example of elastic net fit plot of observed values of gene expression (for SEC61B, ENSG00000106803.10) to elastic net-predicted values. Elastic net models for all 30,887 genes were trained in 10 x cross-validation using the splicing efficiency values at 34 PVT1 3’ splice sites as variables. Right panel, boxplot distributions of the glmnet coefficients across 10 folds (from 10 x cross-validation) for the 34 PVT1 3’ splice sites for this specific gene expression model. Positive glmnet coefficients at certain PVT1 3’ splice sites means that splicing efficiency at these sites contributes positively to target gene expression. Negative glmnet coefficients indicate negative contribution of PVT1 splicing efficiency to target gene expression. The bigger the absolute value of a glmnet coefficient, the greater the impact (either positive or negative) of PVT1 3’ splice site splicing efficiency on target gene expression. (**B**) Empirical cumulative distribution function (ECDF) of elastic net (glmnet) performance metrics, model trained using the 34 PVT1 3’ splice site splicing efficiencies, for all the 30,887 assayed genes. Left panel: ECDF of root-mean-square error (RMSE) values, blue line: mean (average) RMSE from 10 x cross-validation, and red line: minimum RMSE (designating best fold from 10 x cross-validation) per gene expression model. Right panel: ECDF of the coefficient of determination (R^2^) values, blue line: mean R^2^ across the 10 folds, and red line: R^2^ from the best fold (defined as the fold with the minimum RMSE). Genes that satisfied i) best-fold R^2^ ≥ 0.2, ii) a negative correlation between R^2^ and RMSE, and iii) 10-fold mean RMSE < Q₃ + 1.5 × IQR were considered as significant, yielding a total of 365 genes predicted as significant when the elastic net models were trained across all tumor samples. (**C**) miR-200 target genes are significantly enriched among the 365 elastic net-predicted genes (Fisher’s exact test p-value 1.51e-14, odds ratio 3.12). Right panels, GO terms enrichment analysis using clusterProfiler [82], terms related to translation are enriched among the 365 elastic net-predicted genes. (**D**) K-means clustering of the glmnet coefficients from the PVT1-splicing based models. The 365 elastic-net predicted genes were clustered based on the elastic net glmnet coefficients into 6 clusters. miR-200 target genes are significantly enriched in clusters 1 and 3 (Fisher’s exact test p-value 8.14e-07, odds ratio 2.28), and translation-related terms are enriched in clusters 4 and 6 (defined by negative contribution of splice sites ss10, ss13, ss14, ss15, and positive contribution of splice sites ss2, ss4, ss6, ss11, ss12, ss29). (**E**) K-means clustering of 471 genes predicted in Basal subtype, using the PVT1 splicing-based elastic net model (glmnet) coefficients of the 34 splice sites as clustering parameters. These 471 genes were predicted as significant (passing the pre-defined cutoffs) by the PVT1 splicing-based elastic net models trained across 155 Basal subtype samples. Translation-related terms were enriched in clusters 1, 3 and 5, defined by positive contribution of splice sites ss4, ss6, ss7, ss33. miR-200 target genes were significantly enriched in cluster 4 (Fisher’s exact test p-value 2.6e-05, odds ratio 3.63) and 6 (Fisher’s exact test p-value 0.0001556, odds ratio 2.57). Clusters 4 and 6 share positive contributions of splice sites ss13, ss14, ss15. No other specific terms were enriched in clusters 2,4,6.

### Distinct PVT1 splice sites predict different groups of genes

To uncover how splicing at distinct 3′ splice sites of PVT1 may influence regulation of predicted target genes, we applied k-means clustering to group target genes based on the regression coefficients obtained from PVT1 splicing-based linear regression (lm) and elastic net models (glmnet). These coefficients, reflecting the contribution of each splice site to target gene expression, served as input features to identify clusters of genes sharing similar splicing-dependent regulatory profiles. Using the glmnet coefficients of all 34 splice sites in k-means clustering of the 365 predicted genes (passing the performance cutoffs; Figure 3A-C) across 617 BRCA tumor samples, we identified 6 gene clusters defined by distinct contributions of 3’ splice site splicing efficiency (Figure 3D).

More specifically, we found 2 main clusters, 1 and 3, (and a smaller one, cluster 5), that are defined by positive contribution (i.e. positive glmnet coefficients) of splice sites ss15, ss10, ss13, ss5, ss14, ss21, ss23, and negative coefficients of splice sites ss1, s2, ss33, ss17, ss24, ss7, ss29. Features with positive glmnet coefficients indicate a positive contribution to target gene expression, meaning that a higher PVT1 splicing efficiency at those splice sites enhances gene expression. Conversely, PVT1 splice sites with negative coefficients are associated with lower target gene expression. miR-200 target genes are enriched in clusters 1 and 3 (Fisher’s exact test p-value 8.14e-07, odds ratio 2.28). Different GO terms are enriched in clusters 1 and 3, compared to clusters 4 and 6, suggesting that splicing activity at different PVT1 3’ splice sites (and/or alternative PVT1 splicing isoforms) are involved in the regulation of different sets of genes (Figure 3D).

Apart from across all BRCA tumor samples, we also trained splicing-based elastic nets within pam50 subtypes, followed by k-means clustering of the predicted genes based on the respective glmnet regression coefficients (Methods) (Basal subtype in Figure 3E; LumA and LumB in Supplementary Figure S4C-D). For the Basal subtype (155 samples with pam50 subtype ‘Basal’), we found 471 genes passing the performance cutoff, clustered in 6 groups by k-means clustering using the 34 glmnet coefficients (Figure 3E). Like ‘all_BRCA_tumor’ (Figure 3D), Basal clusters are defined by sets of 3’ splice sites anticorrelating in splicing efficiency contribution. Cluster 4, enriched with miR-200/205 target genes (Fisher’s exact test p-value 2.6e-05, odds ratio 3.62), is defined by positive contribution (positive glmnet coefficients) of splice sites ss5, ss10, ss12, ss13, ss14, ss15, and negative contribution (negative glmnet coefficients) of splice sites ss6, ss7, ss33.

For LumA clustering, mir-200 target genes are significantly enriched in clusters 3 and 6 (Fisher’s exact test p-value 1.142e-12, odds ratio 7.024), which are defined by positive contribution of splice sites ss5, ss9, ss10, ss13, ss14, ss15 (Suppl. Figure S4C). Among all genes predicted by PVT1 splicing in LumA (620 genes passing performance cutoff), GO terms enriched are skewed toward RNA localization, RNP organization, nuclear export and translation, while enriched terms varied in clusters 3 and 6 (Suppl. Figure S4C lower panels). Similar for LumB clustering (158 genes), miR-200 target genes are only enriched in clusters 1 and 3 which are defined by positive contribution (positive glmnet coefficients) of splice sites ss13, ss14, ss15 (Fisher’s exact test p-value 0.0137, odds ratio 3.055; Suppl. Figure S4D).

That the clusters are defined by two sets of splice sites that anti-correlate in splicing efficiency contribution could suggest that these belong to at least two distinct PVT1 alternative splicing isoforms involved in regulation of predicted target gene expression, a possibility that we explored further via alternative splicing isoform quantification and differential expression analysis of PVT1 transcripts.

### PVT1 splicing activity distinguishes between tumor subtypes, and is prognostic for tumor progression and treatment outcome

Next, we asked whether PVT1 splicing at specific PVT1 3’ splice sites could be a distinctive feature between BRCA tumor subtypes. Classification analysis was performed using Random Forests and the splicing efficiency values at the 34 3’ splice sites as features, in 10 x cross-validation. Models distinguishing Her2 from all other subtypes (non-Her2) performed best (best average accuracy), followed by classification models of Basal versus rest (non-Basal), and LumB versus rest (non-LumB) (Figure 4A-B).

**Figure 4.**
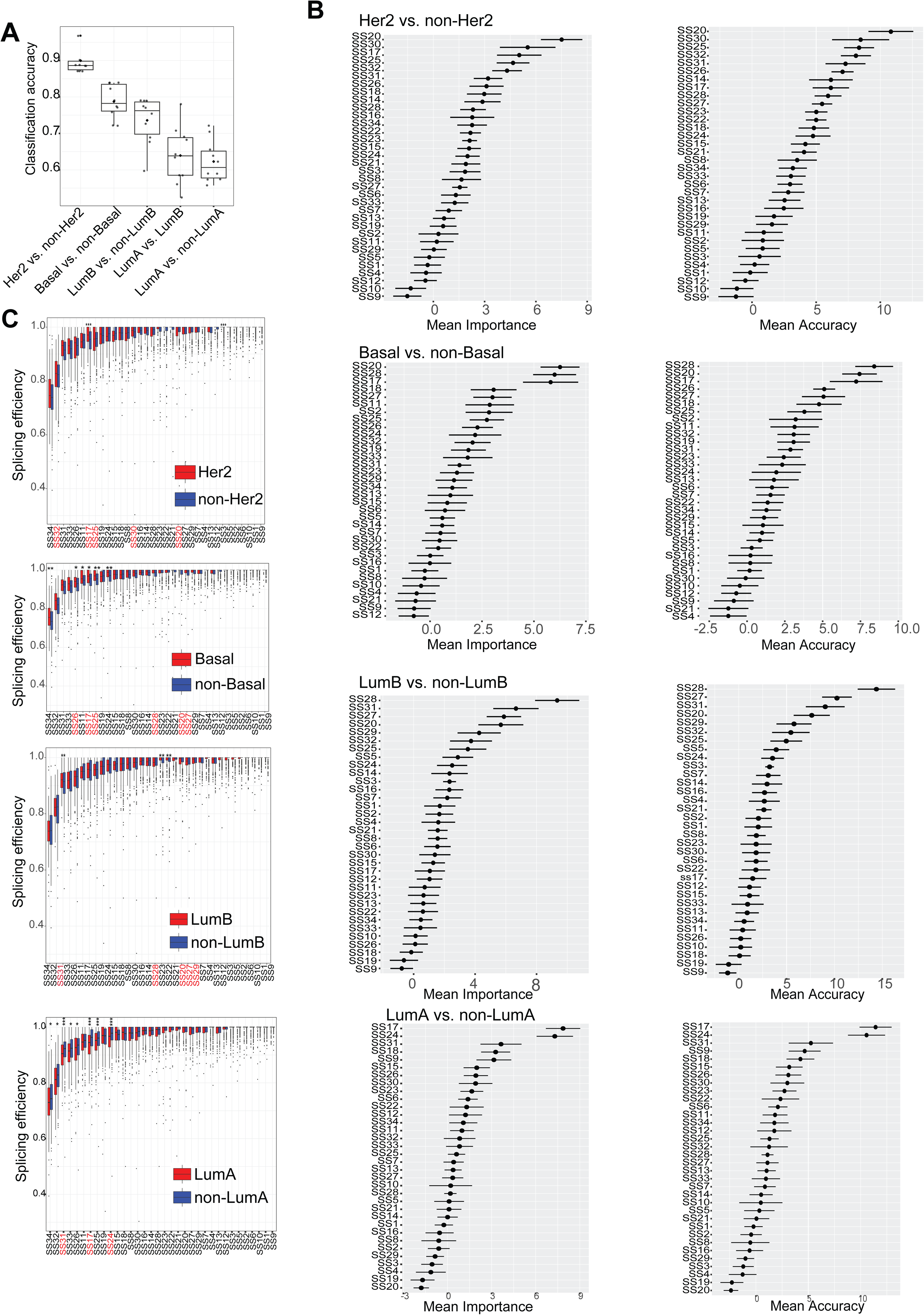
PVT1 splicing activity distinguishes BRCA subtypes (and reflects tumor aggressiveness). (**A**) Distribution of classification accuracy values from 10-fold cross-validation of Random Forest models trained on PVT1 3’ splice site splicing efficiencies to distinguish each subtype from the rest. Highest performance was obtained for Her2, followed by Basal and LumB, while LumA showed mean classification accuracy < 0.65. (**B**) Feature mean accuracy and mean importance plots of Random Forest subtype classification models. Top-ranked splice sites contributing to subtype classification (feature mean importance and mean accuracy scores are plotted; mean Gini scores are shown in Suppl. Figure S5B), uncovering subtype-specific splicing signatures. (**C**) Comparison of PVT1 3’ splice site splicing efficiency values across binary classifications. Top-ranked features from the respective Random Forest classification models are marked in red font.

Top features distinguishing ‘Basal’ from the rest of tumor subtypes are ss20, ss28, ss17, ss27; splicing efficiency at those top feature-splice sites is higher in Basal compared to rest subtypes (Figure 4C). Of note, higher splicing at ss27 and ss28 associates with a worse prognostic outcome (i.e. worse survival rate) in Basal subtype, and across all tumor samples (Suppl. Fig. S6). Among top features distinguishing ‘Her2’ from the rest of tumor subtypes are ss20, ss30, ss25, ss17, ss25, ss32. Similarly, splicing efficiency at those sites is higher in Her2 compared to the rest of samples (Figure 4B-C). On the other hand, top distinctive features (ss20, ss27, ss28, ss29) distinguishing LumB from rest subtypes (with an average model classification accuracy from 10 x cross-validation ∼0.75, median ∼0.77, Figure 4A) exhibit, on average, lower splicing efficiency in LumB compared to the rest of the samples (Figure 4C).

Finally, PVT1 splicing-based classification models are less accurate in distinguishing LumA from rest of tumor samples, or even LumA from LumB, (mean model accuracy from 10 x cross-validation < 0.65, Figure 4A). Top important features, ss17 and ss24, show a lower splicing efficiency in LumA compared to the rest of tumor samples (Figure 4C, bottom panel).

Given that Her2-positive and Basal-like subtypes are more aggressive and often associated with poorer prognosis compared to LumA and LumB, with the Basal triple-negative subtype being most aggressive and metastatic [56] (Suppl. Figure S6F), these findings suggest that subtype-specific alterations in PVT1 splicing efficiency may reflect the molecular progression of breast cancer and may serve as a potential (diagnostic) biomarker of tumor aggressiveness and subtype identity.

The splicing-based classification performance among BRCA subtypes (Her2 > Basal > LumB > LumA) could reflect increasing similarity in PVT1 splicing profiles to the rest of the tumor population, with splicing activity lacking distinctive power in classifying LumA from the rest of the tumor samples (Figure 4A). Thus, PVT1 splicing may vary across tumor subtypes, particularly in Her2-positive and Basal tumors. This could indicate either subtype-specific regulatory splicing programs and/or differential usage of alternative PVT1 splicing isoforms related to subtype biology. Moreover, splicing efficiency at top splice sites driving classification (i.e. displaying subtype-selective patterns), is higher in Basal and Her2 tumors (Figure 4C), suggesting that these splice sites may be preferentially used in the Basal (and Her2) subtype, and that the corresponding PVT1 isoforms may be upregulated or stabilized in Basal/Her2 tumors (which are fast progressed and aggressive subtypes, Suppl. Fig. S6F).

To determine if the subtype-specific differences in PVT1 splicing efficiency values are reflected in alternative transcript isoform usage, and if the relative abundance of alternative PVT1 transcript isoforms can drive tumor classification, we quantified the expression of alternative PVT1 transcript isoforms across all BRCA samples using Salmon [57] (Suppl. Figure S7).

Overall, the production of spliced PVT1 transcripts, inherently linked to splicing activity at the PVT1 locus, is associated with improved prognostic outcome, i.e. longer progression-free and disease-free intervals across all patients expressing certain alternative transcript isoforms above the median (Suppl. Figure S8). Among all PVT1 transcripts profiled in survival analysis, ENST00000665372.1 is weakly associated with a worse prognostic outcome (at progression-free interval-PFI p-value 0.07). Of note, this is the only Salmon-quantified transcript including exon 2 and is low-expressed across samples (Suppl. Figure S2A, Suppl. Figure S7B).

#### Splicing efficiency does not always correlate with alternative transcript isoform abundance

We frame two hypotheses: i) that splicing regulation of PVT1 is not only altered in cancer versus normal tissue (i.e. is not only altered on tumor onset), but also shows subtype-specific signatures, especially in Her2 and Basal-type tumors; and ii) Basal-specific splice junctions may generate PVT1 isoforms with unique functional potential, perhaps linked to the aggressive phenotype of Basal subtype. To address these hypotheses, we performed survival analysis to correlate splicing efficiencies at top classification features (Figure 4) with clinical outcomes (disease-free interval DFI, progression-free interval PFI; Methods) (Suppl. Fig. S6). We find that reduced splicing efficiency at certain splice sites (ss7, ss8) correlates with a worse prognostic outcome (DFI, PFI), suggesting that maintaining PVT1 splicing activity (at those splice sites) may be protective against tumor progression or contributing positively in response to treatment. On the other hand, higher splicing efficiency at ss27, ss28 correlates with worse prognosis across all tumor samples, which is in line with the classification analysis since those were among top features classifying Basal from the rest of subtypes (Suppl. Figure S6, Figure 4).

Within Basal, splicing efficiency at ss7 and ss8 is protective (Suppl. Figure S6B), (which may correlate with a putative role of the expressed host transcript ENST000520913.2 in disease-free interval after treatment, although not statistically significant). Of note, splicing at one adjacent splice site (ss10) is also protective in Basal, (although ss10 was not detected in any of the 19 top Salmon-extracted and quantified transcripts). Splicing at ss27, ss28, and ss31 is also associated with a worse prognostic outcome (shorter PFI and DFI) in LumB (Suppl. Figure S6D). Finally splicing efficiency at the top classifying feature of LumA, ss17 (Figure 4B-C), associates with a worse prognostic outcome in this subtype (DFI p-value 0.02, Suppl. Figure S6E).

Like the PVT1 splicing-based classification models, Random Forest models trained on alternative transcript isoform expression (with 10 x cross-validation) showed the lowest performance (mean accuracy) in classifying LumA from the rest subtypes (non-LumA), preceded by LumB vs. non-LumB, Basal vs. non-Basal, and Her2 vs. non-Her2 performing the best (Suppl. Figure S7A).

Among top features classifying Basal from non-Basal, two transcripts lacking common exon_3, ENTS00000692980.1 and ENST00000690350.1, are upregulated (non-significant) in Basal, with concomitant significant reduction of ENST00000671092.1, ENST00000517838.6, and ENST00000520913.2 that contains exon_3 (Suppl. Fig. S7E, S2A). The 3’ splice site of excluded (skipped) exon_3, ss11, shows however increased splicing efficiency in Basal (Figure 4C), which would, on a first sight, suggest enhanced inclusion of exon 3. Among top expression-based classification features, increased expression of ENTS00000692980.1 in Basal is consistent with increased splicing at the overlapping ss25, serving as a top splicing-based classification feature (Suppl. Figure S7E, Figure 4B-C). Thus, splicing-efficiency-based and alternative-isoform expression-based classification models may agree to some degree, although, in general, splicing efficiency does not always correlate with corresponding alternative transcript isoform abundance. Moreover, alternative splicing profiling at splice-junction level using MAJIQ v2 [58] did not detect exclusion of exon 3 as a significant skipping event in Basal (not shown), and splicing efficiency values per *se* are not effectively predicting relative alternative transcript isoform abundance, neither in linear regression nor in elastic net models (Suppl. Fig. S9E). Thus, apart from local splicing activity or splicing efficiency per *se* at individual splice sites, additional mechanisms (e.g. transcript stability/turnover, promoter usage, differential subcellular/subnuclear localization and protein interactions) are involved in defining the final relative abundance of mature alternative splicing isoform transcripts.

#### Splicing-based models outperform transcript-based models in predicting gene expression

We next examined the impact of alternative PVT1 isoform expression in gene regulation by training elastic nets with 10 x cross-validation (across all 669 BRCA samples) to predict gene expression in genome-wide screens (∼30,887 genes). 611 genes passing the cutoffs (Methods) are predicted at high confidence, with enriched GO terms related to DNA repair (Suppl. Figure S9A). K-means clustering of the predicted genes based on the elastic net glmnet coefficients revealed significant enrichment of miR-200 target genes (odds ratio 4, Fisher’s exact test p-value = 4.42e-12) in a cluster defined by concurrent positive contribution of ENST00000667630.1, ENST00000671092.1 and ENST00000658840.1 (Suppl. Fig. S9B-C; first and third are bi-exonic transcripts, Suppl. Fig. S2A). Notably, the first two transcripts (ENST00000667630.1, ENST00000671092.1) show positive correlation with overall intron retention at the locus (Suppl. Figure S3B), suggesting that the inferred regulation of the enriched miR-200 target genes may be indirect through post-transcriptional mechanisms linked to PVT1 splicing. This interpretation is supported by the presence of exclusively intronic miR-200 seed sites at the locus underscoring the possibility that PVT1-mediated regulation of miR-200 target genes operates through a ceRNA-like activity facilitated by inefficient co-transcriptional processing and chromatin-associated intron retention.

When comparing the performance between splicing-based and transcript-based predictive models, the former performed better, yielding lower RMSEs across cross-validation folds (Suppl. Figure S9D). The lack of a substantial overlap in the gene sets predicted by the splicing-based and transcript-based models is consistent with recent findings suggesting that differential splicing and differential expression often regulate distinct biological programs [54]. Thus, our results reinforce the notion that splicing contributes distinct regulatory signals not captured by expression-level models alone, underscoring the importance of transitioning from gene-centric to isoform- and splicing-centric transcriptomic interpretations.

### Causal inference analysis uncovers the relevance of PVT1 splicing activity and its functional role in shaping regulation of target gene expression

A key question arising from the predictive capacity of PVT1 splicing activity (Figure 3, Suppl. Fig. S2, S4) is whether changes in its splicing pattern are merely correlated with, or rather causally linked to, the regulation of distal gene expression. To address this, we leveraged whole-exome and whole-genome sequencing (WXS/WGS) data from TCGA BRCA to identify tumor-specific single nucleotide variants (SNVs) clustered in proximity to the 3’ splice sites of PVT1 (Methods). These SNVs were used as naturally occurring perturbations to test whether variation in PVT1 splicing efficiency at specific splice sites modulates gene expression of predicted target genes. Tumor-specific somatic SNVs were used before in genome-wide analysis to identify ‘driver’ mutations in lncRNAs promoting cancer cell fitness [59].

By integrating splicing efficiency data with matched tumor-specific SNVs, we implemented a causal inference framework to unravel the impact of splice site-associated SNVs on gene expression (Figure 5A). This approach allowed us to dissect potential functional consequences of splicing modulation and provided mechanistic insights linking PVT1 splicing dynamics to the regulation of gene expression at distal genomic loci.

**Figure 5.**
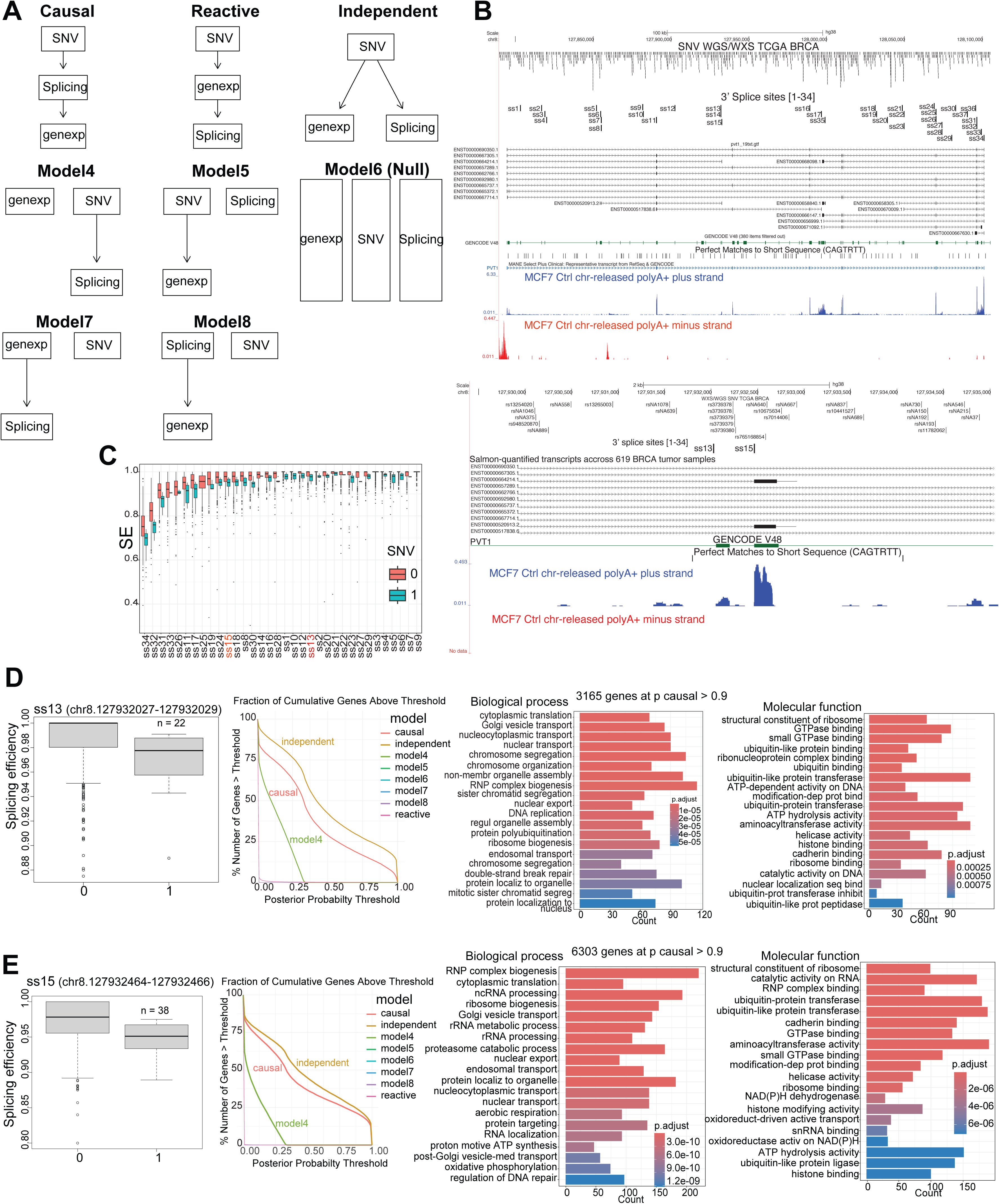
Causal inference analysis identifies specific PVT1 splice sites in affecting distal gene expression. (**A**) Schematic of Bayesian network models tested to infer causal relationships between SNVs, splicing efficiency at PVT1 3’ splice sites, and gene expression. For each of the 30,887 genes and per splice site we tested 8 models: Causal (SNV → splicing → gene expression), Reactive (SNV → gene expression → splicing), Independent (SNV → splicing, SNV → gene expression), Model 4 (SNV → splicing, gene expression varies independently), Model 5 (SNV → gene expression, splicing varies independently), Model 6 – Null (no relationship between the nodes), Model 7 (gene expression → splicing, SNV independent), Model 8 (splicing → gene expression, SNV independent). (**B**) Overview of SNV-to-splice site assignment: Tumor-specific SNVs identified from WXS/WGS data (n = 592 TCGA BRCA tumor samples) were assigned to the closest PVT1 3’ splice site (out of 34, with *bedtools closest*), resulting in a binary SNV matrix per splice site, per sample. (Note that not all samples carried all depicted SNVs; 80 % of the samples carried less than 5 SNVs, either WGS or WXS, at the PVT1 locus). For each SNV–splice site pair, the presence or absence of an SNV was tested for association with changes in splicing efficiency (See also Suppl. Figure S10A). (**C**) Boxplots illustrating splice site-specific splicing efficiency (SE) changes (reduction) associated with nearby SNVs. Samples carrying SNVs at PVT1 3’ splice sites showing reduction in splicing efficiency (SE < median of all SE values) were assigned ‘1’. Splice sites carrying no SNVs or carrying SNVs but not showing reduction in splicing efficiency were assigned ‘0’. The respective results for SNV-splice sites associated with an increase in splicing efficiency are shown in Supplementary Figure S10A. (**D-E**) Causal inference analysis results for the splice sites ss13 and ss15. (D) From the 592 BRCA samples genotyped, 22 samples had an SNV assigned to ss13 associated with a reduction in splicing efficiency. Across all 30,887 SNV-splicing-gene expression triplets tested, variance in the expression of 3165 genes was best explained by the causal model at a posterior probability > 0.9. Gene ontology terms related to translation and RNA processing are significantly enriched among those genes best explained by the causal model. Similar for ss15 SNV-splicing-gene expression triplets (E), translation and RNA processing terms are significantly enriched among the genes best explained by the causal model, whereas no specific terms were enriched among 7159 genes best explained by the independent model (see also Suppl. Figure S10B). As shown in the cumulative plots for ss13 (D) and ss15 (E), for all assayed splice site (SNV – splicing – gene expression) triplets, most genes were best explained by the independent model (SNV → PVT1 splicing, SNV → gene expression), followed by the causal model, while a smaller fraction of genes were best explained by model 4 (SNV → splicing, gene expression varies independently). See also Supplementary Figure S10.

In detail, to uncover whether PVT1 splicing exerts a causal effect on gene expression in *trans*, we leveraged tumor-specific SNV annotated in TCGA BRCA WXS and WGS data (Figure 5B). From the 617 BRCA tumor samples analyzed, 592 were genotyped either by WXS or WGS, and 410 of those harbored at least one SNV in the PVT1 locus. Each SNV was assigned to the closest PVT1 3’ splice site (Methods), resulting in binary annotations across the 34 splice sites: samples with an SNV near a given site were assigned ‘1’, and those without were assigned ‘0’. The remaining 182 samples had no SNVs in the PVT1 locus and were assigned ‘0’ at all splice sites. We noted that for most splice sites, the presence of nearby SNVs was associated with changes in splicing efficiency, with statistically significant differences observed in approximately one-fifth of the splice sites (Suppl. Figure S10A). To increase resolution, we split the samples into groups bearing mutations associated either with splicing upregulation or downregulation, achieving statistical significance in mutation-associated changes in splicing efficiency for most splice sites (Figure 5C, Suppl. Figure S10B-C).

For each PVT1 3’ splice site, we built a dataset with 592 samples and three variables: (1) presence/absence of an SNV near the splice site (“SNV”), (2) splicing efficiency (“Splicing”, continuous value from 0 to 1), and (3) expression of a candidate target gene (“genexp”, reads per million - RPM). We formulated eight Bayesian network models to evaluate different relationships among those variables (Figure 5A); these include the causal model (“SNV → Splicing → Expression”), where splicing mediates the SNV’s effect on gene expression; the reactive model (“SNV → Expression → Splicing”), whereby distal gene expression influences PVT1 splicing; and the independent model, where SNV affects splicing and expression independently. In Model 4, the SNV affects splicing at the analyzed PVT1 3’ splice site, whereas distal gene expression varies independently.

Posterior probabilities were computed for each model using *bnlearn* [60], and we screened across all ∼ 30 K genes (30,887 filtered genes, Methods) to identify SNV – splice site – gene triplets best explained by the causal model (at posterior probability > 0.9). Thus, this in *silico* perturbation strategy allowed us to test whether PVT1 splicing efficiency acts as a functional intermediary (‘mediator’) between local sequence variation and distal gene expression.

Of the 34 splice sites analyzed, SNV-mediated perturbations in PVT1 splicing efficiency downregulated at splice sites ss13 and ss15 were associated with changes in the expression of 3165 and 6303 target genes respectively, based on a causal inference posterior probability > 0.9 (Figure 5D-E). Common GO terms enriched in those two datasets explained by the causal model are related to translation regulation, and miR-200 target genes are enriched at Fisher’s exact test p-value < 2.2e-16 and odds ratio 3.75 and 3.34, respectively (Figure 5D-E). This result suggests a mechanistic insight for a functional link between PVT1 splicing activity and regulation of miR-200 target genes in *trans*, via a ceRNA-mediated miRNA (miR-200) axis.

Those two specific splice sites, ss13 and ss15, were also captured as significant in the elastic net coefficient-based k-means clustering of target gene expression, defining clusters where miR-200/205 target genes were significantly enriched (Figure 3D-E), substantiating their role in shaping gene expression regulation in *trans*.

In addition, SNV-based inference analysis identified splicing perturbations at splice site ss24 and ss16 as causally linked to changes in the expression of a distinct set of genes. Those were strongly enriched for GO terms related to immune response and inflammatory signaling pathways (Suppl. Fig. S10D-E), suggesting that specific PVT1 splicing events may play broader roles in shaping gene regulatory programs associated with tumor–immune interactions.

Notably, although for all splice sites examined via causal inference analysis a relatively bigger fraction of genes is explained by the independent model at posterior probability > 0.9 (Figure 5D-E, Suppl. Figure S10), there is no significant enrichment of specific terms among those datasets, or irrelevant terms appear at high p-adjusted (Suppl. Figure S10). This suggests that only SNV-driven perturbations in splicing with strong causal support are likely to exert coordinated regulatory effects on gene expression programs, particularly those involved in translation and miRNA-mediated post-transcriptional regulation. We note here that i) similar terms were enriched in gene datasets explained by the causal model upon SNV-mediated splicing up- or down-regulation of the same splice sites, and ii) apart from the absence of specific terms, miR-200/205 target genes are not enriched either in any of the datasets explained by the independent models (Suppl. Figure S10B-C).

Together, these results suggest that specific SNV-mediated perturbations of PVT1 splicing efficiency at select 3’ splice sites causally influence distal gene expression, supporting a model in which PVT1 splicing may function as a regulatory hub, plausibly via chromatin-associated intron retention, and with subtype- and context-specific downstream effects, including miRNA-mediated post-transcriptional regulation and modulation of transcriptional programs in cancer.

### Cross-tissue generalizability of PVT1 splicing-based gene expression models

To assess whether the relationship between gene expression and PVT1 3’ splicing efficiency observed in breast cancer models applies to other tissues and cancer types with similar PVT1 expression levels, we applied our regularized linear regression (elastic net) models on model-unseen data using five independent TCGA tumor datasets: prostate, ovary, uterus, testis, and adrenal gland (Methods). These datasets were selected based on comparable PVT1 expression identified in respective healthy tissues (GTEx) and verified in TCGA (Suppl. Fig. S1C, Suppl. Fig. S11A). For each of the five datasets we computed the per-gene RMSE (by applying all the 30,887 elastic net models trained in BRCA, one model per gene) and compared those errors with their predicted in BRCA counterpart genes (Suppl. Figure S11B-C). We used three gene sets to examine generalizability: i) across the full gene dataset complement expressed in each tissue (Suppl. Fig. S11B, Suppl. Fig. S11C left panel; Methods), ii) within the filtered subset of the 365 genes that satisfied significance criteria in BRCA (Figure 6A-F, Methods), and iii) for sets of genes generating an R^2^ > 0.2 (Suppl. Figure S11C, middle panel), or R^2^ > 0.1 (Suppl. Figure S11C, right panel) within each tissue by applying the BRCA-trained models (results summarized in Suppl. Table S1).

**Figure 6.**
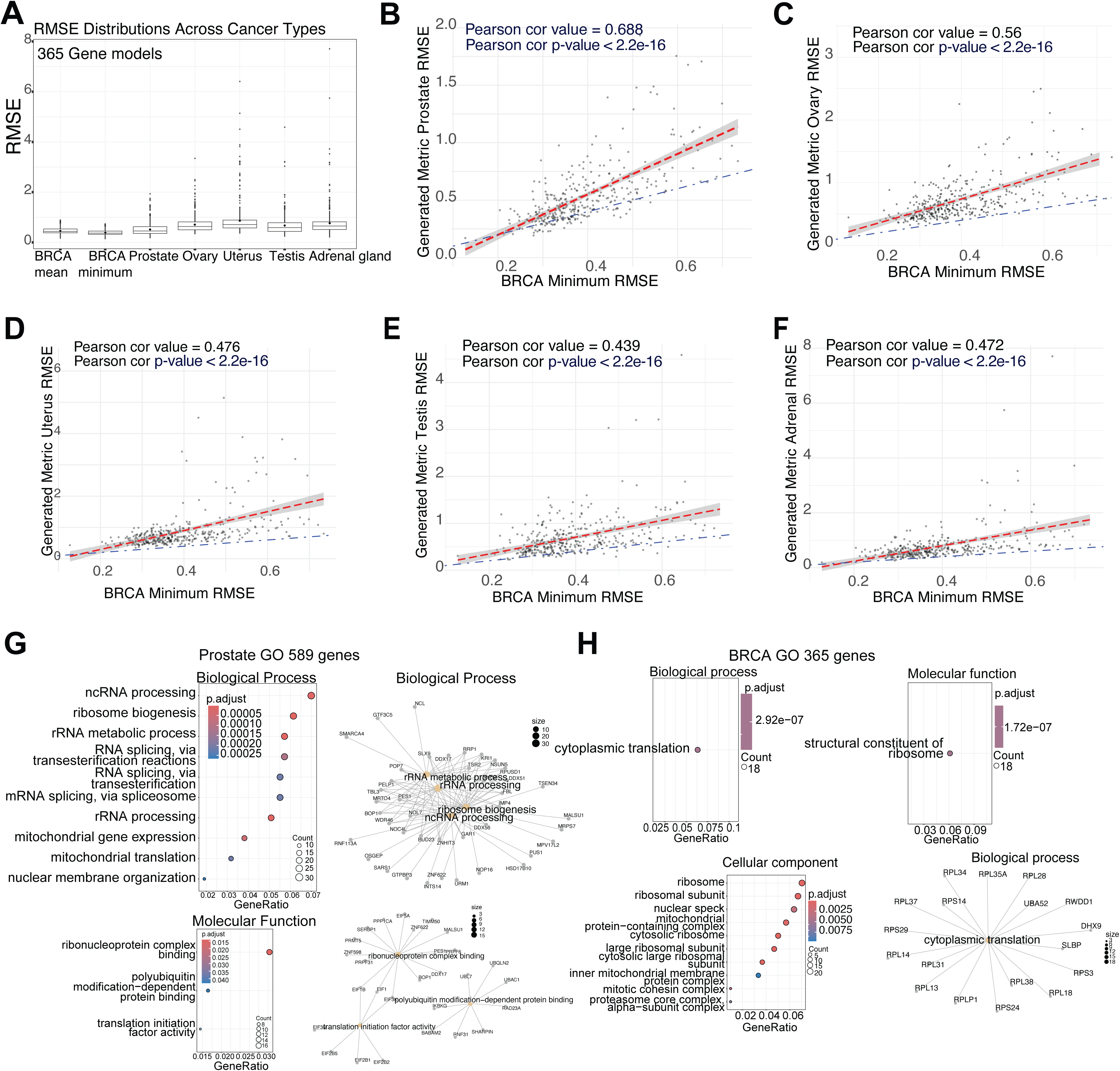
Generalizability of PVT1 splicing-based gene expression models trained in BRCA on unseen data of different cancer types. (**A**) Comparison of PVT1 splicing-based gene expression model performance: RMSE boxplot distributions of the 365 gene models trained in BRCA and applied in different cancer types. RMSE boxplot distributions for 365 gene models trained in BRCA, both mean and minimum RMSE from 10 x cross-validation are shown in BRCA, alongside cross-tissue RMSE values from applying the BRCA-trained models to unseen tumor types (prostate, adrenal, ovary, uterus, testis). (**B-F**) Model generalizability across five TCGA tumor cohorts: Elastic net models trained in BRCA for the 365 selected genes were applied to unseen tumor datasets (using PVT1 3’ splice site splicing efficiencies as features and gene expression as response). The resulting per-gene prediction error (RMSE) (*y*-axis) was compared to BRCA performance (the minimum RMSE from the best fold of 10 x cross-validation in BRCA, on the *x*-axis), using Pearson correlation, in prostate (B), ovary (C), uterus (D), testis (E) and adrenal gland (F). Red lines denote linear regression fits with 95% confidence intervals (gray shading), while blue identity lines (y = x) indicate equal performance between BRCA and the test tissue. See also Suppl. Figure S11. (**G-H**) Gene ontology (GO) enrichment analysis and cnetplots of predicted gene sets in (G) prostate (n = 589 with performance exceeding R^2^ > 0.1) and (H) BRCA (n = 365 that passed filtering criteria). Shared enriched terms (Biological Process) include RNA processing and translation-related categories.

Overall, the BRCA-built PVT1 splicing-based elastic nets found best model applicability in the prostate compared to the other four unseen datasets (Figure 6A-F). By setting a cutoff at R^2^ > 0.2, prostate yielded 158 genes (Suppl. Figure S11C middle panel) (followed by the adrenal gland with 15 genes, 7 for ovary, zero for uterus and 1 for testis; Suppl. Table S1). At R^2^ > 0.1, prostate yielded 589 genes (Suppl. Fig. S11C right panel), followed by 154 for adrenal, 46 for ovary, 80 for uterus, 112 for testis, all showing similar positive RMSE correlations to BRCA, comparable to full gene datasets, ranging from r = 0.85 (testis) to 0.94 (adrenal) and 0.98 (ovary) (not shown-summarized in Suppl. Table S1). Moreover, common GO terms related to translation and RNA processing are enriched among the 589 prostate- and the 365 breast-cancer predicted genes (Figure 6G-H), which likely suggests that common post-transcriptional regulatory programs may underlie the cross-tissue generalization of BRCA-trained PVT1 splicing-based predictive models.

Collectively, these findings suggest that PVT1 3’ splicing-based gene expression models trained in BRCA generalize most effectively to prostate cancer, with moderate but consistently positive applicability in other PVT1-expressing tissues. Model performance correlates with tissue-specific gene expression levels and sample size, and highlights shared post-transcriptional regulatory programs, particularly in the prostate and breast tissues (Figure 6G–H, Suppl. Note S3). High RMSE correlations across all five tissues indicate that absolute error patterns learned in BRCA are preserved, pointing to a shared dynamic range of PVT1 splicing–driven predicted gene expression. However, robust cross-tissue generalization is restricted to gene sets with sufficient expression variance in both source (BRCA) and target tissues. Among the examined cohorts, prostate (and to a lesser extent, ovary) showed the strongest generalization, supported by shared gene ontology enrichment for splicing/RNA processing, and translation-related common terms (Figure 6G–H, Suppl. Note S3). In contrast, uterus, testis, and adrenal gland showed weaker overlap with BRCA-predicted genes and minimal term enrichment, suggesting that further model refinement may be necessary to enhance generalization to these tissues (see also Suppl. Note S1).

### PVT1 is a top candidate among chromatin-associated lncRNAs for splicing-modulated intronic ceRNA activity

Our machine learning models uncovered PVT1 as a lncRNA whose splicing activity is predictive of distal target gene expression, identified through genome-wide in *silico* screens (Figure 3). For at least some 3’ splice sites, including ss13 and ss15, this predictive relationship was validated by causal inference using tumor-specific SNVs that perturb local splicing efficiency (Figure 5, Suppl. Figure S10). Notably, miR-200/205 target genes were significantly enriched among genes regulated by splicing efficiency at these splice sites (Figures 3, 5; Suppl. Fig. S4), prompting us to examine a potential ceRNA-based mechanism. We presume that PVT1 may act as a chromatin-associated lncRNA with intron-retained ceRNA activity, sequestering miR-200/205 molecules through intronic seed sites, thereby modulating post-transcriptional regulation of target genes [61] (Figure 7E, Model). Consistent with this hypothesis, perfect 8-mer and 7-mer miR-200 seed matches are located exclusively within PVT1 introns (Suppl. Figure S1A-B), suggesting that splicing modulation, and specifically intron retention, could influence the availability of these binding regions and, consequently, the competing endogenous ceRNA activity of PVT1.

**Figure 7.**
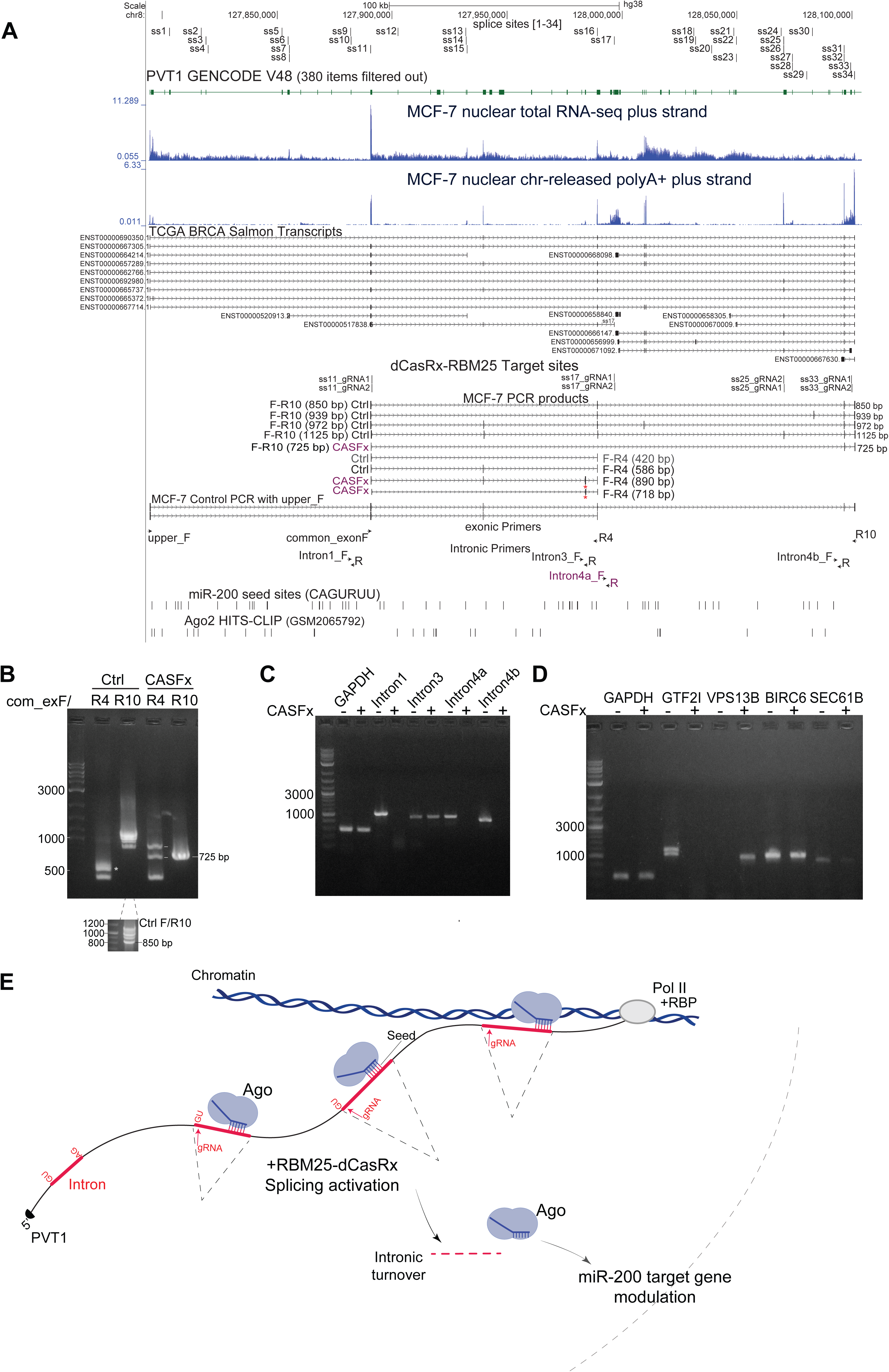
Artificial splicing activation of PVT1modulates its chromatin-associated intronic RNA and alters expression of miR-200 target genes in *vitro*. (**A**). UCSC overview of the PVT1 locus showing positions of gRNA oligos designed to target four splice sites (ss11, ss17, ss25, s33) for splicing activation with dCasRx-RBM25 (CRISPR Artificial Splicing Factor, CASFx). Positions of exonic and intronic primers used in (B-C) are indicated. (**B**) PCR analysis on cDNA from nuclear chromatin-released dT-primed RNA in MCF-7 cells 48 h after CASFx transfection shows altered PVT1 splicing patterns. In PCR with primer pair common_exonF/R4 a major band of 586 bp (star marked) is shifted to two alternative splicing products 718 and 850 bp by activated 3’ splice site exon inclusion (marked with a red star in UCSC track in (A); see also Suppl. Figure S13F). A lower 420 bp band is not altered. PCR with common_exonF/R10 detects several bands in a range 800-1200 bp corresponding to several alternative transcript isoforms detected in control condition. Upon CASFx, all alternative splicing activity downstream of ‘common_exon’ is likely suppressed and PCR with common_exonF/R10 detects one major splicing isoform with a 725 bp PCR product. (**C**) Upon CASFx, intronic signal from introns ‘1’ and ‘4’ is eliminated on random-primed chromatin-associated RNA (but not on nuclear chromatin-released dT-primed RNA; Suppl. Figure S13B). (D) Semi-quantitative PCR on cDNA from nuclear chromatin-released dT-primed RNA with gene-specific primers detects alterations in GTF2I, VPS13B, SEC61B (see also Suppl. Fig. S13C and S14). (**E**) Model of chromatin-associated intronic ceRNA activity mediated by chromatin-associated PVT1 and expected effect of splicing enhancement. PVT1 intronic miR-200 seed sites sequester miRNAs in association with nuclear Ago. Artificial splicing enhancement of PVT1 with CASFx results in increased turnover of spliced-out intronic sequences and releases miR-200/Ago effective availability.

To uncover whether this intronic ceRNA functional potential is unique to PVT1 or part of a broader splicing-based (or intron-retention-based) regulatory hub, we conducted a genome-wide search for transcripts harboring miR-200 seed sites in their introns. Genes were ranked based on intrinsic features, including average expression across the BRCA cohort, intron retention levels, and intronic seed motif count (Methods; Suppl. Table S3). As expected, lncRNAs generally showed higher intron retention compared to protein-coding genes (Suppl. Figure S12A), likely due to an overall less efficient RNA processing [62]. Among top-ranked candidates, PVT1 emerged as the best-scoring lncRNA with intronic ceRNA potential, followed by FTX, another chromatin-associated lncRNA, DLEU2 and CASC15. Among protein-coding genes, top-ranked candidates include RAD51B, SLC30A8, and ERBB4 (Suppl. Figure S12B).

To test whether splicing of those candidates mirror the gene-expression predictive capacity of PVT1, we repeated the splicing efficiency-based modeling pipeline (Figure 1, Methods) for each one of them. For FTX, elastic net models identified 171 target genes whose expression could be predicted by its 3’ splice site splicing efficiency, with significant enrichment for miR-200/205 targets (Fisher’s exact test p-value 1.42e-05, odds ratio 2.62). Rest lncRNAs tested (DLEU2, CASC15) rendered no significant miR-200/205 enrichment in predicted genes (Suppl. Table S2). Among the 6 mRNAs tested, despite their similar intronic seed site density, intron retention and splicing efficiency values (Suppl. Figure S12B-C), only PTPRT rendered some enrichment in predicted miR-200/205 target genes, albeit less profound (odds ratio 1.99, Fisher’s exact test p-value 0.00178), and while PTPRT splicing-based elastic net predicted genes (n = 201) show significantly higher RMSE, indicating worse model performance (Suppl. Figure S12D-G, Suppl. Table S2). These findings suggest that the presence of intronic seed sites is not sufficient on its own for ceRNA potential; rather, additional factors, such as chromatin retention or release dynamics (chromatin-association halftimes), and/or RNA-binding protein interactions [7], may be necessary for intronic ceRNA activity. Moreover, by comparing performance across all splicing-based gene expression models (mean and minimum RMSE across ten folds for genes passing the significance cutoffs, Methods), the PVT1 splicing-based model performed the best (Suppl. Figure S12D-G), establishing its validity in predicting expression of certain gene sets across tumor samples (see also Suppl. Note S2). Collectively, these results position PVT1, and potentially FTX, as key players in an expanded post-transcriptional regulatory landscape shaped by lncRNA splicing, intron retention, and chromatin tethering.

### Artificial modulation of PVT1 splicing uncovers regulation of target gene expression

The results from the in *silico* predictive models prompted us to validate the causality of PVT1 splicing activity in regulation of target gene expression via artificial splicing modulation in *vitro*. We used the CRISPR Artificial Splicing Factor (CASFx) technology [36, 37] to artificially enhance splicing at four 3’ splice sites across the PVT1 locus (ss11, ss17, ss25, ss33; Figure 7A, Suppl. Figure S13A). gRNAs were designed to target PVT1 RNA sequence just downstream of the adjacent exon of the targeted 3’ splice site, and gRNA blocks were constructed to accommodate recognition and binding of dCasRx-RBM25 splicing activator on target sites (Suppl. Figure S13D-E) [37].Constructs expressing gRNAs for all four splice sites were co-transfected in MCF-7 (Methods). After 48 h of transfection and dCasRx-RBM25 expression (Suppl. Figure S13D), we observed a change in the splicing profile of PVT1 captured by PCR on nuclear chromatin-released dT-primed RNA (Figure 7A-B, Suppl. Figure S13A). Across several possible alternative transcript isoforms (validated by Sanger sequencing of PCR products), a shared 586 bp tri-exonic splice junction was shifted to two alternative splicing events arising from 3’ splice site activation, an 890 bp derived from exon inclusion, and a 718 bp derived from concomitant exon inclusion and exon skipping (Figure 7A-B, Suppl. Figure S13A, S13F schematic representation). At the control condition, downstream of ‘common exon’, several alternative splicing-junction structures were validated (Figure 7A-B, Suppl. Figure S13A). Although the designated target 3’ splice sites were seemingly not activated, CASFx treatment suppressed all the alternative PVT1 splicing activity downstream of ‘common exon’ (perhaps by 3’ splice site activation in 3D-proximity), giving rise to one major splicing product (725 bp, encompassing ss32 and ss34; Figure 7A-B, Suppl. Figure S13A).

We presumed that CASFx-mediated enhanced splicing at a relatively few (certain) 3’ splice sites, (tending to resemble more (mRNA-like) constitutive splicing), shifting from alternative to constitutive splicing mode at the PVT1 locus, and an overall increase of splicing activity at the PVT1 locus would result in increased processing of spliced-out introns. Enhanced intronic turnover of spliced-out sequences could in turn mitigate a chromatin-tethered, intron-retained ceRNA activity attributed to the numerous miR-200 seed sites (perfect 7-mers or 8-mers) found exclusively within introns of PVT1 (Figure 7A, Suppl. Figure S1A-B). Indeed, we confirmed loss of intronic signal in chromatin-associated RNA (cDNA primed with random primers, Methods), for three out of four intronic amplicons (Figure 7C, Suppl. Figure S13B), and observed a decrease in RNA levels for two miR-200 target genes (GTF2I and SEC61B) checked by semi-quantitative PCR and qPCR (Figure 7D, Suppl. Figure S13C); a third miR-200 target gene remained unaltered (BIRC6), while VPS13B increased significantly (Suppl. Figure S14). Of note, loss of intronic signal was only observed in chromatin-associated RNA (random-primed cDNA), but not in nuclear chromatin-released RNA (dT-primed cDNA; Suppl. Figure S13C), suggesting that CASFx-mediated splicing enhancement is mostly co-transcriptional, and any chromatin-released intron-retained transcripts escape CASFx activity.

While aspiring for a genome-wide transcriptomic profiling, these results suggest that CASFx-mediated artificial splicing enhancement of PVT1 in *vitro* may bring about target gene expression changes in *trans*, potentially via modulating its chromatin-associated, intron-retained miR-200 ceRNA activity (Figure 7E, Model).

## DISCUSSION

Long non-coding RNAs have emerged as regulators of gene expression, acting through diverse mechanisms, including transcription activity from enhancers, RNA-binding protein interactions, chromatin association and miRNA sequestering (miRNA sponging or ‘competing endogenous’ ceRNA function). Yet, the mechanistic contribution of lncRNA splicing activity, particularly in relation to distal gene regulation and disease progression, remains to be defined. In this study, we frame PVT1 as a chromatin-associated lncRNA, whose splicing activity not only correlates with but may also causally shape genome-wide gene expression programs, including those enriched for miR-200/205 target genes, a pathway known to be dysregulated in epithelial-to-mesenchymal transition (EMT) and cancer metastasis [44, 46, 47, 63, 64].

Our analyses results support a model in which PVT1 splicing activity may function as a regulatory hub integrating signals from chromatin localization, intronic miRNA seed site content, and tumor-specific splicing profiles. First, machine learning models trained on PVT1 3’ splice site splicing efficiencies predicted gene expression in *trans*, with miR-200/205 targets significantly enriched among high-confidence predictions. This predictive power was not captured by PVT1 expression levels alone, and, in comparison with expression-based models trained on relative abundances of PVT1 alternative transcript isoforms, splicing-efficiency-based models performed with higher accuracy (Suppl. Fig. S9D). These results highlight the functional relevance of splicing beyond lncRNA abundance, and align with growing evidence that differential splicing and expression can underlie distinct biological pathways [54].

Second, we showed that distinct 3’ splice sites of PVT1 contribute non-redundantly to gene regulation, with k-means clustering revealing splice site-specific gene expression modules. These modules varied in their miRNA target-gene content and functional annotation, suggesting that i) splicing activity at certain splice sites may relate with differential intronic sequence turnover/metabolism and miRNA availability (due to the presence of perfect intronic miR-200 seed sites), and ii) it may give rise to or promote the expression of alternative PVT1 isoforms that may exert isoform-specific regulatory effects. Nonetheless, that the splicing efficiency per *se* did not always (or weakly) correlated with alternative transcript isoform abundance, and a small overlap between the splicing-based and alternative isoform-based elastic net model predicted genes and functional annotations, re-aligned with the notion that expression and splicing may underlie distinct biological pathways [54]. Moreover, the relationships between PVT1 splicing activity and gene expression were to some extent subtype-specific, with Basal and Her2 showing PVT1 splicing heterogeneity and prognostic relevance. These observations support a model in which splicing dynamics of chromatin-associated lncRNAs like PVT1 are not static but remodeled across tumor subtypes, possibly reflecting or driving subtype-specific gene regulatory programs.

Another recent work defined cancer subtypes based on splicing changes rather than gene expression alone (by employing an unsupervised Bayesian-based framework for discovery of splicing-based subtypes, capturing structured splicing variation across patient cohorts [65]), underscoring the relevance and emerging power of splicing-based or splicing-centered modeling in refining tumor classification and predicting clinical outcomes. While our study differs being mechanistically anchored on a single lncRNA locus (as a framed paradigm), our finding that PVT1 splicing patterns stratify breast cancer subtypes, predict gene expression patterns and are prognostic of clinical outcomes (Suppl. Fig. S6), supports a broader statement that splicing provides valuable information beyond mutation- or expression-based stratification.

Third, we validated the causal role of PVT1 splicing in regulating distal gene expression using tumor-specific SNVs as natural perturbations. This approach uncovered specific splice sites (ss13, ss15) whose altered splicing due to nearby SNVs led to consistent expression changes in downstream sets of target genes, which were enriched for miR-200 targets. Notably, these same splice sites had been prioritized by elastic net regression and clustering, underpinning their central role in shaping regulatory potential of PVT1. This tumor-specific somatic perturbation-informed causal inference framework provides support for a mechanistic model where local splicing events at the PVT1 locus transmit regulatory effects to distal targets, plausibly via (chromatin-associated) intron-retained ceRNA activity.

Fourth, we demonstrated that CASFx-mediated splicing enhancement of PVT1 in *vitro* alters its splicing profile and reduces chromatin-associated intron retention, accompanied by changes in expression of selected miR-200 target genes. This result, although limited in scope, offers experimental support for our proposed intronic ceRNA-based post-transcriptional regulatory model, where splicing refines intron availability for miRNA binding (Figure 7E). It also underlines the context-dependence of PVT1 conferred intronic ceRNA activity, which likely requires chromatin retention, inefficient co-transcriptional processing, and low intronic turnover rates (i.e. a relatively high intronic stability).

Our findings resonate with previous observations that chromatin-retained or chromatin-associated lncRNAs are enriched for regulatory potential, particularly in nuclear processes such as splicing/RNA processing, chromatin organization or modification [10, 12, 66]. We extend this view by framing splicing activity as a quantifiable and predictive layer of lncRNA functionality, suggesting that the splicing machinery may act as a gatekeeper not just for transcript maturation, but for broader post-transcriptional regulatory networks involving miRNAs, with their levels buffered by chromatin-associated intron-retained ceRNA activity.

Our cross-tissue generalization analysis suggests that PVT1 splicing-based gene expression models trained in breast cancer are most predictive in prostate cancer, suggesting that shared post-transcriptional gene regulatory programs, e.g. those involving translation and RNA processing, may allow cross-tissue model applicability (Suppl. Note S3; [67]). Such observations may hold promise for translational applications, such as building tissue-specific or pan-cancer biomarkers based on lncRNA splicing profiles.

Nonetheless, our study has some limitations. We show that PVT1 lncRNA splicing activity is predictive of target gene expression in tumor condition, but to what degree this applies in healthy tissue remains unresolved. We note here that our attempts to uncover PVT1 splicing quantitative trait loci (cis-sQTL) and putative relationships to gene expression effects in *trans* (eQTL), using germline mutations (SNVs) from 234 normal samples (healthy tissues) available on TCGA (Suppl. Fig. S1D, Suppl. Methods), did not yield any genetic variants that would affect both splicing patterns of PVT1 (cis-sQTL) and expression of distal genes (trans-eQTL). This could be because a higher number of samples should be analyzed, highlighting the need to explore additional datasets.

Currently, the mechanism by which chromatin-associated, intron-retained ceRNA activity occurs at the PVT1 locus requires further experimental validation. In our model (Figure 7E), we presume that the numerous chromatin-associated intronic miR-200 seed sites (perfect 7-mers) sequester mature miRNAs in association with nuclear AGO, keeping it in a ‘poised’ state. Beyond the well-characterized cytoplasmic miRNA functions, emerging studies uncover the presence of mature miRNAs in the nucleus, and shed light on potential nuclear functions of miRNAs, highlighting co- and post-transcriptional roles in gene regulation, even on nascent RNA [61, 68–70]. Of note, by examining available CLIP-seq data of AGO in MCF-7 cells [71], we found several intronic binding sites, some of which near miR-200 seed sites (Figure 7A and Suppl. Figure S13A). We presume that upon artificial splicing enhancement with CASFx in *vitro,* spliced-out intronic sequences undergo turnover by nuclear exonucleases [72]; in addition, upon splicing enhancement, chromatin-released intronic sequences could become available substrates for AGO-interacting effector proteins [73]. This may result in the observed intronic turnover (Fig. 7C) and an increased effective availability of miR-200-AGO for target gene modulation (Fig. 7D-E).

In summary, we frame PVT1 as a chromatin-associated lncRNA whose splicing dynamics affect genome-wide gene regulation, partially through miRNA-dependent ceRNA mechanisms relying on nuclear intron retention. Our integrative framework combining splicing quantification, machine learning, causal inference, and experimental perturbation contributes to a functional splicing-centric view of lncRNA biology in cancer and beyond, and may inform the development of novel biomarkers and therapeutic targets based on splicing-modulated (chromatin-associated/intron-retained) nuclear regulatory hubs.

## METHODS

### TCGA BRCA Controlled Data

Authorized access to TCGA controlled data was granted to EN through signed agreement (research project #35033). To define our cohort, we applied a series of filters through the TCGA data portal.

In cohort builder, in the General tab, we selected samples from the “*TCGA-BRCA”* project with a disease type classified as “*ductal and lobular neoplasms”* and a primary diagnosis of “*infiltrating duct carcinoma, NOS”*, with Tissue of origin being *breast, NOS.* Under Demographic, we limited the cohort to Gender *female*, Race “*asian”, “white”, “black or african american”,* and *“not reported”*, with a Vital Status of “*dead”* or “*alive”.* We excluded individuals with Prior malignancy or Prior Treatment (*“no”*) under the Disease Status and History tab. For Biospecimen characteristics, we chose Tissue Type “*tumor”* (“*normal”* for control), Specimen Type to be “*solid tissue” (“solid tissue normal”* for control) *and* Tumor Descriptor *“primary”* keeping the “*ffpe”* Preservation method. Finally, from the Available Data tab, only samples generated via Experimental Strategy *RNA-Seq* with a Workflow Type of *“STAR 2-Pass Genome”* were included. We downloaded the bam controlled files and ended up with 696 BRCA and 80 matched control samples, for a total of 776.

For samples used in generalization:

i) Prostate (*TCGA-PRAD*); only samples of Specimen Type “*solid tissue*” without Prior malignancy, Prior Treatment or Synchronous Malignancy were selected
ii) Ovary (*TCGA-OV*); Primary Diagnosis “*serous cystadenocarcinoma, nos”,* Disease Type *“cystic, mucinous and serous neoplasms”* of Tumor Grade “*g3*” and *“g2”,* Tissue or Organ of Origin and Primary Site “*ovary*”, without Prior Treatment and Synchronous Malignancy *“not reported”* and *“no”*
iii) Uterus (*TCGA-UCEC*); Primary Diagnosis *“endometrioid adenocarcinoma, nos”*, Disease Type *“adenomas and adenocarcinomas”* and *“cystic, mucinous and serous neoplasms”,* Primary Diagnosis *“endometrioid adenocarcinoma, nos”,* Tissue or Organ of Origin *“endometrium”* and Primary Site *“corpus uteri”*
iv) Testis (*TCGA-TGCT*); Primary Diagnosis *“embryonal carcinoma, nos”, “seminoma, nos”, “mixed germ cell tumor”* and *“not reported”,* Disease Type *“germ cell neoplasms”* Tissue or Organ of Origin *“testis, nos”*
v) Adrenal gland (*TCGA-ACC*); Primary Diagnosis *“adrenal cortical carcinoma”*, Tissue or Organ of Origin *“cortex of adrenal gland”*

Race, Tissue Type (*“tumor”*), Experimental Strategy and Workflow Type for all generalization tissues were the same as in TCGA-BRCA filters.

### BAM preprocessing

The 776 BAM files were sorted (chromosomal coordinates) and re-indexed. To streamline downstream processing, for each sample, reads were extracted from a defined genomic region (CHROM:START-END -- chr8:127784541-128197101 for PVT1) before converting to BED format, using the *-cigar* flag, which retains the CIGAR string annotation and was consequently used to identify spliced alignments (GENCODE v44 comprehensive gene annotation; GRCh38.p14). Reads were filtered to keep only uniquely mapped alignments (mapping quality = 255), from chromosomes (1–22, X, Y). TCGA RNA-Seq reads are unstranded (GDC Documentation - mRNA Analysis Pipeline) and while the majority originated from paired-end sequencing, we identified exceptions of single-end and treated them accordingly. Finally, reads were partitioned into split (containing ‘N’ in the CIGAR string) and non-split reads, to later quantify splicing efficiency (*cluster_SE.sh*).

### Splicing efficiency values at PVT1 3’ splice sites

Using the GENCODE V44 comprehensive gene annotation (standard chromosomes; GRCh38.p14), we extracted exon entries of PVT1 and splice site coordinates for 3’ and 5’ splice sites. For 5′ splice sites (donor), a 2 bp window was defined spanning the last nucleotide of the exon and the first nucleotide of the intron. Conversely, for 3′ splice sites (acceptor), the 2 bp window is spanning the last nucleotide of the intron and the first of the adjusted exon. Only canonical 5’ and 3’ splice sites were kept having GT and AG at the two first and two last intronic nucleotides, respectively (extracted with *bedtools getfasta* - hg38.fa -s). Unique entries were extracted by collapsing coordinates using *bedtools groupby* [74]. Splicing efficiency (*θ* value) was computed as before ([7, 27, 75]). Split and non-split read coverage spanning unique splice site coordinates was computed using *bedtools coverage* -f 1. We kept splice sites with a minimum total read coverage (split + non-split) of 10. Splicing efficiency (*θ* value) was calculated as the ratio of split to total (split + non-split) read counts (*get_splicing_eff.py*).

### Splice site (feature) selection

We extracted 3′ splice site coordinates for the PVT1 gene from GENCODE v44 (Comprehensive) and retained only canonical AG-bearing sites. Splicing efficiency was quantified across 776 BRCA samples (696 tumor and 80 control) by summing split and non-split read coverage at each splice site, and extracting the ratio of split to total (split + non-split) reads (using *get_splicing_eff.py*). Sites with fewer than 10 total reads in a sample were excluded. The number of detectable splice sites varied per sample, with a median of 50. To ensure a consistent input feature set, we excluded samples with fewer than 50 splice sites and proceeded with a splice site selection process (*parse_transform_multi.py*).

Starting with the sample containing the highest number of splice sites (maximum of 53), we built an initial feature list and then iteratively included additional samples (sorted in descending order by splice site count). At each step, we retained only those splice sites that were common to all samples considered so far. A user-defined threshold (‘*individual_threshold*’) was applied to exclude any sample that would reduce the retained set of splice sites below this threshold. This process resulted in a final set of 34 shared 3′ splice sites across 670 samples (617 tumor and 53 control). A similar strategy was used for 5′ splice sites, yielding a final set of 61 sites across 620 samples (575 tumor and 45 control).

For other assayed gene candidates, where lower expression or variability in splicing activity was observed (e.g., SMYD3, Suppl. Fig. S11A), a modified feature selection approach was used. Specifically, samples were included only if they preserved a minimum number of shared splice sites for each gene, defined by a gene-specific cutoff (‘*drop_cut*’). Rather than imputing missing values—which could introduce noise due to the narrow range of splicing efficiency scores—we excluded samples with insufficient splice site coverage.

The final script used was *parse_transform_multi.py*.

### Linear Regression Models - Regularized linear regression (elastic nets)

We used regularized linear regression to determine the most important subset of features (splices sites) for prediction of target gene expression, with soft constraints on non-zero coefficients. This helps assigning similar weights for correlated variables, which was particularly useful given the presence of few sets of correlated features in our data (Figure S3A). We used Elastic Nets [76] as implemented in the *glmnet* package for R [52]. By including all exon-level raw counts (genome-wide/all genes) for the 776 BRCA cohort we removed any genes that were not expressed in any of samples and filtered lowly expressed genes using *filterByExpr* of the package *edgeR* [77], ending up with 30,887 genes (this becomes the ‘response’ variable in our models).

We kept only samples included in our final splice-site defined cohort (53 control and 617 tumor) and log-transformed the raw read counts (see Methods paragraph Quantifying genome-wide gene expression).

Based on the number of samples, we performed 10 x cross-validation per gene when samples exceeded n ≥ 500 (otherwise 5 x cross-validation was performed). An extra internal filtering step was performed (per gene) to exclude potential folds with non-informative response values (i.e. when the response variable exhibited very low or zero variance; *refine_cv_linear.r*). Splicing efficiency (SE) values at 34 PVT1 3’ splice sites were used as predictor variables. Elastic Net models were trained using an *α* of 0.5 to balance L1 (Lasso) and L2 (Ridge) penalties using the *glmnet* package [52] in R. For each fold, the optimal regularization parameter (*λ*) was selected using internal cross-validation of *glmnet* (*cv.glmnet(x_train, y_train, alpha=0.5, type.measure=“mse”, family=“gaussian”)*) and the selected *λ* (lambda.min) was then used to fit the final fold-specific model (*glmnet(x_train,y_train,alpha=0.5, lambda=lam)*) from which we calculated the root mean squared error (RMSE; equation 2) and the raw coefficient of determination (R^2^; equation 1).

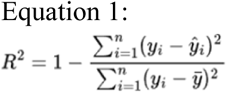

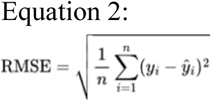

After each fold was processed we identified the fold with the lowest RMSE as the “best fold” and the following metrics were saved: i) the mean of R^2^ and RMSE from all folds (mean_raw_r_sq, mean_rmse), ii) best fold metrics (best_fold_R^2^, min_RMSE) iii) correlation test results between R^2^ and RMSE from all folds (Pearson correlation value, correlation p-value) and iv) best fold’s model coefficients. We considered significant models those that passed three filtering criteria; a) best_fold_R^2^ ≥ 0.2 b) correlation value R^2^ with RMSE < 0 c) all_fold_mean_RMSE < Q3 + 1.5*IQR, which yielded 365 genes.

### Linear regression models – lm

The same pre-processing steps and cross-validation steps were applied for the multivariate linear regression models using the *lm* function in R) to predict gene expression with 34 variables (splicing efficiency values at the 34 PVT1 3’ splice sites). We calculated and reported the following metrics: i) the mean of *lm*() quantified ”Adjusted R-squared” from all folds and the correlation between observed and predicted values (all_fold_mean_adjR^2^, all_fold_mean_Pearson_corr), ii) best fold performance metrics (best_fold_adjR^2^, best_fold_corr), iii) coefficient estimates with the associated lm quantified significance level Pr(>|t|) (*refine_cv_linear.r*). We defined significant models using mean adjusted R^2^ ≥ 0.2 and mean correlation > 0.2 resulting in a total of 316 genes predicted across all Tumor samples.

### Random Forest classification

Random Forest (RF) classification models were built to examine the contribution of individual PVT1 splice sites (34 splicing efficiency values) or PVT1 Salmon-quantified alternative splicing isoform transcript expression (19 features, tag per million -TPM expression values) to breast cancer subtype classification, with the *randomForest* R package. Five classification models were examined (LumA versus non-LumA, LumB versus non-LumB, Her2 versus non-Her2, Basal versus non-Basal and LumA versus LumB). The number of *trees* (*ntree* parameter) was fixed to 1000, while the number of variables for each tree (*mtry* parameter) was optimized using the *train()* function of the *caret* R package with a grid search in a ten-fold cross validation on the training data. The best *mtry* was selected, such that it maximizes the model accuracy. Model training was repeated ten times and each time 10% of the data was left out for testing, while the other 90% was used for model optimization, as described above. The importance of each feature was computed as ‘mean decrease in Importance index’, ‘mean decrease in Accuracy index’ and ‘mean decrease in Gini index’. Variables with large positive importance values correspond to features which are crucial for model classification, whereas variables with values close to zero or negative correspond to unimportant features.

### K-means clustering

K-means clustering was performed with R function *kmeans ()*, with a defined k the number at which the total within-cluster sum of squares reached a minimum plateau, and *nstart*=100

### Survival Analysis

Survival analysis was performed as in [78].

Clinical metadata for BRCA was downloaded from TCGA, which contained OS (overall survival), disease-specific survival (DSS), disease-free interval (DFI), progression-free interval (PFI) for 610 subtyped samples. Survival was assessed across all tumor samples and within each subtype. We stratified samples based on the median value of either the splicing efficiency at each of the 34 PVT1 3’ splice sites (cutoff SE ≥ median and < median; Suppl. Fig. S6; *BRCA_survival.R*), or Salmon-quantified transcript expression (tags per million, TPM) for each of the 19 top (most variable, Suppl. Fig. S7B-C) PVT1 isoforms (cutoff TPM > median and TPM ≤ median; Suppl. Fig. S8; *BRCA_SalTPMsurvival.R*).

We used the R package *survival* [79, 80]. Kaplan–Meier (KM) curves were fitted with the *survfit* function, and we assessed the significance of the curves using the log-rank test in *survdiff*. P-values were derived from the chi-squared distribution with *k*-1 degrees of freedom, where *k* is the number of groups (*k*=2 since we defined the 2 groups as “Above Median” and “Below Median”).

*Surv(time = survival_groups[[metric_info$time]], event = survival_groups[[metric_info$event]])*

*survfit(surv_object ∼ Q_group, data = survival_groups)*

*survdiff(surv_object ∼ Q_group, data = survival_groups)*

*pval = 1 - pchisq(survdiff_res$chisq, df = length(survdiff_res$n) - 1)*

*ggsurvplot(km_fit, data = survival_groups, color = “Q_group”, pval = TRUE, censor = TRUE, risk.table = TRUE, surv.median.line = “hv”)*

### Causal Inference Analysis

For modeling the *SNV* - *splicing* - *putative target gene expression* relationships, we ran causal inference analysis using PVT1 splice site-proximal tumor-specific SNVs.

To test whether changes in PVT1 splicing efficiency causally affect distal gene expression, we utilized tumor-specific single nucleotide variants (SNVs) from TCGA BRCA WXS and WGS data as in *silico* perturbations. From the 617 BRCA tumor samples analyzed, 592 were genotyped either by WXS or WGS, and 410 of those harbored at least one SNV in the PVT1 locus.

Each SNV was mapped to its nearest PVT1 3′ splice site with *bedtools closest*, resulting in a binary annotation per splice site (1 = SNV present near site; 0 = no SNV). The remaining 182 samples had no SNVs in the PVT1 locus and were assigned ‘0’ across all splice sites. To increase resolution, we split the samples into groups bearing mutations associated with either splicing upregulation or downregulation, achieving statistical significance in mutation-associated changes in splicing efficiency for most splice sites (Figure 5C, Suppl. Fig. 10A). For each of the 34 PVT1 3’ splice sites, we constructed a dataset with 592 rows (samples) and three columns (variables):

SNV: binary indicator (0 or 1) of SNV proximity to the given splice site

SE: splicing efficiency of that splice site (continuous; range 0–1)

Expression: transcript-level (exon_counts) RPM values of a putative regulated gene.

Using these data, and running the script in a loop for all 30,887 genes with measured RPM expression across all 592 BRCA samples, we defined and tested eight Bayesian network models (Figure 5A):

Causal: SNV → Splicing → Expression

*model2network*(“[snv][splicing|snv][genexp|splicing]”)

Reactive: SNV → Expression → Splicing

*model2network*(“[snv][genexp|snv][splicing|genexp]”)

Independent: SNV affects splicing and expression independently: SNV → Expression, SNV → Splicing

*model2network*(“[snv][splicing|snv][genexp|snv]”)

Model 4: SNV affects only splicing, gene expression varies independently: SNV → Splicing, [Expression]

*model2network*(“[snv][splicing|snv][genexp]”)

Model 5: SNV affects only gene expression, splicing varies independently: SNV → Expression, [Splicing]

*model2network*(“[snv][genexp|snv][splicing]”)

Model 6: Null model

*model2network*(“[snv][splicing][genexp]”)

Model 7: Gene expression affects splicing independently of SNV

*model2network*(“[snv][genexp][splicing|genexp]”)

Model 8: Splicing affects gene expression independently of SNV (i.e. to test whether PVT1 splicing independently explains gene expression variability)

*model2network*(“[snv][splicing][genexp|splicing]”)

Bayesian network fitting and model selection were performed using the R package *bnlearn* [60]. For each *SNV* – *splice site* – *gene* triplet, we computed posterior probabilities for all eight models. Cases where the Causal model exceeded a defined threshold 0.9 were selected for downstream analysis.

### Generalization analysis on unseen data

To assess the generalizability of our BRCA-trained splicing-expression models, we selected five TCGA tissue types where PVT1 expression is comparable to BRCA (Suppl. Fig. S10A; n number of samples): Ovary (n = 393; 341 post-filtering), Prostate (n = 431; 300), Uterus (n = 502; 83), Testis (n = 144; 43), and Adrenal Gland (n = 79; 38). For each sample, we extracted (i) splicing efficiency values at the 34 selected PVT1 3’ splice sites and (ii) genome-wide exon-level gene expression values, processed identically to BRCA (see Methods: Splicing efficiency values at PVT1 3’ splice sites; Quantifying genome-wide gene expression). Genes were filtered to match the 30,887-gene BRCA set.

Only samples with splicing efficiency values for all 34 splice sites were retained (*parse_transform_multi.py*, single_selected mode). BRCA-trained elastic net coefficients (from the best fold of the 10-fold cross-validation) were applied to predict gene expression in each tissue. For every gene and tissue, we computed predicted expression, coefficient of determination (R^2^), and root mean squared error (RMSE) (*refine_generalize_linearCV.r*).

We then compared the performance of the original BRCA models (minimum fold-RMSE and best fold-R^2^) with their performance in each non-BRCA tissue using Pearson correlation test (*cor.test() in* R). Correlations were evaluated across: (i) all genes in each tissue (prostate in Suppl. Fig. S10C; not-shown but values reported in Results for the rest), (ii) the 365-gene BRCA subset (Fig. 6B–F), and (iii) subsets with R^2^ > 0.2 or R^2^ > 0.1 (Suppl. Fig. S11C).

### Gene expression quantification

Gene-level expression was quantified using *featureCounts* [81] with GENCODE v44 comprehensive annotations. BAM files were processed according to library type (single- or paired-end; *cluster_refineFeat_final.sh*). Exon-level counts were obtained for each gene (-t exon -g gene_id) using uniquely mapped reads (-Q 255).

Raw counts were filtered to exclude non-expressed or lowly expressed genes. Specifically, genes not expressed in any sample were removed, followed by exclusion of genes with raw counts <10 in fewer than 10 samples. For construction of the final BRCA gene set (30,887 genes), only the first filtering step was applied (using edgeR::*filterByExpr*).

Counts were normalized to Reads Per Million (RPM), winsorized at the 99th percentile to reduce the impact of outliers, and log-transformed using a pseudo-count of 1e−9. For elastic net regression models predicting transcript expression from PVT1 3’ splicing (190 alternative transcript isoforms with Salmon quantified TPM), a higher pseudo-count of 1e−5 was used prior to log transformation.

### Enrichment Analysis of Gene Ontology Terms

Gene ontology (GO) enrichment analysis was performed using the clusterProfiler package [82] in R, with gene annotations provided by the org.Hs.eg.db library. Analyses were conducted using the “SYMBOL” or “ENSEMBL” gene identifiers, depending on the input list, and ontologies were set to “ALL” (inclusive of BP, MF, and CC) or separately to “BP” (Biological Process), “MF” (Molecular Function), or “CC” (Cellular Component), as appropriate. Enriched terms were computed as follows:

ego <-enrichGO(gene = genes, OrgDb = org.Hs.eg.db, keyType = “SYMBOL”, ont = “ALL”, pAdjustMethod = “BH”)

barplot(ego, showCategory = 20)

In Figure panels Fig. 5G–H, enrichment analysis results were visualized with dotplot and cnetplot [83] using GO <-enrichGO(gene = geneid$gene, keyType = “ENSEMBL”, OrgDb = org.Hs.eg.db, ont = c(“BP”, “MF”, “CC”), pAdjustMethod = “fdr”, pvalueCutoff = 0.05, qvalueCutoff = 0.2, minGSSize = 5, readable = TRUE) dotplot(GO)

cnetplot(GO, categorySize = 10)

Downstream visualizations and filtering were performed using tissue_GO_analysis.R. Adjusted p-values (FDR) were used for term prioritization. Only significantly enriched terms were retained for plotting.

**PAM50 molecular subtyping** on BRCA tumor samples using TCGA RNA-seq data

Raw exon-level counts from 696 TCGA-BRCA tumor samples (n = 39,938 genes) were filtered to remove non-expressed genes. Counts were normalized using *edgeR* () [77], applying trimmed mean of M-values (TMM) scaling and log₂-transformation of counts per million (CPM). Ensembl gene identifiers (with version suffix) were mapped to HGNC gene symbols using biomaRt v2.56 [84]. Genes without one-to-one matches were discarded, and for symbols mapping to multiple Ensembl IDs, the instance with the highest expression was retained, resulting in 24,267 unique gene symbols. Among the PAM50 genes, 46 of 50 were present; missing genes included CDCA1, KNTC2, MIA, and ORC6L. Molecular subtypes were assigned using molecular.subtyping() from the *genefu* R package [85], with no additional mapping required. This resulted in 263 LumA, 182 LumB, 87 Her2, 155 Basal, and 9 ‘Normal’ samples. For consistency with the splicing analysis (see Methods, “Splice site feature selection”), 617 tumor samples were retained (LumA: 235, LumB: 178, Her2: 62, Basal: 135).

### Enrichment analysis of miR-200 and miR-205 putative target genes

We compiled a list of putative target genes for the miR-200 family using miRPathDB 2.0 [51], querying “hsa-mir-200a/b/c,” “hsa-mir-141,” and “hsa-mir-429”, and retained only genes with experimental validation (“experimental (all)” and “experimental (strong)”). This resulted in 1,136 unique putative targets.

Additional 1,916 miR-200 family targets were curated from five previously published studies [45, 47, 48, 50]. After resolving gene name redundancy and converting all entries to Ensembl stable gene IDs (based on GENCODE v44 Comprehensive annotation), this yielded a non-redundant set of 2,356 miR-200 family target genes.

We also included 26 validated targets for hsa-miR-205-5p and -3p from miRPathDB, resulting in a final combined set of 2,382 unique putative target genes (2,356 miR-200 and 26 miR-205 targets).

To evaluate whether these targets were enriched in gene sets of interest, we applied Fisher’s exact test (Suppl. Table S2; *check_enrich.R*).

miRNA target gene lists are available at /splicing_pvt1_brca/data/mirnaTG_initial_lists/ within the associated GitHub repository.

Experimental procedures

### Cell Culture

MCF-7 cells cultured in Dulbecco’s Modified Eagle’s Medium (DMEM, Gibco, 41966052) supplemented with 5% fetal bovine serum (FBS, Gibco, A5256701), glutamine and sodium pyruvate.

### Fractionation of MCF-7 cells and RNA extraction

Cell fractionation was done as in [14] resulting in chromatin-associated and nuclear chromatin-released RNA samples. Briefly, cells were scraped in PBS and lysed with 400 uL lysis buffer (10mM Tris pH 7.5, 150 mM NaCl, 0.15% NP-40). The lysates were layered on top of 2.5 volumes of 24 % sucrose buffer and precipitated for 8 min at 5000 g, resulting in nuclei precipitation. The cytoplasmic fraction (supernatant) was discarded, and nuclei were washed with 1ml ice-cold PBS-EDTA and resuspended in 250 uL 50 % Glycerol Buffer. An equal volume of nuclear lysis buffer (10 mM Hepes pH 7.6, 7.5 mM MgCl2, 0.2 mM EDTA, 0.3 mM NaCl, 1 M Urea, 1 NP-40) was added and samples were incubated on ice for 5 min. After brief vortexing, nuclear lysates were precipitated at 14000 g for 2 min at 4 °C, resulting in chromatin fraction (pellet) and nucleoplasmic fraction (supernatant). The pelleted chromatin fraction was washed with ice-cold PBS-EDTA and resuspended in 15 nM Tris-HCL (pH 7.0), followed by mechanical disruption using a 21-gauge needle attached to a syringe. RNA was extracted from all fractions with acidic Phenol (pH 4.5) and CHCl3, followed by precipitation with 3.5-4 volumes 100% EtOH and 1/10 volume NaAc (pH 5.2).

### cDNA synthesis and PCR

cDNA synthesis was done from 150 ng of chromatin-associated RNA using random primers, and 400 ng of nuclear chromatin-released RNA using a dT20VN+adapter primer (AP_RT_3’RACE), according to standard protocol (NEB, M0368). Intronic turnover was checked with PCR (NEB, M0273) on cDNA, and target gene expression levels were checked with both semi-quantitative and qPCR (Applied Biosystems). All primer sequences are listed in Table 3.

### Artificial splicing modulation with CASFx

gRNA design was done according to published method [37], using Cas13design [86] or TIGER [87], resulting in 2 gRNAs for each target splice site. To generate gblocks containing 2 gRNA sequences and a 36-nt direct repeat between them, the sequence was broken down to short oligos (IDT) duplexed in a way to create sticky ends, which were then phosphorylated (NEB, M0201) and ligated (NEB, M0202), to be combined into a single double-stranded molecule. The resulting ligation products were then amplified by PCR (NEB, M0491) and ligated into a gRNA expressing plasmid (gRNA-CasRx_SD1-pSV40-TagBFP, Addgene #221004). All gRNA and oligo sequences are listed in Table 2.

**Table 1.** PVT1 34 3’ splice sites, plus strand coordinates (hg38)

|  |  |  |  |
| --- | --- | --- | --- |
| chr8 | 127803149 | 127803151 | ss1 |
| chr8 | 127816827 | 127816829 | ss2 |
| chr8 | 127818899 | 127818901 | ss3 |
| chr8 | 127819908 | 127819910 | ss4 |
| chr8 | 127851933 | 127851935 | ss5 |
| chr8 | 127854727 | 127854729 | ss6 |
| chr8 | 127855154 | 127855156 | ss7 |
| chr8 | 127855162 | 127855164 | ss8 |
| chr8 | 127881800 | 127881802 | ss9 |
| chr8 | 127882061 | 127882063 | ss10 |
| chr8 | 127890588 | 127890590 | ss11 |
| chr8 | 127902455 | 127902457 | ss12 |
| chr8 | 127932027 | 127932029 | ss13 |
| chr8 | 127932052 | 127932054 | ss14 |
| chr8 | 127932464 | 127932466 | ss15 |
| chr8 | 127989161 | 127989163 | ss16 |
| chr8 | 127996560 | 127996562 | ss17 |
| chr8 | 128031057 | 128031059 | ss18 |
| chr8 | 128031958 | 128031960 | ss19 |
| chr8 | 128038916 | 128038918 | ss20 |
| chr8 | 128048447 | 128048449 | ss21 |
| chr8 | 128049584 | 128049586 | ss22 |
| chr8 | 128049587 | 128049589 | ss23 |
| chr8 | 128069068 | 128069070 | ss24 |
| chr8 | 128070159 | 128070161 | ss25 |
| chr8 | 128070161 | 128070163 | ss26 |
| chr8 | 128074069 | 128074071 | ss27 |
| chr8 | 128074072 | 128074074 | ss28 |
| chr8 | 128080302 | 128080304 | ss29 |
| chr8 | 128082752 | 128082754 | ss30 |
| chr8 | 128096517 | 128096519 | ss31 |
| chr8 | 128096520 | 128096522 | ss32 |
| chr8 | 128099369 | 128099371 | ss33 |
| chr8 | 128100979 | 128100981 | ss34 |

**Table 2.**
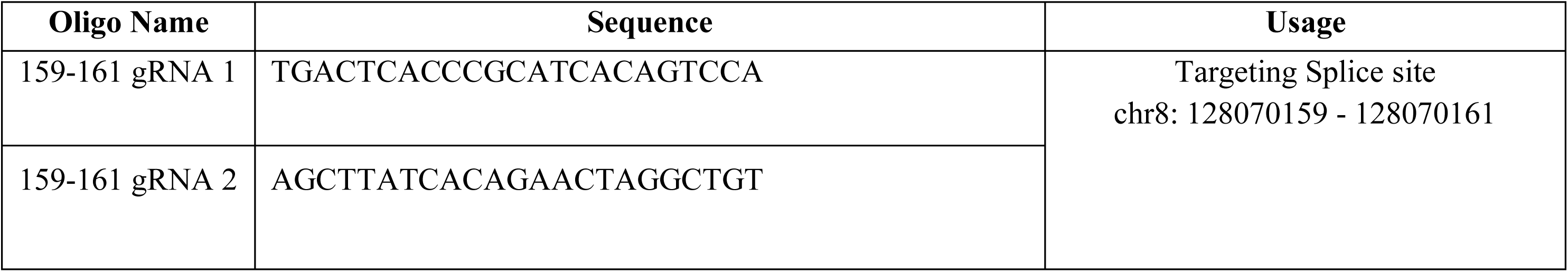
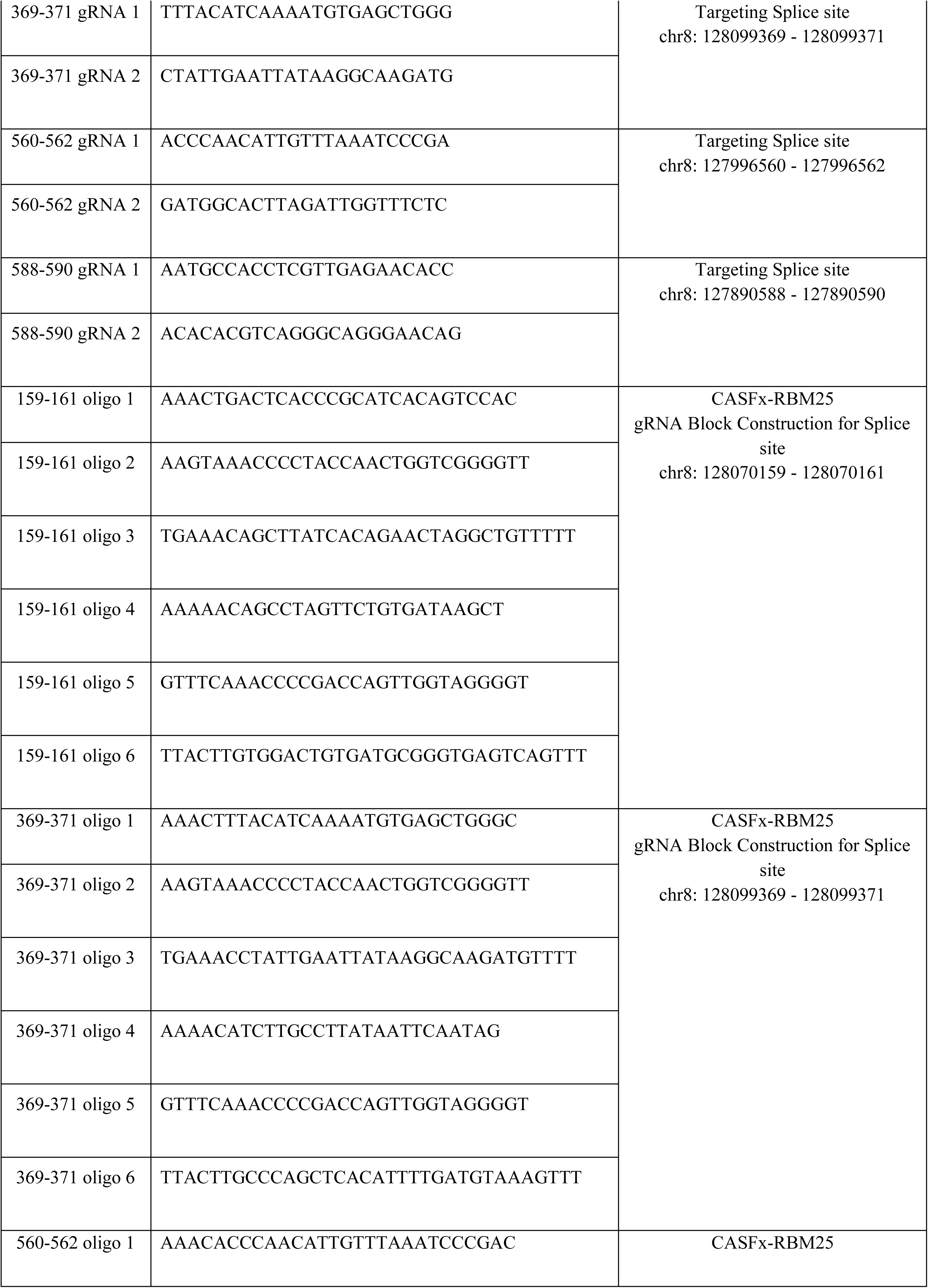
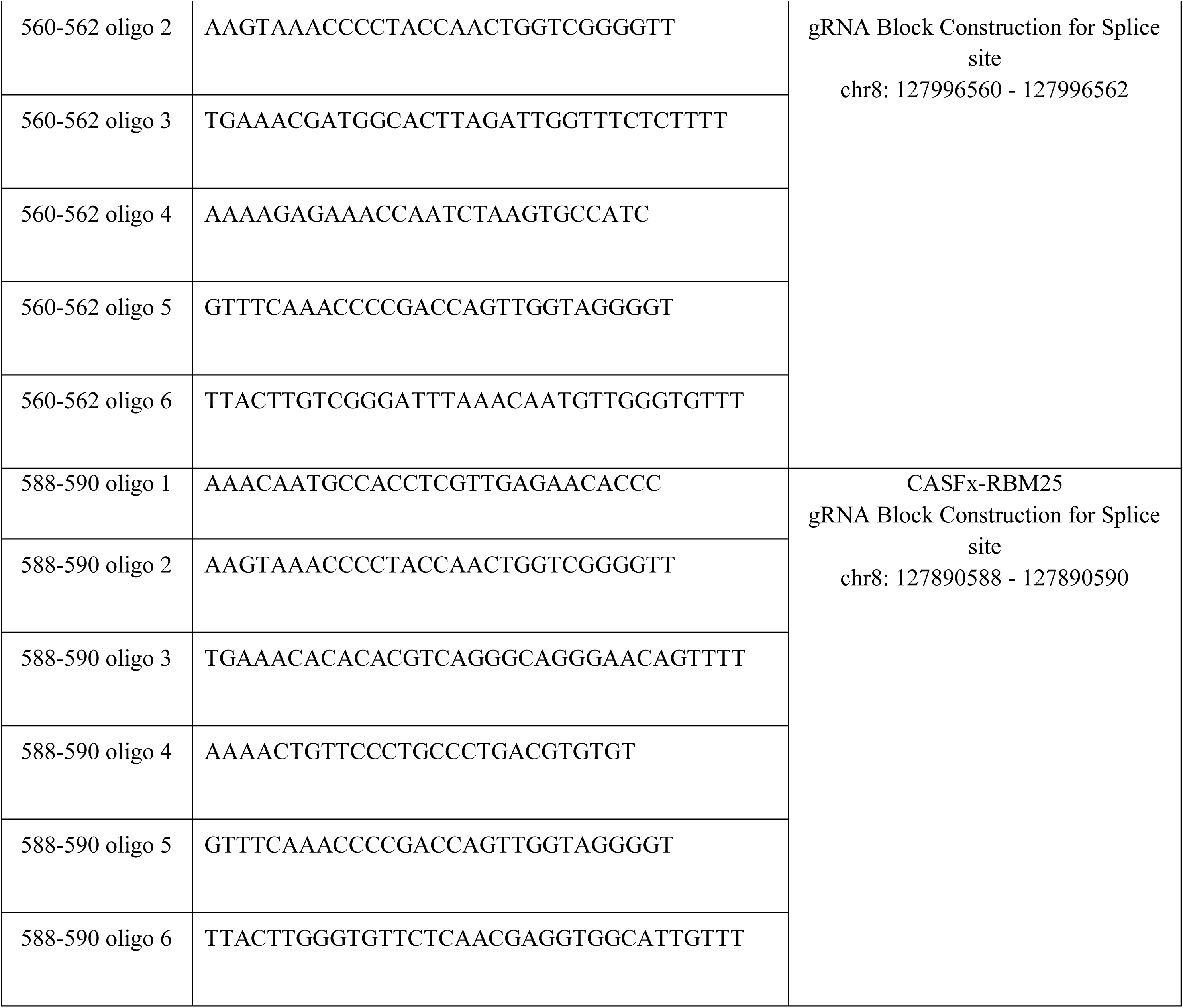
gRNA and DNA Oligo Table.

**Table 3.**
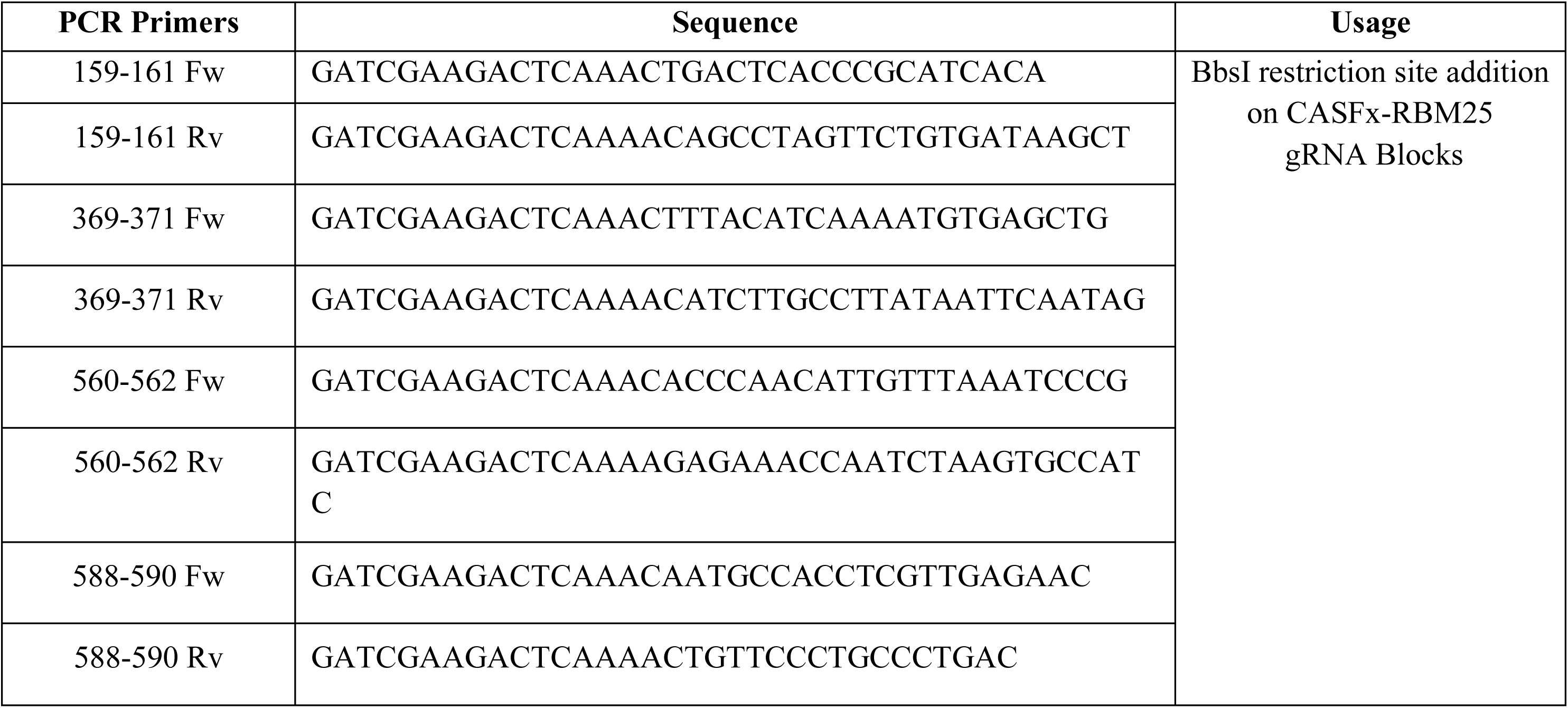
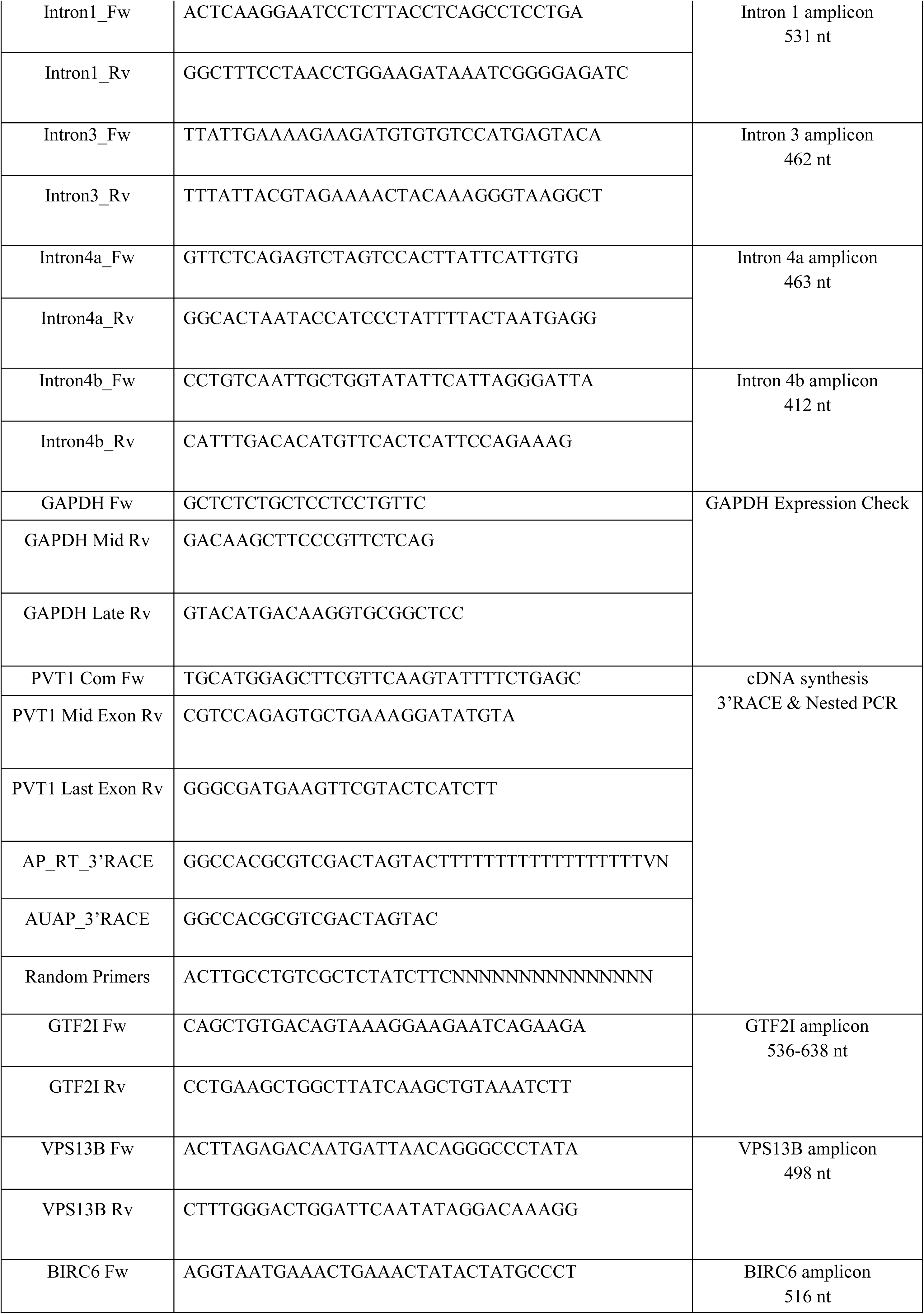
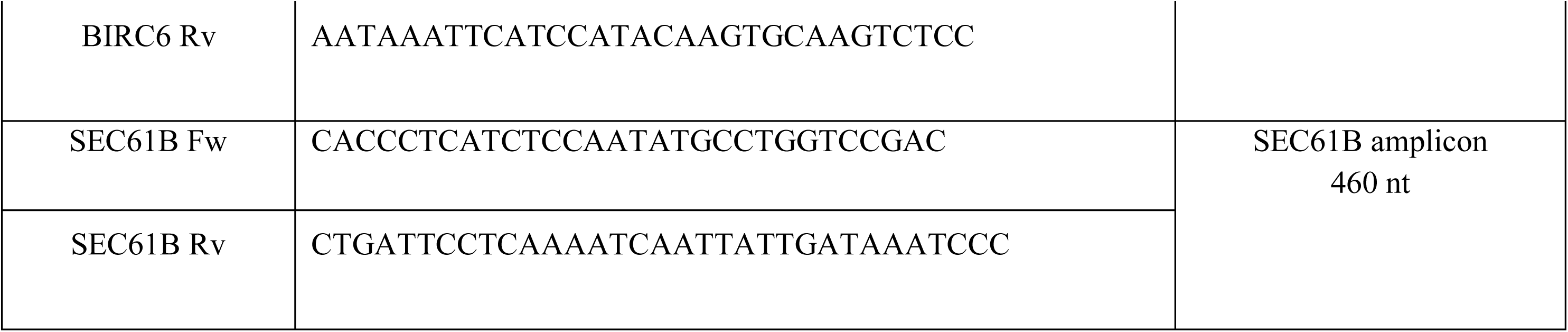
PCR primers.

MCF-7 cells were transfected at 70 % confluency in 100 mm plates using jetPEI (Polyplus, 101000053) with 5 ug EF1a-dCasRx-RBM25 (Addgene #221001) and 5 ug of plasmids expressing gRNAs targeting 4 different splice sites (1.25ug for each plasmid). Transfection was repeated after 24 hours only with the gRNA expressing plasmids (5 ug total) and cells were harvested 24 hours after the second transfection and fractionated, resulting in acquiring chromatin-associated and nuclear chromatin-released RNA samples. Concentration of RNA was measured by Nanodrop.

### 3’ RACE and Transcript Variants Validation

cDNA from nuclear chromatin-released RNA was used as template for 3’RACE PCR with “PVT1 Com Fw” and “AUAP_3’RACE” primers according to NEB’s protocol (NEB, M0273). Then, nested PCR reactions were performed with “PVT1 Com Fw” and “PVT1 Mid Exon Rv (R4)” or “PVT1 Last Exon Rv (R10)” (NEB, M0273) and got analysed by agarose gel electrophoresis (1%). Single bands were excised, and DNA was extracted with Monarch Spin DNA Gel Extraction Kit (NEB, T1120). Validation of amplicon sequences was done via Sanger sequencing by Genewiz (Azenta Life Sciences), with the primers used to acquire each band. All primer sequences are listed in Table 3.

### RNA-seq from the chromatin-released nuclear RNA fraction

MCF-7 cells at 70 % confluency were treated with 50 uM cisplatin (Tocris, 2251) for 48 hours before harvesting. Control and cisplatin-treated cells were fractionated, resulting in acquiring nucleoplasmic (nuclear chromatin-released) RNA samples. The concentration of RNA was measured with Qubit and 1.5 ug of chromatin-released RNA was used as input for polyA+ enrichment (NEBNext® Poly(A) mRNA Magnetic Isolation Module, E7490). Library construction was performed with NEBNext® Ultra II Directional RNA Library Prep Kit for Illumina (NEB, E7760), NEBNext® Multiplex Oligos for Illumina® (NEB, E6440) and NextSeq 1000/2000 P3 XLEAP SBS-Reagent kit (Illumina, 20100989) flow cell was used for the sequencing on an Illumina NEXTSEQ2000 sequencer.

Reads were aligned to hg38 with STAR [88], sam alignments were converted to bam files with *samtools* [89]and bam to bed with *bedtools bamtobed -split*. Only uniquely mapped reads were kept (filtering for score 255), and strand-specific reads were converted to wig (with normalization scale reads per million), and strand-specific bigwig files with *wigToBigWig* from the UCSC utilities.

### Data Availability

NIH authorized access to TCGA data was granted to EN (research project #35033).

Ago2 HITS-CLIP peaks in MCF7 were downloaded from GSM2065792 (GSE78059) [71] and coordinates were lifted over to hg38 using *liftover* from UCSC utilities.

Raw RNA-seq data from MCF-7 and MCF-10 whole-cell total RNA were retrieved from GSE71862 [90] and re-processed to generate strand-specific bigwig files for visualization. Strand-specific bigwig files of chromatin-associated nascent RNA-seq in MCF-7 were from GSE218726 [7], and of total nucleoplasmic RNA-seq from GSE110149 [14].

RNA-seq data from chromatin-released polyA+ selected nuclear RNA (random-primed cDNA) in MCF-7 (conditions control and 50 mM cisplatin 48 hr treatment), and strand-specific bigwig files for visualization have been deposited on GEO accession nr xxx.

### Code Availability

All code is available on github https://github.com/AngelosKoz/splicing_pvt1_brca

## Figure Legends

**Supplementary Note S1** (related to Figure 6 and Suppl. Figure S11)

RMSE and R^2^ quantify distinct aspects of model performance. R^2^ measures the fraction of expression variance explained by the predictors and therefore depends on the total sum of squares (TSS) of the response variable. When expression variance is small, as we observed for many genes outside BRCA, TSS approaches the residual sum of squares (RSS) and R^2^ collapses toward zero irrespective of prediction accuracy. In contrast, RMSE reflects the absolute magnitude of prediction errors and is insensitive to the scale of the response.

Consistent with this property, RMSE correlations between BRCA and every external tissue were uniformly high (p-value < 1e-16 for all Pearson tests), whereas the corresponding R^2^ correlations were weak or null (sometimes slightly negative) except in the Prostate. The pattern held across different gene-sets definitions: full dataset (all screened genes), 365 genes predicted at high confidence in BRCA, and gene sets passing tissue-specific R^2 filters.

This indicates that the BRCA-derived models transfer their error structure captured by RMSE to other tissues, but they fail when it comes to the proportion of variance explained, which is modulated by the variance of the target gene in the new context. Accordingly, RMSE is the more informative metric for judging cross-tissue portability of our Elastic Net (glmnet) models. Still, further improvements and fine tuning is required for cross-tissue generalization.

**Supplementary Note S2** (related to Suppl. Figure S12)

When building splicing-based gene expression models using elastic nets, a combination of sample number where the model is trained and number of features (variables, i.e. number of splice sites) likely affects the performance. Across all nine splicing-based gene expression models compared, the three that rendered worse performance (higher mean and best-fold (minimum) RMSE) were trained in less than 200 samples. There, we could not reach a higher number of samples because we did not detect common splicing activity (splice sites) amongst samples, thus, to include more splice sites and have a meaningful feature set, we excluded samples where less splice sites were used (see also Methods paragraph ‘Splice site (feature) selection’ and *parse_transform_multi.py* on github). It is expected that a higher number of training samples and available features (variables) contribute to more stable splicing-based gene expression models (Suppl. Fig. S12 D-G).

**Supplementary Note S3** (related to Figure 6 and Suppl. Figure S11)

A described key commonality between the prostate and breast, absent in the ovary, adrenal gland, and uterus, is their strong dependence on sex hormones (androgens and estrogens) for normal development and function, making them particularly susceptible to hormone-related cancers. This shared hormonal dependence may shape their transcriptional programs and influence cancer development [67].

## Acknowledgments

E.N. acknowledges a Fondation Santé research grant. C.K. acknowledges an H.F.R.I. PhD fellowship.

## Author contributions

A.K. did bioinformatics work, computational analyses, interpreted data. C.K. performed experiments. E.N. conceived the study, supervised research, interpreted data and wrote the manuscript with input from A.K. All authors have read and approved the manuscript.

