## Supplementary Figures S1-S14 for "PVT1 splicing activity predicts genome-wide gene expression with miRNA regulatory signatures"

### Suppl. Figure S1

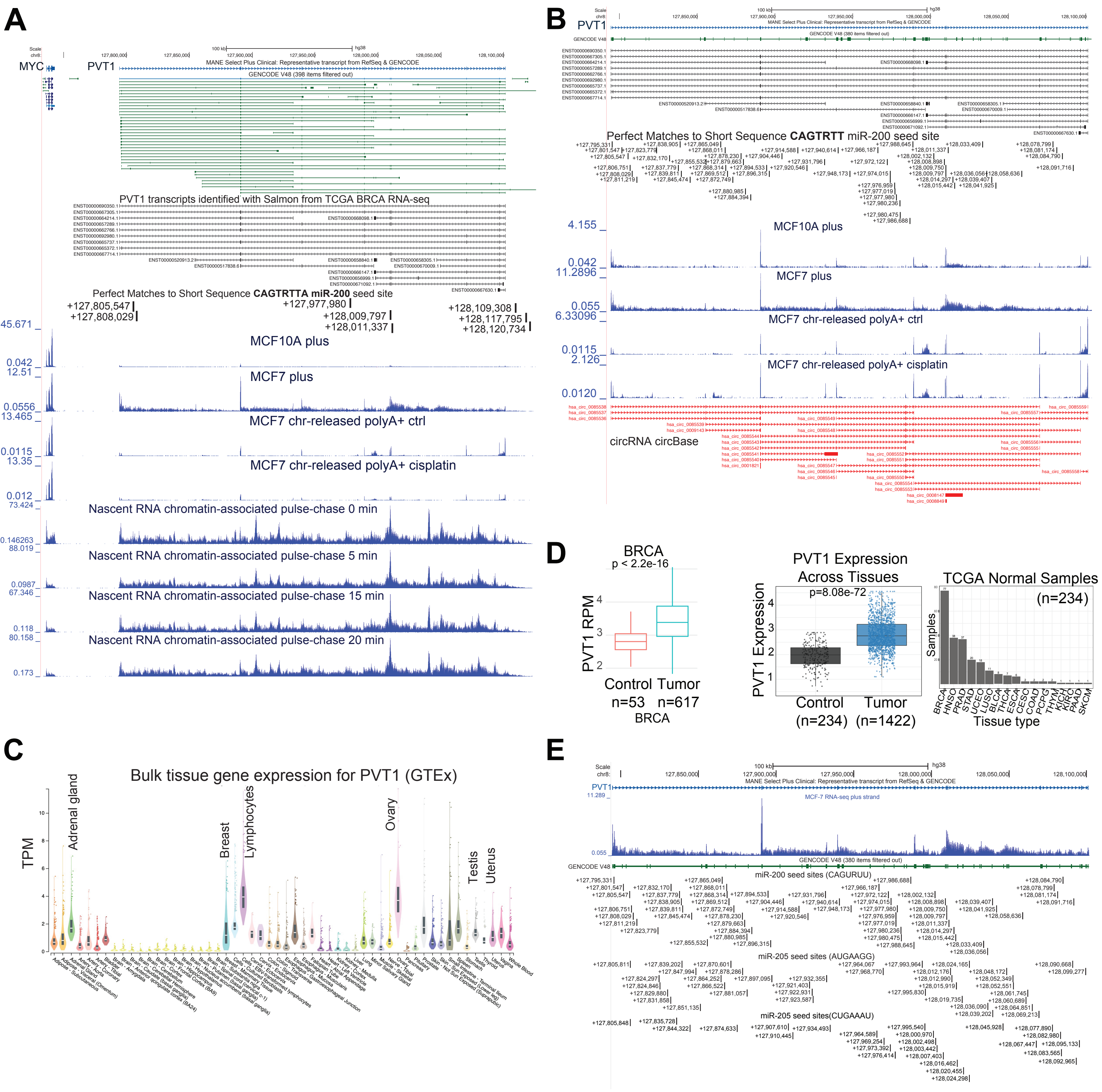

### Suppl. Figure S2

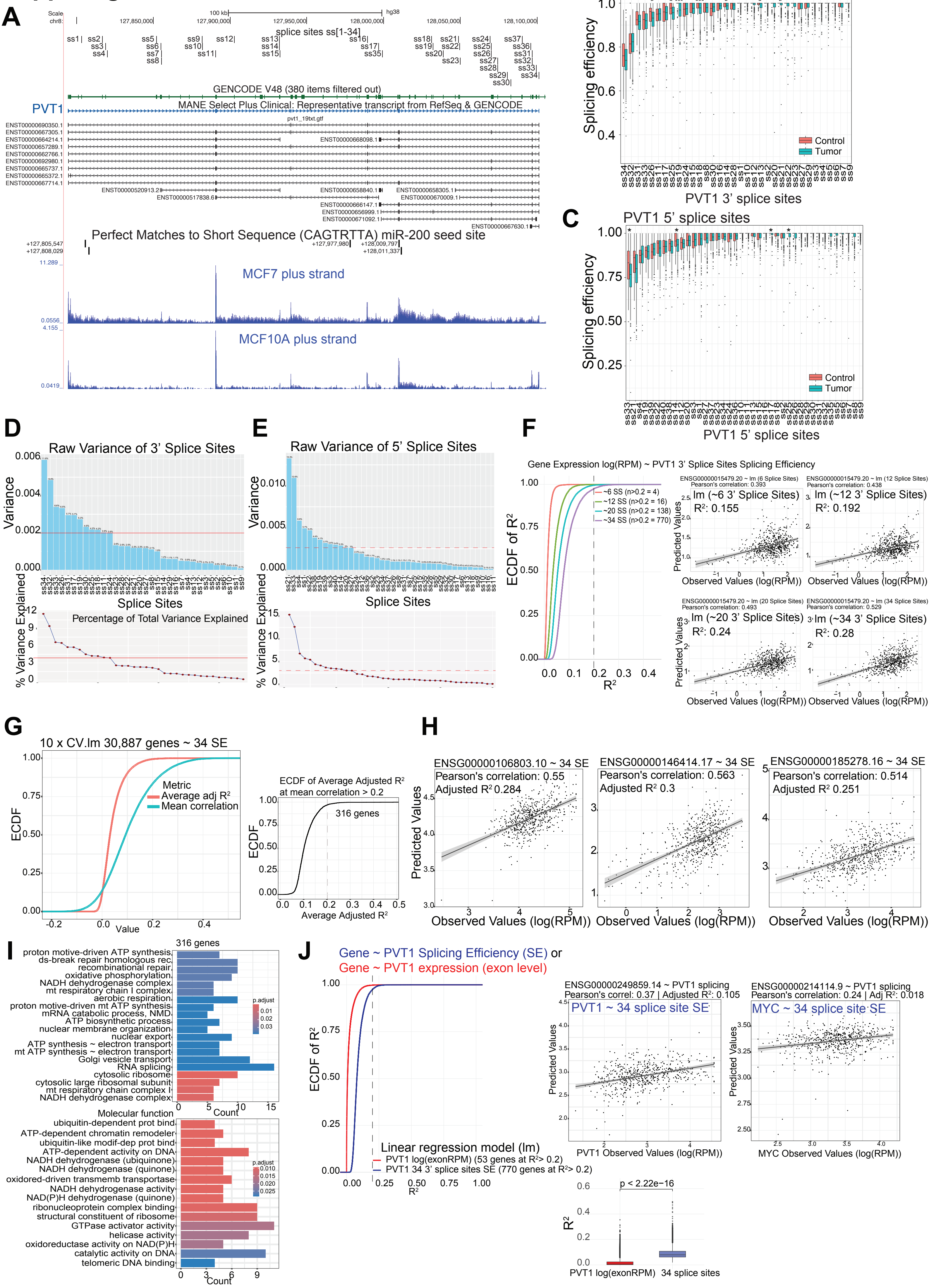

**Suppl. Figure S3**

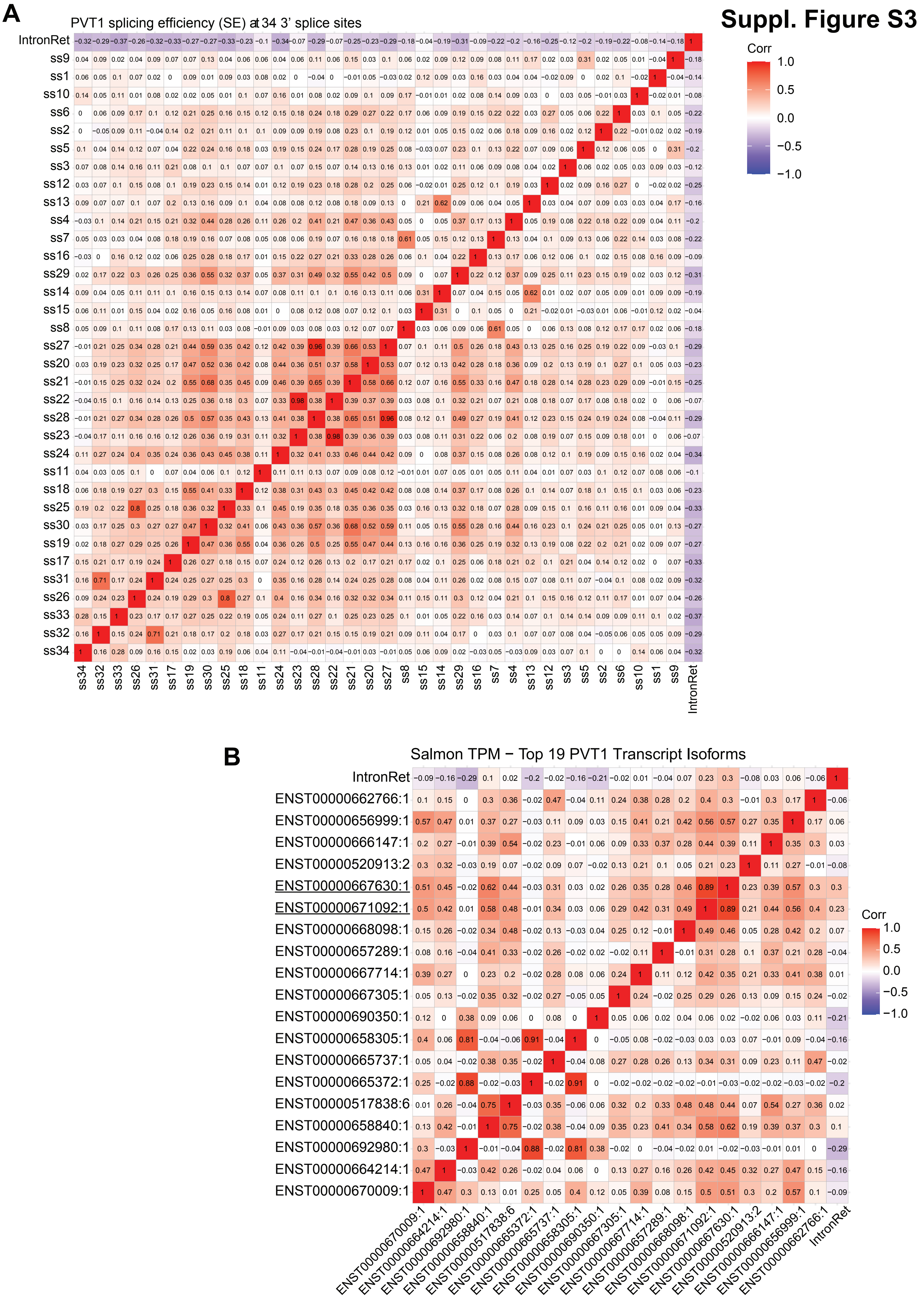

**C**

#### Suppl. Figure S3 C

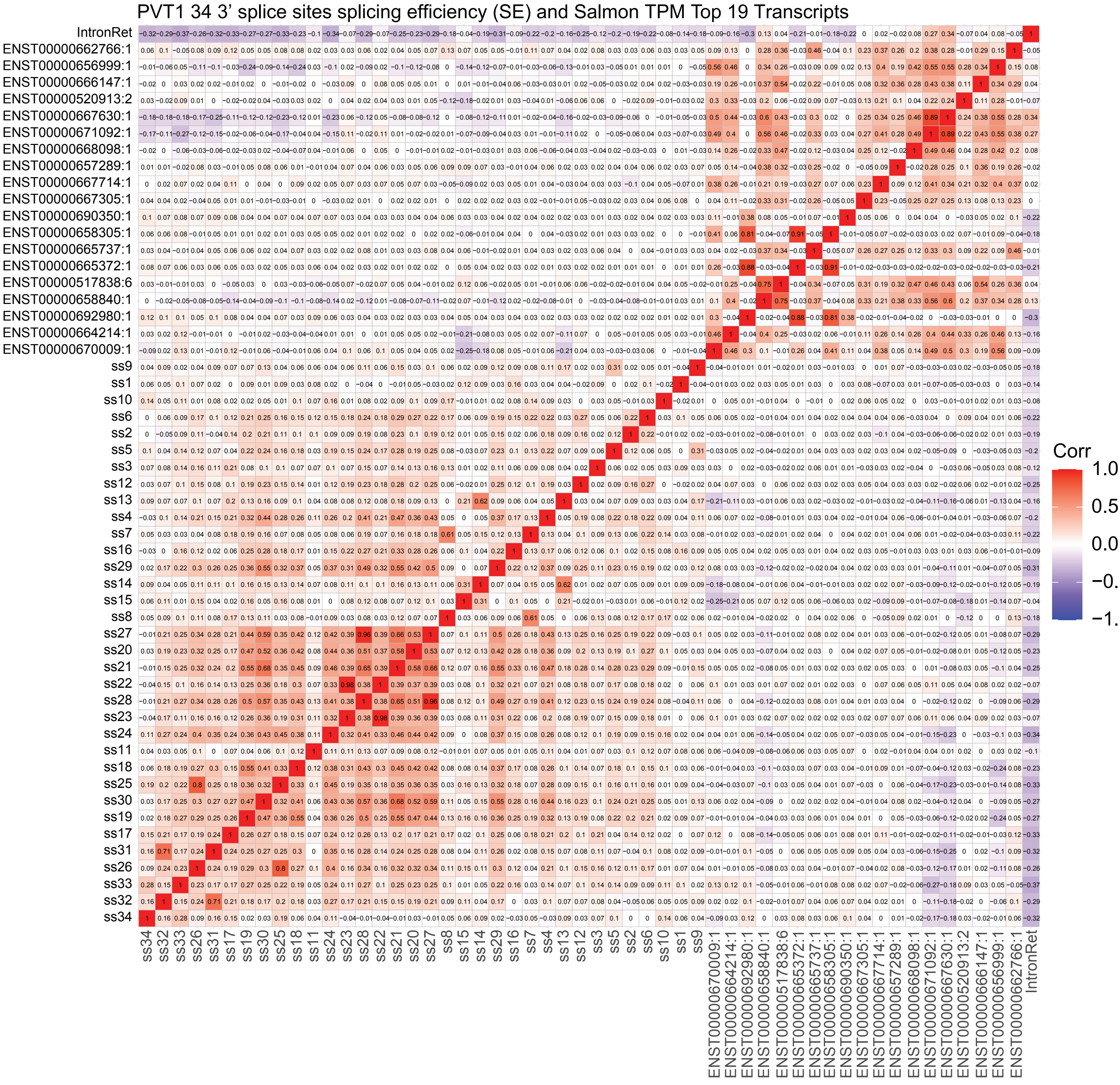

### Supplementary Figure S4

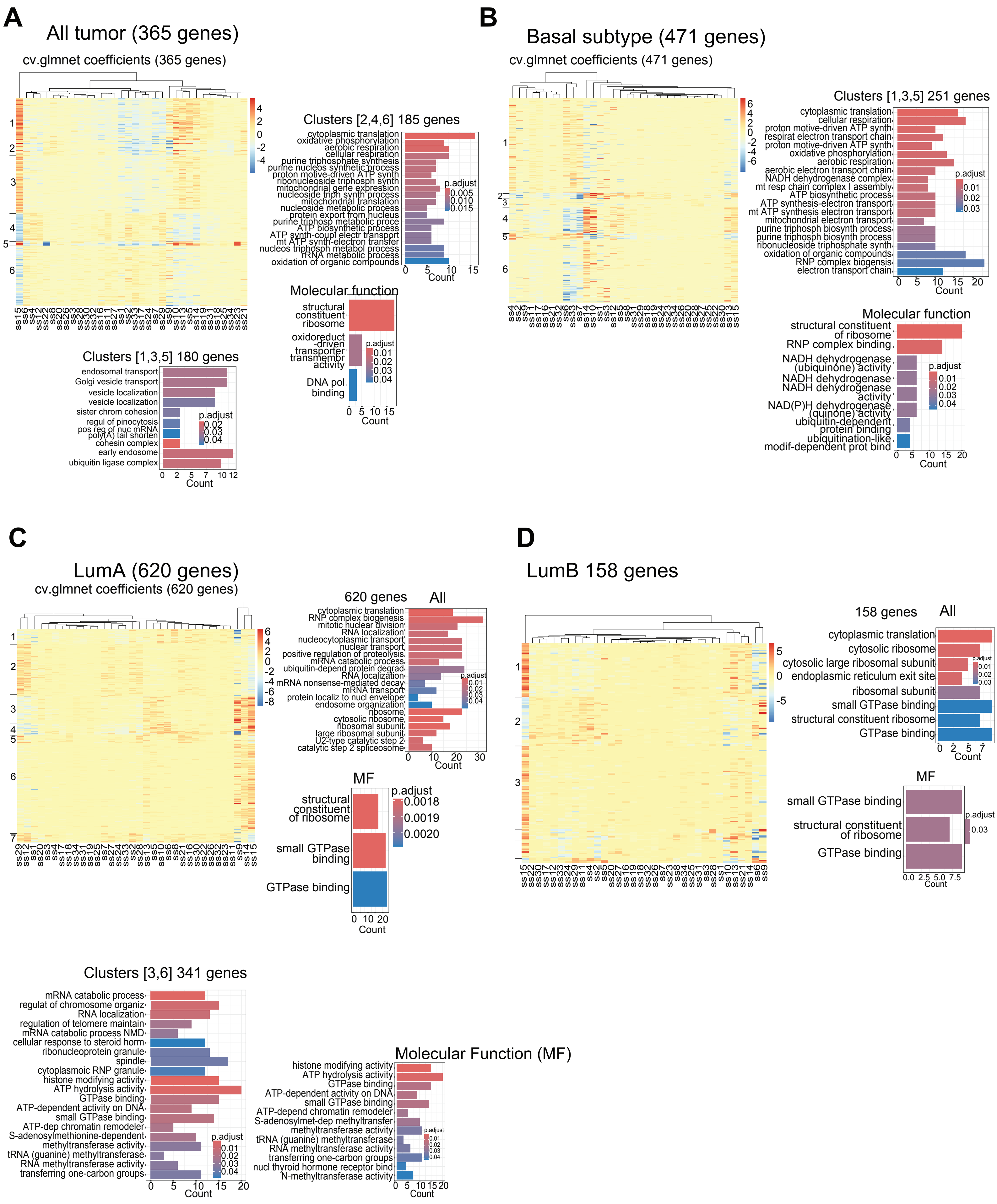

### Suppl. Figure S5

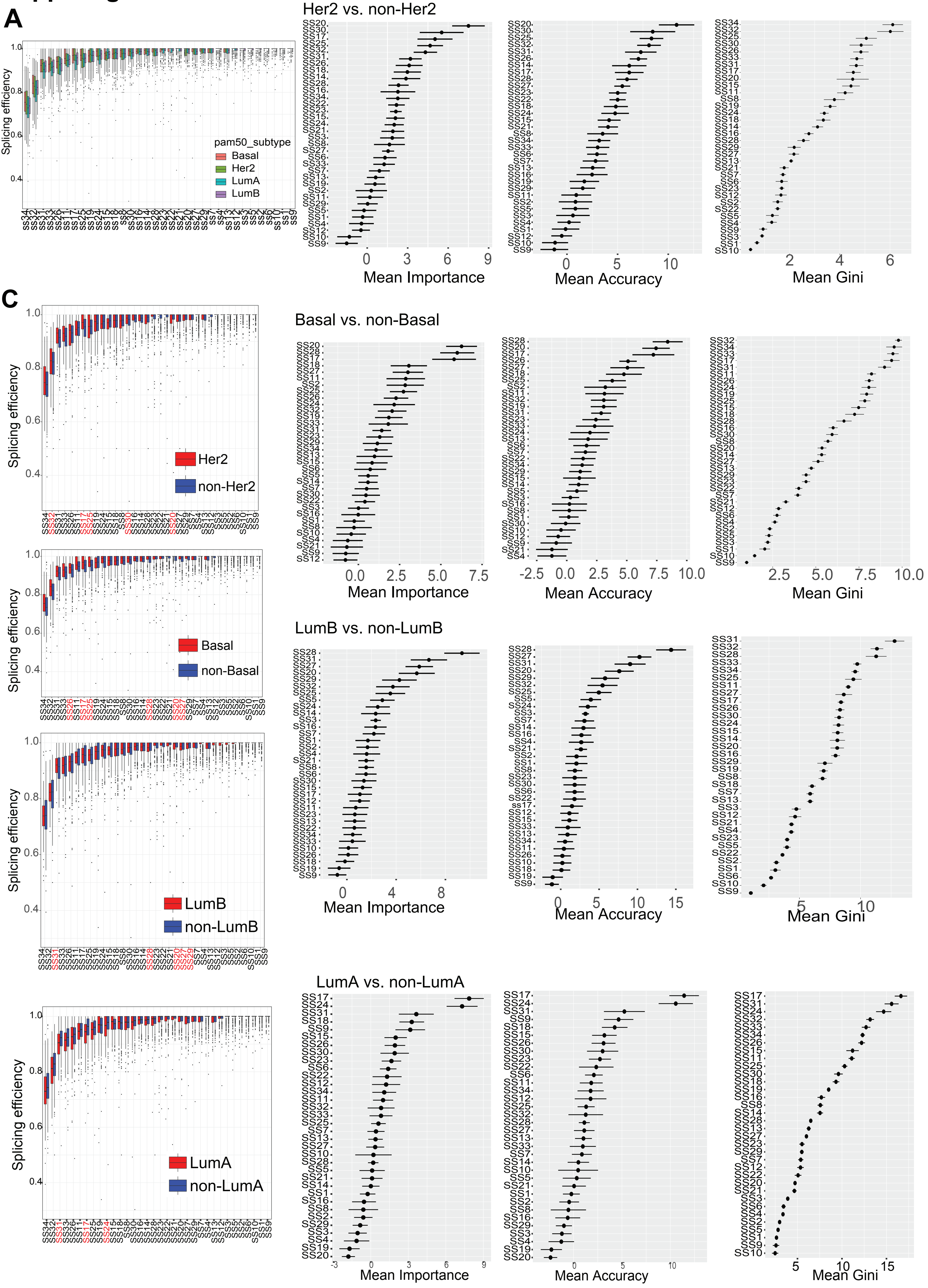

#### Suppl. Figure S6

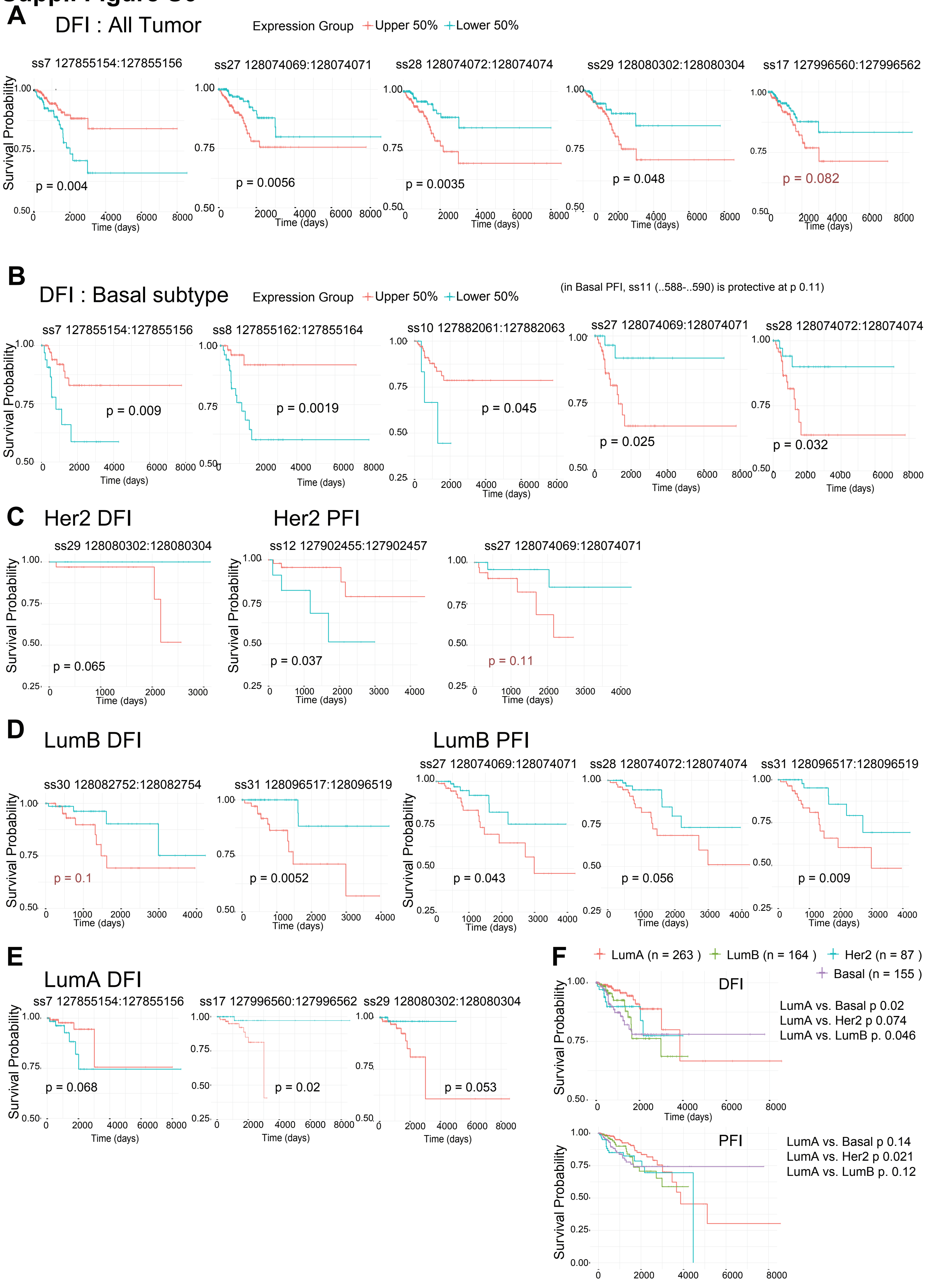

### Suppl. Figure S7

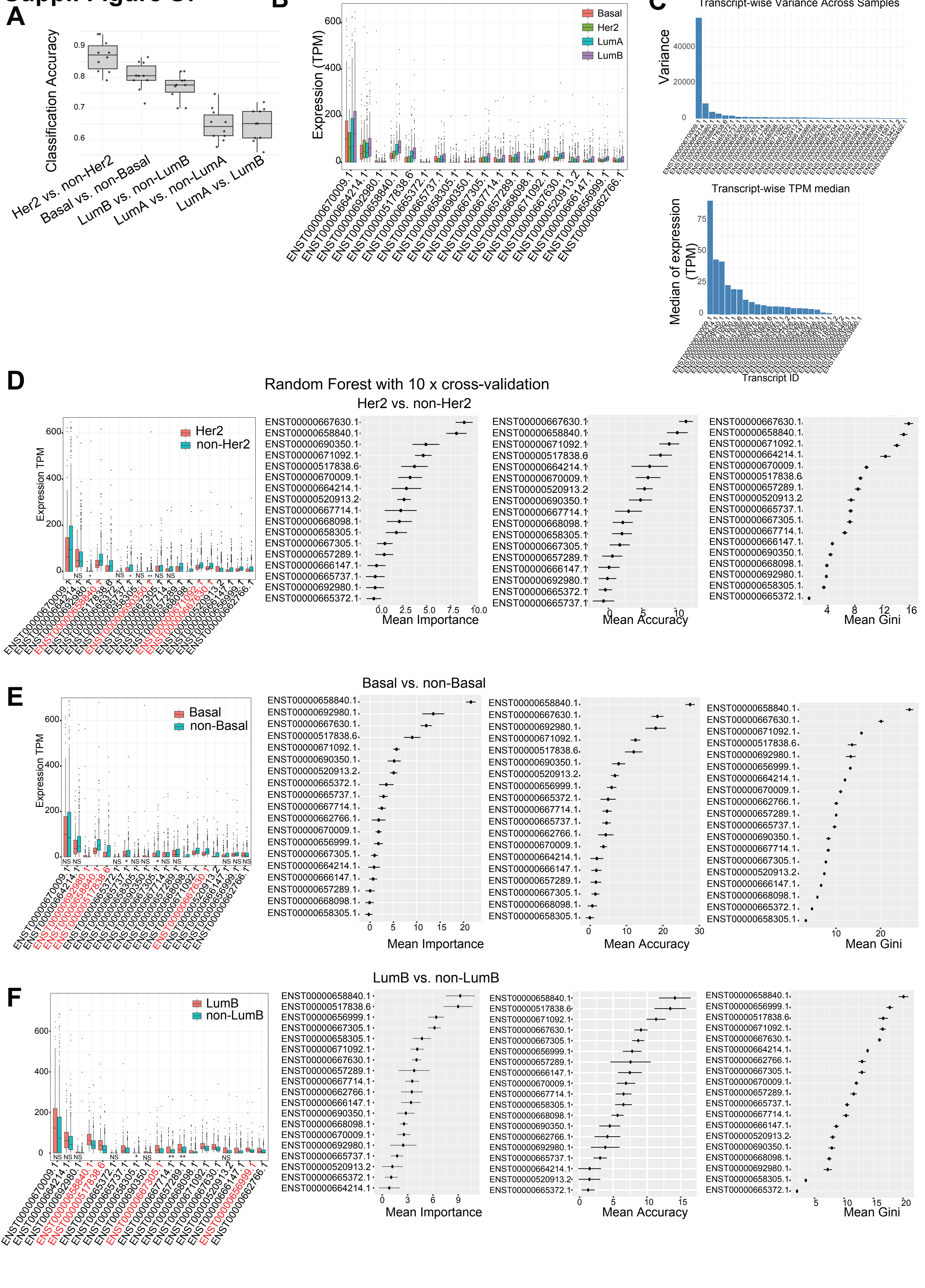

**Suppl. Figure S8**

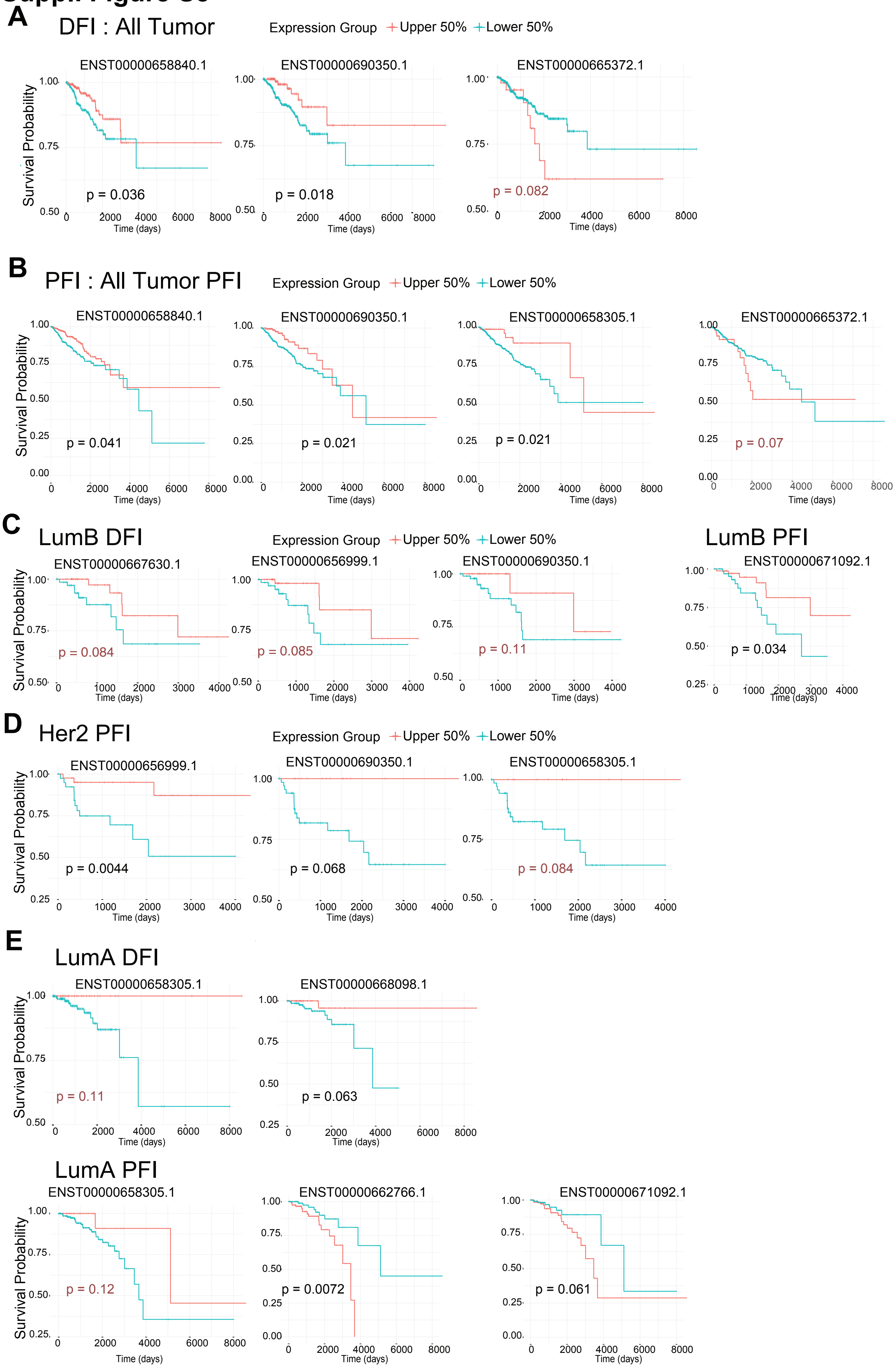

### Suppl. Figure S9

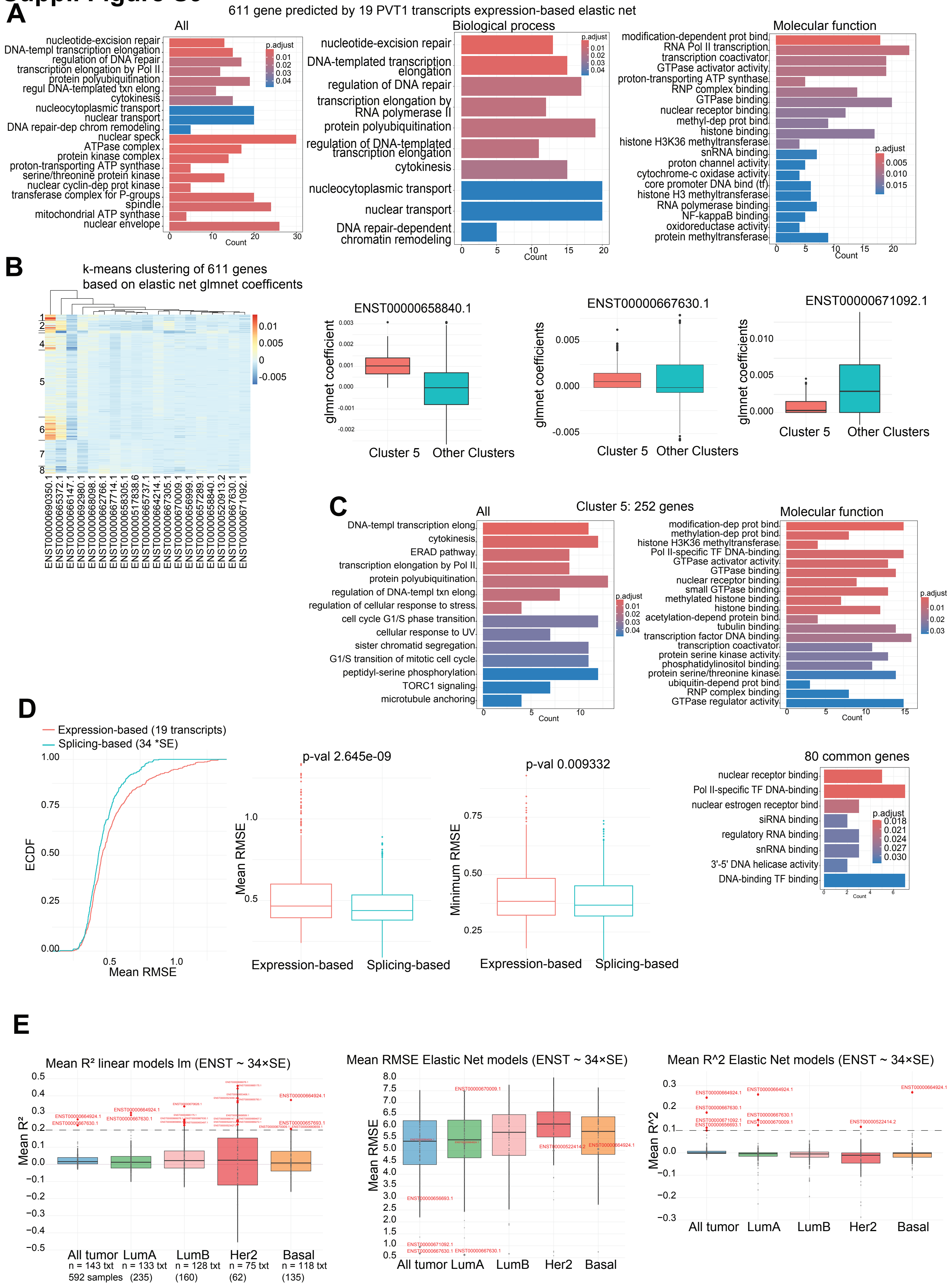

### Suppl. Figure S10 (A-C)

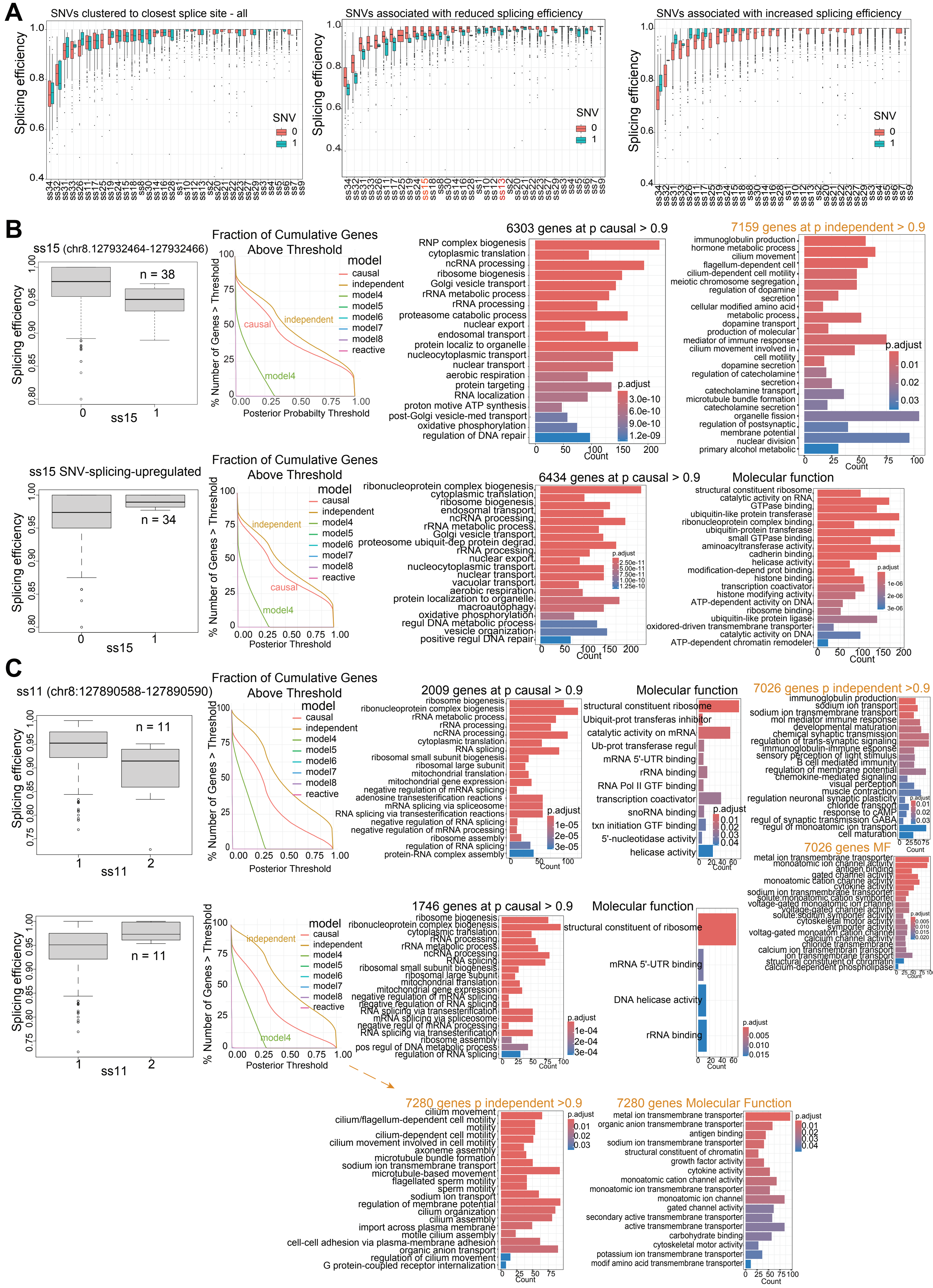

### Suppl. Figure S10 (D-E)

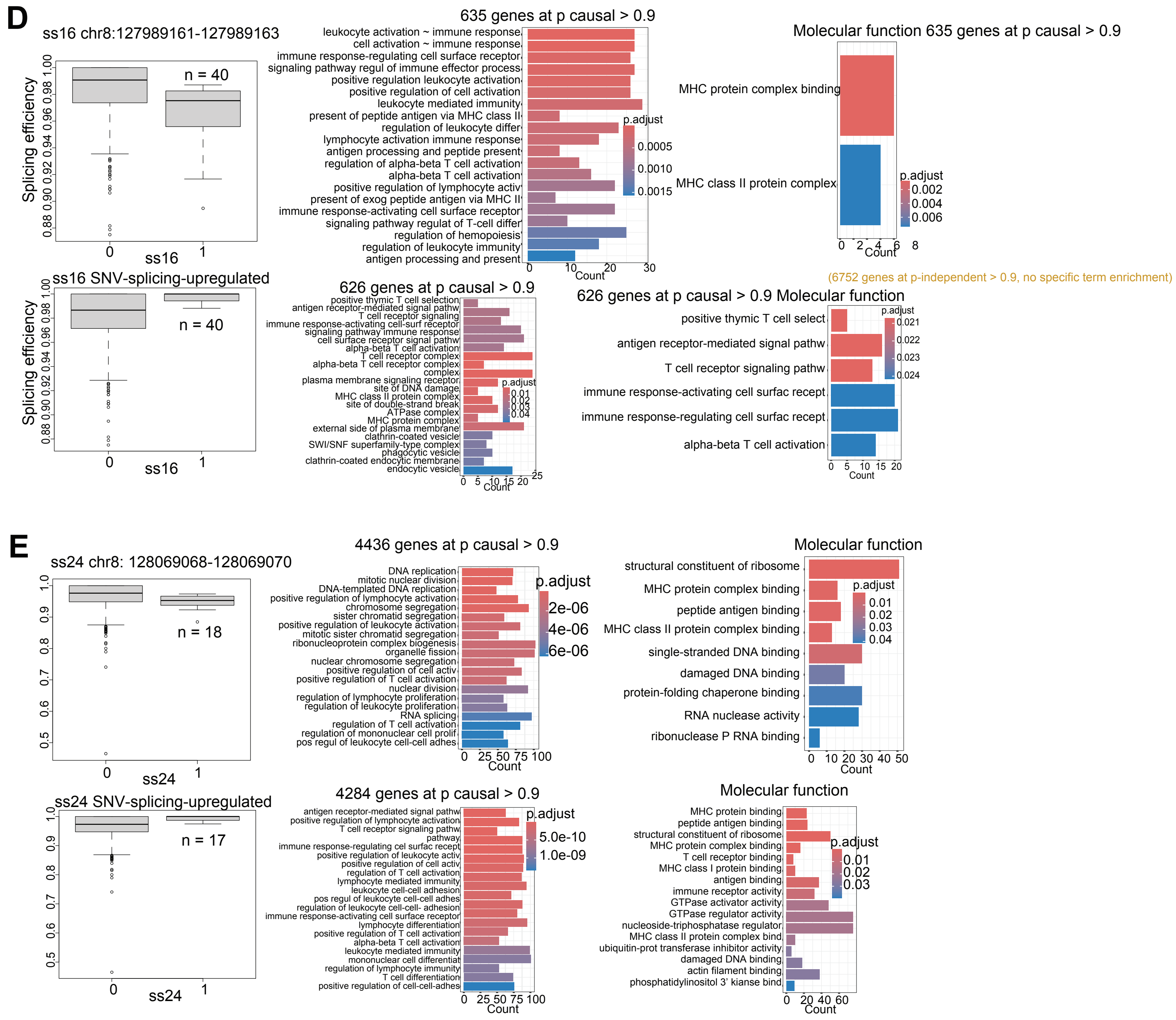

### Suppl. Figure S11

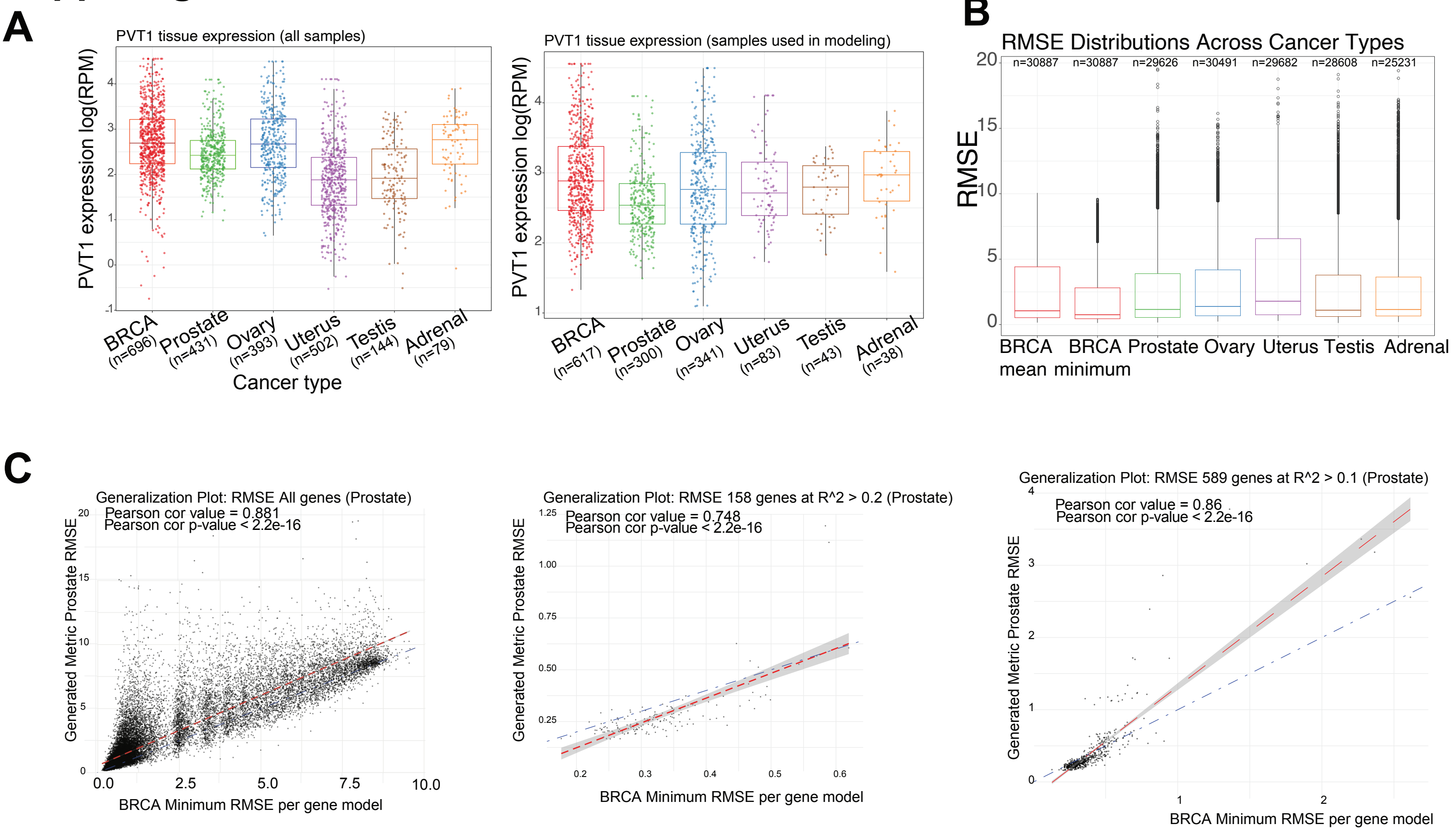

### Suppl. Figure S12

**A**

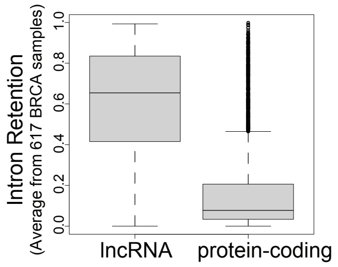


**B**

| Gene Id | Intronic motif count | Total intron length (bp) | Gene name | Type | gene length (bp) | Intron retention | gene RPM | exon RPM |
|---|---|---|---|---|---|---|---|---|
| ENSG00000231607.15 | 13 | 129611 | DLEU2 | lncRNA | 169050 | 0.707 | 11.256 | 3.267 |
| ENSG00000249859.14 | 10 | 382815 | PVT1 | lncRNA | 392575 | 0.576 | 63.691 | 23.45 |
| ENSG00000164756.13 | 12 | 203337 | SLC30A8 | protein_coding | 226441 | 0.534 | 81.469 | 54.78 |
| ENSG00000188001.10 | 14 | 276244 | TPRG1 | protein_coding | 378090 | 0.502 | 78.576 | 50.41 |
| ENSG00000272168.10 | 38 | 966985 | CASC15 | lncRNA | 990271 | 0.496 | 20.603 | 8.834 |
| ENSG00000196090.14 | 27 | 798940 | PTPRT | protein_coding | 1117218 | 0.479 | 125.18 | 54.11 |
| ENSG00000178568.16 | 72 | 1071189 | ERBB4 | protein_coding | 1163124 | 0.399 | 78.63 | 47.53 |
| ENSG00000230590.13 | 20 | 339565 | FTX | lncRNA | 353139 | 0.336 | 43.235 | 27.49 |
| ENSG00000185420.19 | 62 | 580717 | SMYD3 | protein_coding | 757970 | 0.29 | 51.326 | 33.62 |

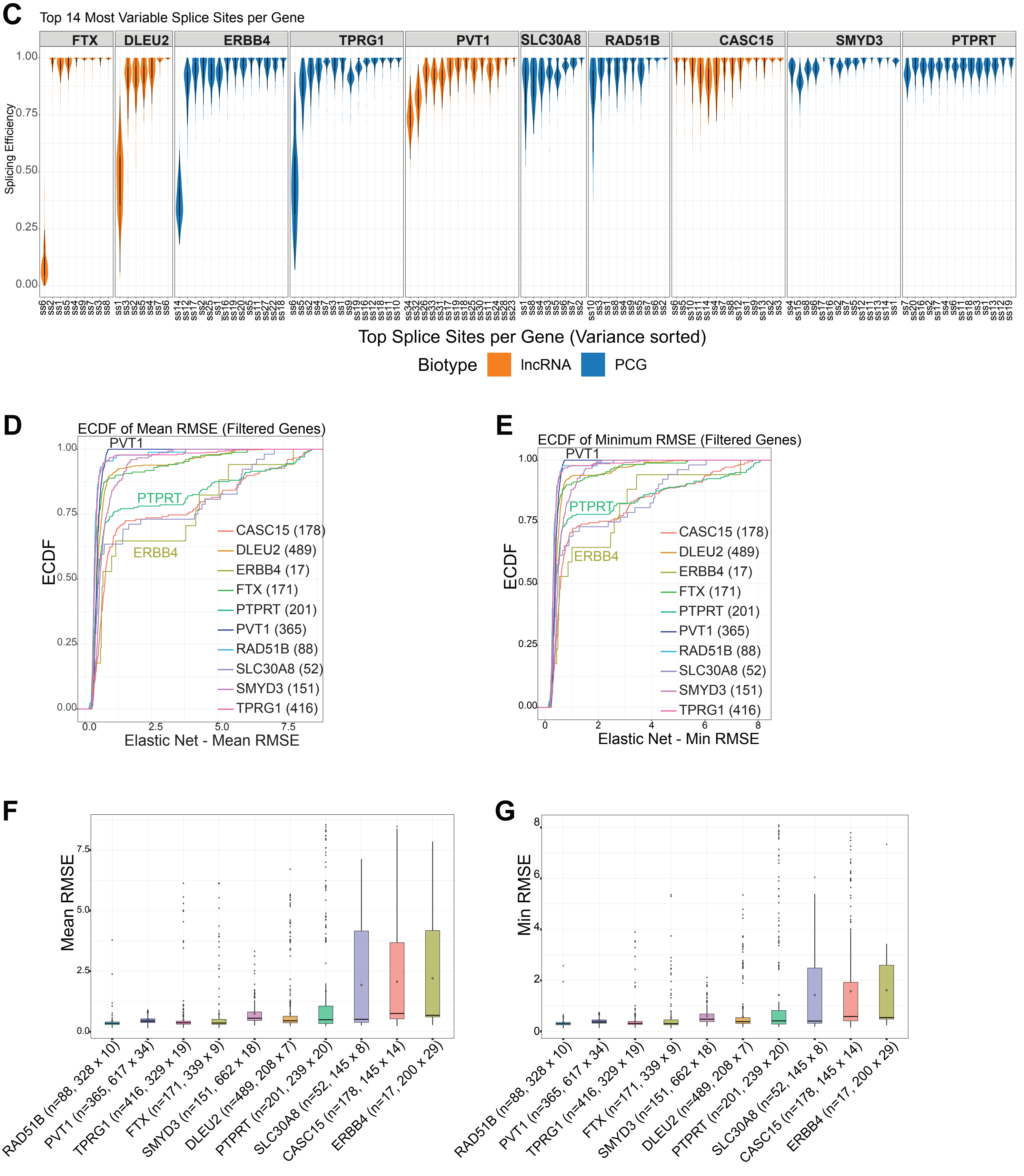

Suppl. Fig. S13

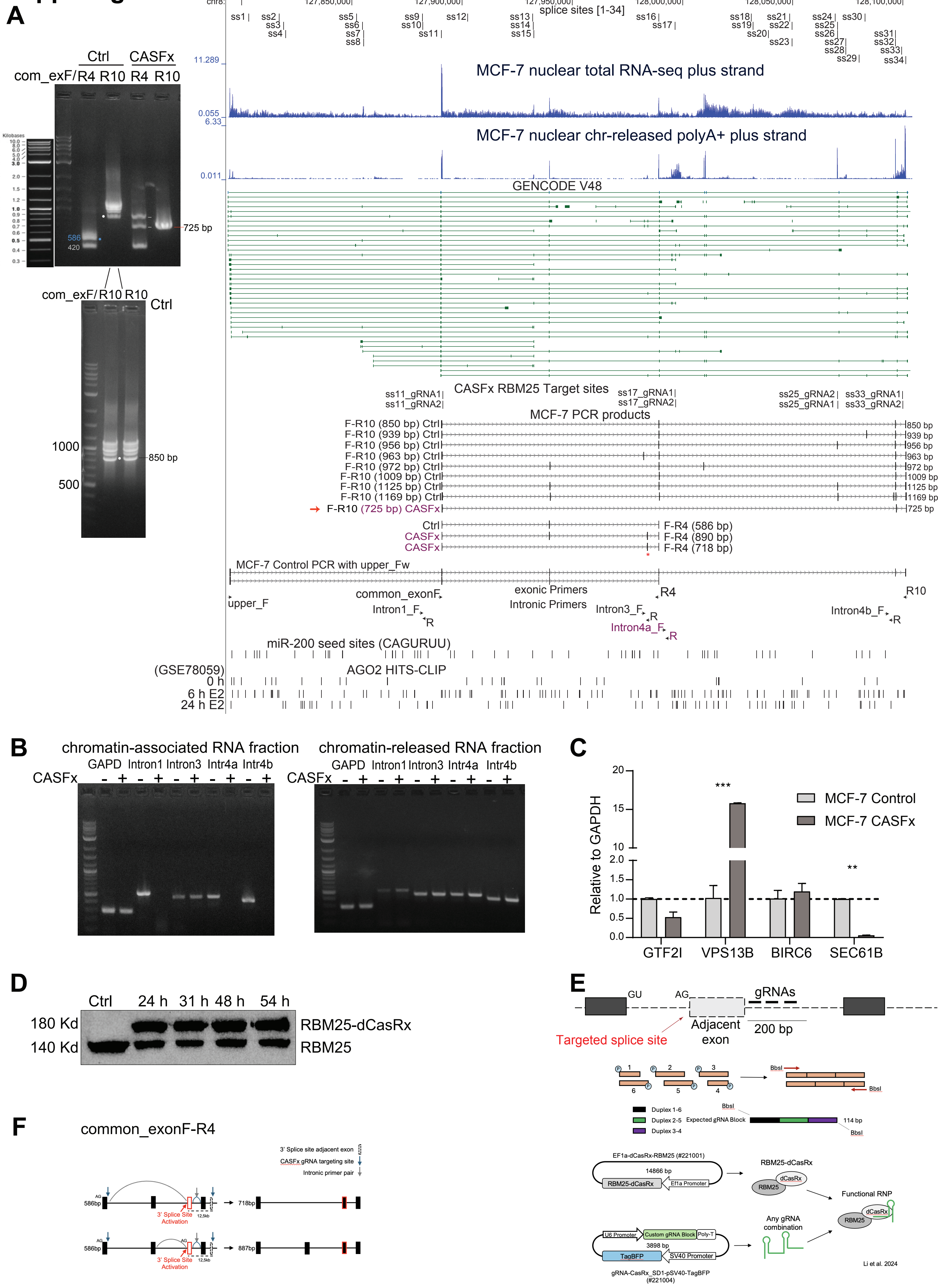

#### Suppl. Figure S14

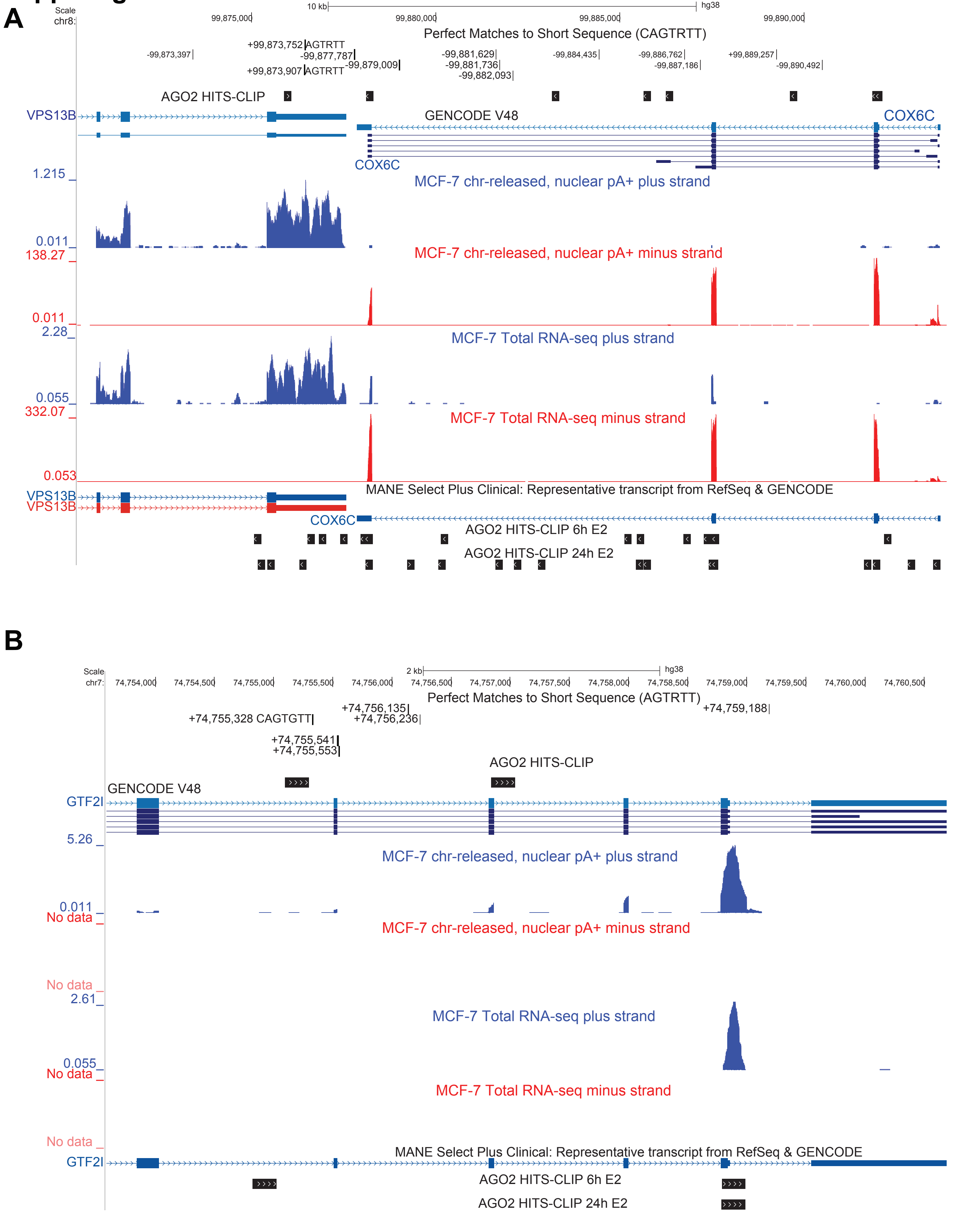

**PVT1 splicing activity predicts genome-wide gene expression with miRNA regulatory signatures**
Kozonakis A. et al.

Supplementary Figure Legends

**Suppl. Figure S1**. (**A**) UCSC screenshot of the MYC-PVT1 locus. Annotated transcripts from GENCODE V48 are shown (dense view), and alternative PVT1 transcript isoforms that were extracted with Salmon across 670 BRCA TCGA RNA-seq data. Only the top 19 transcripts are shown (most variable in expression) from a total of 190 detected (see also Suppl. Fig. S7B-C). Perfect 8-mer miR-200 seed sites (CAGURUUA) are found within PVT1 intronic intervals. Strand-specific (plus strand) bigwig files showing expression in: whole-cell total RNA-seq from MCF-7 and MCF10 (re-processed data from [1]), MCF-7 nuclear chromatin released polyA+ enriched RNA-seq in control and 48 h cisplatin treatment (this study, GEO submitted), and chromatin-associated nascent RNA-seq from different 4-SU pulse-chase time points from [2]. (**B**) Same as in (A) zooming at the PVT1 transcription unit and showing intronic positions of 7-mer miR-200 seed sites (CAGURUU). PVT1 circRNAs are from circBase [3]. (**C**) PVT1 expression (tags per million, TPM) in different tissues, data from GTEx [4]. (**D**) Left: PVT1 expression (reads per million of uniquely mapped reads, RPM) across 53 control and 617 TCGA BRCA samples used in splicing analysis. Middle panel: PVT1 expression in control and tumor samples across 17 different pooled tissue types (shown on the x-label of the histogram plotting number of normal samples from the corresponding healthy tissues). (E) UCSC screenshots showing PVT1 intronic positions of miR-205 seed sites.

**Suppl. Figure S2**. (**A**) UCSC screenshot showing positions of the 34 3' splice sites analyzed in this study. (miR-200 8-mer seed site intronic positions and expression tracks as in S1A). (**B-C**) Boxplot distribution of splicing efficiency values across PVT1 (B) 3' and (C) 5' splice sites in 53 control and 617 TCGA BRCA samples analyzed. (**D-E**) Raw variance in splicing efficiency values of (D) 3' splice sites and (E) 5' splice sites across all BRCA samples analyzed. Red horizontal line in (D) indicates the 12 most variable 3' splice sites that were used in linear regression to predict gene expression (green line in F). (**F**) ECDF plots of performance metric R-squared (R^2) from all 30,887 linear regression models (one model per gene). Linear regression models were trained across all BRCA samples to predict gene expression using as variables either the top 6 most variable 3' splice sites (red line, 6 genes passing significance cutoff R^2 > 0.2), or the top 12 most variable splice sites (green line, 16 genes passing the cutoff), or top 20 splice sites (light blue line, 138 genes passing cutoff), or all 34 3' splice sites (magenta line, 770 genes passing the cutoff). Panels on the right, comparison of lm fit (R^2 values and Pearson correlation between observed and predicted values) for the same gene by training the linear models with different sets of PVT1 3' splice site splicing efficiencies (6, 12, 20, and 34). (**G**) ECDF of linear regression model metrics by training the model with 34 PVT1 3' splice site splicing efficiencies (~34 SE) across all tumor samples in 10 x cross-validation (10 x CV.lm), one model per gene (30,887 genes were screened). The average adjusted R-squared and mean correlation (between observed and predicted values) from all 10 folds are plotted. Right panel: ECDF of adjusted R-squared values (average from 10-fold cross-validation) of predicted genes that satisfy the criterion of a mean correlation (between observed and predicted values) from all 10 folds > 0.2. 316 genes were considered as significant PVT1 splicing-based linear regression model predictions. (**H**) Examples of linear fit for 3 genes (out of the 316) predicted by the PVT splicing-based (~34 SE) linear regression models trained in 10 x cross-validation across all tumor samples. (**I**) Gene ontology (GO) terms enrichment analysis of the 316 genes predicted by the PVT1 splicing-based (~34 SE) linear regression model using clusterProfiler [5]. (**J**) ECDF linear regression R-squared metric from 30,887 genes predicted either by the pVT1 ~34 SE splicing-based (~34 SE) model or PVT1 expression (exon-level counts). On the right, PVT1 gene expression prediction linear fir (observed vs. predicted values) by its own splicing-based (~34SE) linear regression model, and MYC gene expression prediction by the PVT1 splicing-based model; neither PVT1 nor MYC fits pass the cutoff or R-squared > 0.2. Lower panel, boxplot distribution of PVT1 expression-based (exon-level) and splicing-based (~34 SE) linear regression metric R-squared values from all 30,887 genes models.

**Suppl. Figure S3**. Correlation heatmaps (**A**) Correlation heatmap of splicing efficiency values at all PVT1 splicing efficiency values (across all BRCA 670 samples) and Intron retention at the locus; intron retention was extracted as a proxy: (gene-level raw counts – exon-level raw counts) / gene-level raw counts. (**B**) Heatmap showing correlation among top 19 PVT1 alternative transcript isoforms quantified with Salmon [6] across all 670 BRCA samples. Only ENST00000667630.1 and ENST00000671092.1 (underlined) show positive correlation with overall intron retention at the locus. (**C**) Composite correlation heatmap PVT1 splicing efficiencies at 34 3' splice sites and Salmon quantified top 19 alternative transcript isoforms.

**Suppl. Figure S4**. Clustering of PVT1 splicing-based (~34 SE) elastic net-predicted genes based on glmnet coefficients. (A-B) Same as Main Figure 3D-E. (**C**) K-means clustering of 620 filtered genes predicted by PVT1 splicing-based (~34 SE) elastic net (passing significance cutoffs) trained in 10 x cross validation across 235 BRCA samples of LumA pam50 subtype. On the right, GO terms enrichment plotting with clusterProfiler (terms enriched skewed toward translation and ribosomal components). miR-200 target genes are significantly enriched in clusters 3 and 6 (Fisher's exact test p-value 1.142e-12, odds ratio 7.024) which are defined by positive contribution of ss5, ss9, ss10, ss13, ss14, ss15. (**D**) K-means clustering of 158 filtered genes (passing significance cutoffs) predicted by PVT1 splicing-based (~34 SE) elastic net model trained in 10 x cross validation across 178 BRCA samples of LumB pam50 subtype. miR-200 target genes are enriched in clusters 1 and 3 which are defined by positive contribution (positive glmnet coefficients) of splice sites ss13, ss14, ss15 (Fisher's exact test p-value 0.0137, odds ratio 3.055).

**Suppl. Figure S5**. PVT1 splicing activity distinguishes BRCA subtypes. (A) Boxplot distribution of PVT1 splicing efficiency values at 34 3' splice sites across BRCA tumor subtypes. (B-C) Supplementary to main Figure 4 including also the Random Forest Mean Gini plots for each binary classification.

**Suppl. Figure S6. (A-E)** Kaplan-Meier survival analysis plots for BRCA patients split based on PVT1 3' splice site spicing efficiency. Probing survival probabilities (disease-free interval, DFI; progression-free interval, PFI) was performed for all 34 3' splice sites across all tumor samples, and within pam50 subtypes; here we show splice site results at p -value < 0.05 and some splice site that showed a non-significant trend (clear line separation). Samples were split based on splicing efficiency median value. (**F**) DFI and PFI survival probabilities for all pam50 subtypes. P-values indicate log-rank test results (from *survminer* package [7, 8]) of binary subtype comparisons.

**Suppl. Figure S7.** Evaluation of PVT1 alternative transcript isoform expression-based classification models. (**A**) Boxplot distribution of Random Forest classification accuracy values from 10-fold cross-validation distinguishing a pam50 subtype from the rest of samples; Random Forest were trained in 10xcross-validation using Salmon-quantified expression (tags per million, TPM values) from 19 PVT1 alternative transcript isoforms. (**B**) Boxplot distribution of Salmon-quantified expression values (TPM) for 19 identified PVT1 transcripts quantified across BRCA subtypes. (**C**) Raw variance (top) and median of expression (bottom plot) for top sorted PVT1 alternative transcript isoforms quantified with Salmon [6] across all BRCA tumor samples. The top 19 most variable transcripts were used in downstream analysis (k-means clustering, survival analysis, genome-wide gene expression prediction with elastic nets). (D-F) K-means clustering for binary classification of BRCA subtypes (specific subtype versus the rest) using the 19 most variable Salmon-quantified PVT1 transcripts as classification features in 10 x cross-validation (**D**) Her2 vs. non-Her2 (**E**) Basal vs. non-Basal and (F) LumB vs. non-LumB.

**Suppl. Figure S8**. Survival Analysis (DFI, Disease free interval probability; PFI, progression free interval probability) was performed across all tumor samples (**A-**B), and within subtypes: (C) LumB, (D) Her2, (E) LumA (and Basal, no significant results), by splitting samples based on the median of PVT1 alternative transcript expression. All top 19 most variable PVT1 Salmon-quantified transcripts were assayed, here we show significant results (at $p < 0.05$), and some non-significant (due to sample size) with clear line separation.

**Suppl. Figure S9.** Evaluation of PVT1 alternative transcript isoform expression-based elastic net models in predicting genome-wide gene expression. (**A**) GO terms enrichment analysis using clusterProfiler [5] of 611 genes (out of 30,887) predicted above certain significance cutoffs (Methods) by elastic net models trained in 10 x cross-validation across all BRCA tumor samples using the Salmon-quantified expression values of the top 19 most variable PVT1 transcripts as variables. (**B**) K-means clustering of the 611 high-confidence genes predicted by the pVT1 transcript expression-based elastic nets used the model's glmnet coefficients as clustering parameters. miR-200 target genes are significantly enriched (odds ratio 4, Fisher's exact test p-value = 4.42e-12) only in one cluster, cluster 5, which is defined by positive contribution of ENST00000667630.1, ENST00000671092.1 and ENST00000658840.1. Notably ENST00000667630.1, ENST00000671092.1 (and to a lesser extent ENST00000658840.1) are the only transcripts that show positive correlation with the overall intron retention at the PVT1 locus (Suppl. Figure S3B), suggesting the mir-200 target gene regulation may be indirect through intron-retention mechanisms related to the presence of miR-200 seed sites (perfect 7-mer matches) located exclusively within introns at the pVT1 locus (Suppl. Fig. S2A). (**C**) GO terms enriched analysis of the 252 genes of cluster 5 using clusterProfiler [5]. (**D**) Comparison of PVT1 splicing-based (splicing efficiency values at 34 3' splice sites, ~34 SE) and alternative transcript isoform expression based (~19 ENST) elastic net performance in predicting genome-wide gene expression; metrics from all 30,887 gene models are plotted: ECDF of mean RMSE from 10 x cross-validation (left panel); comparison in boxplot distribution of splicing-based (~34 SE) and transcript expression-based (~19 ENST) elastic net model mean (from 10 x cross-validation) RMSE (middle panel); and minimum RMSE (right panel). (**E**) PVT1 splicing efficiency at 34 3' splice sites (~34 SE) splicing-based models perform poorly, either in linear regression (left panel) or in elastic models (middle and right panels), in predicting expression of PVT1 alternative transcript isoforms (all 190 Salmon-quantified transcripts were assayed), with low mean R-squared in linear regression models (left panel, marked red transcripts that passed the cutoff of $R^2 > 0.2$ (n denotes number of predicted transcripts, out of all 190 assayed, that could be modeled using cross validation elastic net). Middle and left panels: boxplot distributions of mean and minimum RMSE from cross-validation folds of splicing-based elastic nets trained in 10 x cross-validation (5 x cross-validation within subtypes) to predict PVT1 alternative transcript isoform expression. All models were trained either across all tumor samples (10 x cross validation) or within each subtype (5 x cross validation; number in parenthesis denotes number of samples).

**Suppl. Figure S10.** Causal Inference results, supplementary figure to main Figure 5. n = 592 samples in total, with an assayed splice site either assigned an SNV ('1'), or not ('0').

**Suppl. Figure S11**. Cross-tissue generalization of PVT1-splicing based gene expression elastic net models. Splicing-based elastic net models were trained across all BRCA tumor samples using PVT1 splicing efficiency at 34 3' splice sites (~34 SE) as variables in 10 x cross validation to predict gene expression genome-wide (30,887 gene models). The coefficients from the best fold (the fold with minimum RMSE in BRCA) were used in cross-tissue assay of elastic net model applicability. (**A**) Comparison of PVT1 expression across cancer types samples, all (left panel) and used (right panel). (**B**) Boxplot distribution of RMSE across all assayed gene models which were trained in BRCA in 10 x cross-validation (10-fold mean and minimum RMSE are shown) and applied in unseen data of five other tumor types (n denoted number of gene models; see also main Figure 6A). (**C**) Cross-tissue generalization of BRCA-trained PVT1 splicing-based models in the prostate (PRAD), plotting the RMSE per gene calculated in the prostate (y axis) to the BRCA-trained counterpart (minimum RMSE from best fold on x axis) for all gene models (left panel), genes passing R-squared > 0.2 (middle panel) and R-squared > 0.1 (right panel).

**Suppl. Figure S12.** PVT1 as a top candidate acting as a chromatin-associated lncRNA with intronic-retained ceRNA potential. (**A**) Comparison of overall intron retention among lncRNA and protein-coding genes. Intron-retention was calculated as a proxy as normalized RNA-seq read coverage (gene_level - exon_level)/gene_level read counts. The average intron retention value across 617 BRCA tumor samples is plotted. (**B**) We performed a genome-wide in silico screen to identify genes that contain strand-specific intronic miR-200 seed sites, perfect matches to the 8-mer CAGTRTTA, and provide a list in Suppl. Table S3. Gene candidates were sorted based on intron retention, gene expression, intronic motif count. Candidates assayed in splicing-based machine learning modeling to predict gene expression genome wide are shown. (**C**) Elastic net models were trained in cross-validation across BRCA tumor samples to predict gene expression genome-wide (same workflow as for the PVT1 splicing-based gene expression models in genome-wide screens, Methods). Comparison of splicing efficiency values at the 14 most variable splice sites across all gene candidates assayed in splicing-based predictive models (Methods). (**D-G**) Comparison of elastic net metric mean and minimum RMSE per splicing-based model for filtered predicted genes. Per splicing-based model of each candidate, (PVT1 splicing-based model included), predicted genes were filtered passing the same specific significance cutoffs as in the PVT1 workflow (Methods). In the x-label of (F-G), in the parenthesis 'n' = number of significant genes predicted by a given candidate splicing-based model, alongside 'number of samples that the model was trained' x 'number of 3' splice sites used as variables'.

**Suppl. Figure S13**. Experimental validation of PVT1 intron-retained ceRNA potential with CASFx in *vitro*; figure supplementary to main Figure 7.

**Suppl. Figure S14**. UCSC screenshots of two candidate miR-200 target genes with altered expression upon CASFx artificial splicing enhancement of PVT1 in vitro; figure supplementary to main Figure 7.

References (cited in Supplementary Figure Legends)

7. Therneau, T.M. A Package for Survival Analysis in R. 2024.

8. Terry M. Therneau, Patricia M. Grambsch. Modeling Survival Data: Extending the Cox Model. 2000.
