## Supplementary Methods for "PVT1 splicing activity predicts genome-wide gene expression with miRNA regulatory signatures"

### Quantitative trait loci

We built our cohort using all the available TCGA normal tissue samples, for a total of 713 across 23 different projects. We downloaded the corresponding germline mutation files for all 713 samples using the following filters: i) Experimental Strategy: Genotyping Array ii) Data Category: simple nucleotide variation iii) Data Type: Simple Germline Variation iv) Workflow Type: Birdseed v) Platform: affymetrix snp 6.0 and included both solid tissue (n=363) and peripheral blood nos (n=350). Splicing efficiency at the 34 PVT1 3' splices sites and genome-wide exon-level gene expression was computed in the same manner as BRCA (see Methods: Splicing efficiency values at PVT1 3′ splice sites; Quantifying genome-wide gene expression). After extracting splicing efficiency, our cohort was reduced to 234 samples where BRCA, HNSC and PRAD accounted for 152 of the samples (Suppl. Fig. S1D). Genes with low expression (raw counts less than 10 in less than 10 samples) were kicked out, resulting in a total of 38,700 genes. Affymetrix genotype data was filtered (Genome-wide human SNP 6.0 array GPL6801 after remapping to hg38) to retain only mutations in the PVT1 locus (n=220 unique SNPs). For the 234 samples, we processed the corresponding germline mutation files (birdseed) using the Affymetrix array, while also keeping SNP calls with a confidence < 0.05. Each sample was defined with "0" for homozygous (no mutation), "1" for heterozygous, "2" for homozygous alternate and "-1" if missing and then genotyped (tabix -p vcf).

The splicing efficiency values and the genotyped VCF were used as input to *sQTLseekeR2* [1] to find potential splicing quantitative trait loci (cis-sQTL); only 1 was discovered with a significant p-value < 0.05. As covariates, we tried including gene level expression of PVT1 (RPM) and/or tissue type information, with no changes in significance.

For discovery of trans-acting expression quantitative trait loci (trans-eQTL) we used *QTLtools* [2]. We created a VCF file from the Affymetrix array (0/0, 0/1, 1/1 or ./. for homozygous, heterozygous, homozygous alternate and missing respectively) and a BED file with RPM exon level gene counts (n=38,700 genes).

*QTLtools trans --seed 12345 --permute 1000 --normal –vcf SNP.vcf.gz –bed expression.bed.gz*

The analysis revealed 185 significant trans-eQTLs (p-val < 1e-6), which dropped to 72 and 69 when tissue type and/or gene expression PCA were included as covariate files (using QTLtools pca; *QTLtools pca --bed expression.bed.gz --center --scale*).


1. Garrido-Martín, D., B. Borsari, M. Calvo, et al. Identification and analysis of splicing quantitative trait loci across multiple tissues in the human genome. *Nat Commun* 2021;**12**:727.

2. Delaneau, O., H. Ongen, A.A. Brown, et al. A complete tool set for molecular QTL discovery and analysis. *Nat Commun* 2017;**8**:15452.
